# Multiscale Modes of Functional Brain Connectivity

**DOI:** 10.1101/2024.05.28.596120

**Authors:** S Rezvan Farahibozorg, Samuel J Harrison, Janine D Bijsterbosch, Mark W Woolrich, Stephen M Smith

**Affiliations:** FMRIB, Oxford Centre for Integrative Neuroimaging, Nuffield Dept. of Clinical Neuroscience, Oxford University, Oxford, UK; Department of Radiology, Washington University in St Louis, St. Louis, USA; OHBA, Oxford Centre for Integrative Neuroimaging, Department of Psychiatry, Oxford University, Oxford, UK

## Abstract

Information processing in the brain spans from localised sensorimotor processes to higher-level cognition that integrates across multiple regions. Interactions between and within these subsystems enable multiscale information processing. Despite this multiscale characteristic, functional brain connectivity is often either estimated based on 10-30 distributed modes or parcellations with 100-1000 localised parcels, both missing *across*-scale functional interactions. We present Multiscale Probabilistic Functional Modes (mPFMs), a new mapping which comprises modes over various scales of granularity, thus enabling direct estimation of functional connectivity within- and across-scales. Crucially, mPFMs were not formulated, but emerged from data-driven multilevel Bayesian modelling of large functional MRI (fMRI) populations and every individual. We demonstrate that mPFMs capture both distributed brain modes and their co-existing subcomponents. In addition to validating mPFMs using simulations and real data, we show that mPFMs can predict ∼900 personalised traits from UK Biobank more accurately than current standard techniques. Therefore, mPFMs can offer a new basis for functional connectivity modelling and yield enhanced fMRI biomarkers for traits and diseases.

## 1 Introduction

Within the complex system of 86 billion neurons (Herculano-Houzel, 2009) in the human brain, large ensembles of neurons work in synchrony, such that they produce distinct modes of extended, correlated activity (Smith et al., 2009). These functional modes exhibit a spectrum of scales and functionalities, from localised brain modes facilitating specialised information processing, such as responses to sensory or motor stimuli, to higher-level cognitive modes integrating across multiple regions through long-range connections. Functional modes underlie spontaneous brain activity during task-free periods (often referred to as resting state networks, RSNs), adapt dynamically during cognitive tasks, and undergo permanent changes in neurodegenerative diseases (Biswal et al., 1995; Buckner and Vincent, 2007; Calhoun et al., 2008; Damoiseaux et al., 2006; Raichle et al., 2001; Smith et al., 2013b). Growing evidence, especially from new datasets with thousands of individuals, has highlighted that spatiotemporal characteristics of functional modes vary systematically across individuals, providing biomarkers for traits and disease, akin to a diagnostic assay for the brain (Elliott et al., 2018; Finn et al., 2015; Jiang et al., 2020; Kong et al., 2019; Smith et al., 2015; Vidaurre et al., 2017). In this paper, we introduce a novel representation, Multiscale Probabilistic Functional Modes (mPFMs), which, unlike the existing single-scale modes, encapsulates an ensemble of modes across multiple scales. mPFMs have the potential to provide a shift in how functional connectivity has been modelled to date, by offering a new basis to gain insights into spatiotemporal connectivity within and across diverse processing scales, and providing more sensitive biomarkers from functional neuroimaging.

Functional MRI (fMRI), with a spatial resolution of ∼2mm and a temporal resolution of ∼1s, provides a suitable spatiotemporal resolution to characterise functional modes non-invasively. This has largely been done using two distinct approaches: a) high-dimensional (highD) decomposition of fMRI data into functional parcellations with 100-1000 modes (i.e., parcels), where each mode is typically *localised* to a single brain region (Bellec et al., 2010; Craddock et al., 2012; Schaefer et al., 2018); b) low dimensional (lowD) decomposition of fMRI data into ∼20-30 modes (Beckmann and Smith, 2004; Calhoun et al., 2001) - the conventional RSNs - each *distributed* across multiple distant brain regions, and often with some spatial overlap between the modes. The latter is often carried out using spatial Independent Component Analysis (ICA) (McKeown et al., 1998), or lowD parcellations (Yeo et al., 2011). Each of these two approaches offer distinct benefits. On the one hand, fine-grained, highD parcellations capture local details in the brain’s organisation, especially details that have proved useful for cross-individual variability modelling, and prediction of traits and disease (Dadi et al., 2020; Pervaiz et al., 2020). On the other hand, lowD large-scale modes capture the more global functional configuration of the brain. Distant brain regions with intrinsic long-range connections are grouped into biologically meaningful neural entities such as language and attention networks, allowing for easier interpretation and putative links to cognitive functions (Power et al., 2011; Smith et al., 2009; Yeo et al., 2011). Additionally, lower data dimensionality makes subsequent statistical analyses more manageable. Therefore, distributed vs localised modes each offer distinct benefits and limitations for capturing global vs local functional architecture of the brain. Crucially, however, neither of these approaches directly model inherent interactions across multiple scales of information processing in the brain.

In this context, we report the finding of mPFMs, which can bridge this gap by providing a unified representation that encompasses modes across multiple scales. mPFMs were identified through high dimensional decomposition of resting state fMRI using the sPROFUMO framework (Farahibozorg et al., 2021). sPROFUMO defines a Bayesian hierarchy with two levels of group and individuals across big data populations, where group priors are iteratively used for top-down regularisation of individuals, and evidence across individuals is accumulated and fed back to the group. Two key distinguishing aspects of the model are that: 1) sPROFUMO is subject-specific, i.e., inferring the modes for populations and individuals *simultaneously*; 2) unlike ICA or parcellation-based techniques, sPROFUMO does not impose strong constraints on spatial or temporal independence between the modes, thus yielding modes that are more flexibly interacting, spatially and temporally (Farahibozorg et al., 2021; Harrison et al., 2020).

We report that this added flexibility in modelling population variations, and in spatiotemporal connectivity between the modes, resulted in the discovery of mPFMs (Figure 1), which preserve conventional large-scale modes, and add new modes with different scales of granularity to the ensemble. Figure 1 shows direct comparisons between lowD vs highD Probabilistic Functional Modes (PFM) decomposition of rfMRI data in HCP, where a force-directed network layout is used for visualisation (Bastian et al., 2009). LowD decomposition (shown in yellow) yielded the conventional distributed-only resting state networks, hereafter referred to as large-scale RSNs. The highD decomposition yielded a mixture of modes across various scales. By spatial pairing of the modes from highD and lowD decompositions, we found a one-to-one correspondence between a set of highD modes (shown in orange) and large-scale RSNs, indicating that as we increase the dimensionality from 25 to 150, conventional RSNs are preserved, and new modes are added (shown in blue). This is a novel type of decomposition, and since it includes functional modes across multiple scales, we refer to it as Multiscale Probabilistic Functional Modes, mPFMs.

**Figure 1.**
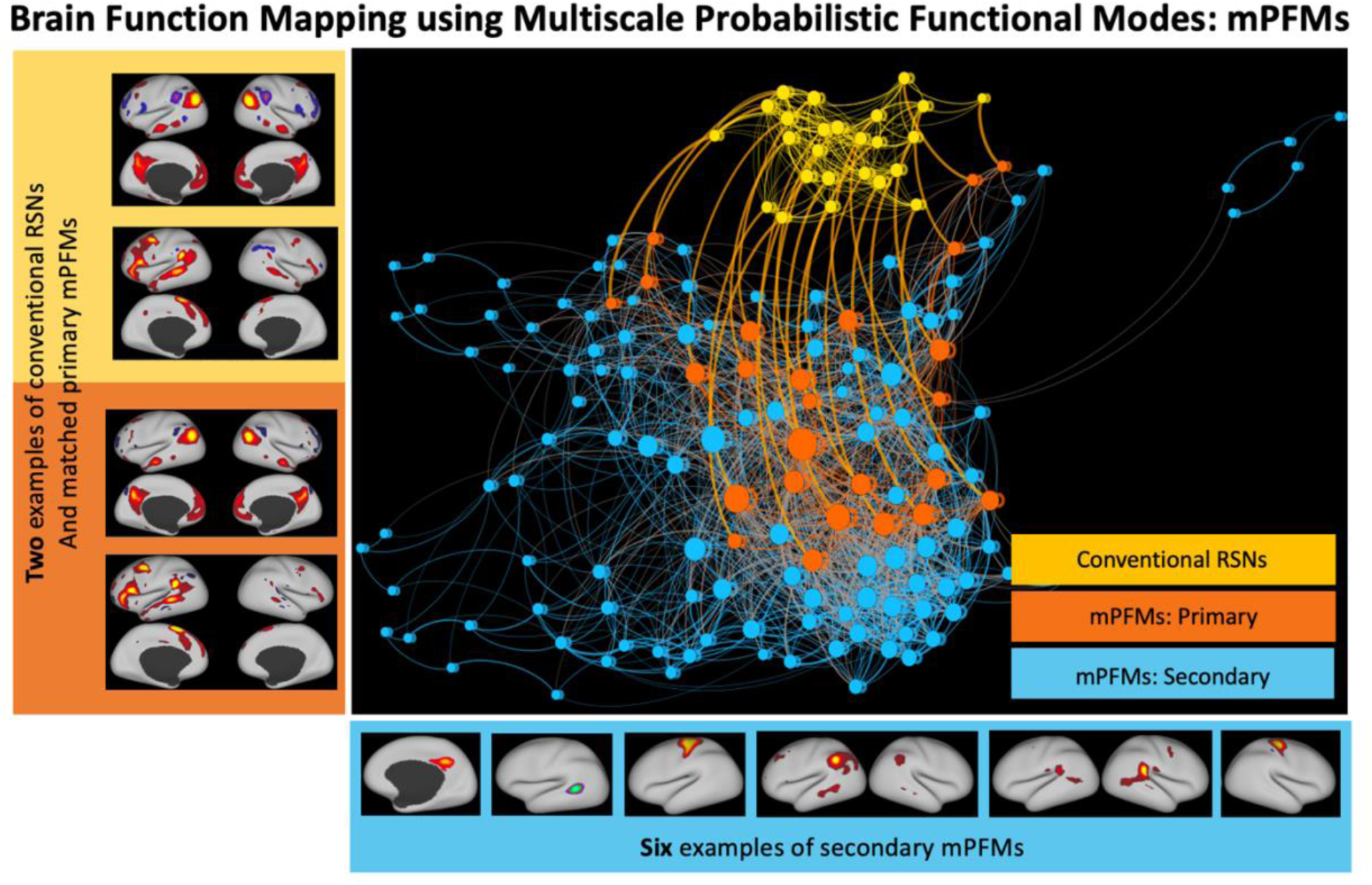
Brain function mapping using Multiscale Probabilistic Functional Modes (mPFMs) High-dimensional (150) Probabilistic Functional Modes estimated from resting state fMRI (1003 HCP subjects) yield a new representation of the brain’s functional organisation which consists of an ensemble of conventional distributed-only (i.e., large-scale) resting state networks (RSNs) and newly-added modes across more fine-grained scales. We refer to this new representation as Multiscale Probabilistic Functional Modes, mPFMs. When paired with large-scale RSNs from 25-mode decomposition (yellow), 25 of the mPFMs showed a clear matching, labelled as Primary mPFMs (orange). Secondary mPFMs (blue) are the remaining 125 mPFMs, that start appearing with increased dimensionality. A force-directed layout is used to visualise the spatial similarities (i.e. correlations) between the modes. In this layout, each circle denotes a functional mode, which is a node in the connectivity matrix, and each line is the connection between a pair of modes. The size of each mode shows the average strength of its connections, line widths denote the strength of spatial correlations (i.e. overlaps) between two modes, and the distance between the modes denotes how close two modes are in the connectivity matrix, both within and between low-dimensional vs high-dimensional decompositions. All connections within-large-scale and within-multiscale, as well as paired connections between large-scale and multiscale were maintained in visualisation. Matrix-based visualisation of these results is shown in Figure S 2.

mPFMs can be considered an alternative to conventional (“single scale”) resting state networks (RSNs). We report that: a) mPFMs emerge to explain the existence of multiple distinct time-courses within each large-scale mode, thus capturing voxel-to-voxel variability of temporal dynamics that cannot be captured by these distributed modes alone; b) we validate mPFMs using several methods, including synthetic data, reproducibility between Human Connectome Project (HCP) and UK Biobank (UKB) data, and link to HCP’s Multimodal Parcellation (Glasser et al., 2016); c) we further show the added benefit of cross-scale interactions of mPFMs, compared with within-scale interactions, for capturing cross-individual variability and predicting personalised traits; d) finally, we show that, compared with two commonly used techniques, spatial ICA modes and parcellations from the Schaefer Atlas (Schaefer et al., 2018), mPFMs (of matched high dimensionality) yield more accurate prediction of a wide range of imaging- and non-imaging-derived phenotypes (IDPs/nIDPs) related to cognition, cardiovascular health, brain anatomy, and task fMRI in UK Biobank.

## 2 Materials and Methods

**Figure S 1** shows a flowchart of different subsections of materials and methods, and their link to the corresponding results subsections.

### 2.1 Data

#### 2.1.1 Human Connectome Project (HCP)

We used minimally pre-processed fMRI data from Human Connectome Project (HCP) from S1200 data release (https://www.humanconnectome.org/study/hcp-young-adult), with acquisition and processing details outlined in (Glasser et al., 2013; Smith et al., 2013a). Resting state data from 1003 subjects aged 22-35 years were used here and included 4×15 minute runs per subject (i.e., 1 hour recording per subject). With repetition time (TR) of 0.72s, this dataset includes 4800 time points per individual. Study protocol of the HCP study was approved by the WU-Minn HCP Consortium’s institutional review boards, and written informed consent was obtained from all participants prior to data collection. We used two data formats, volumetric and HCP CIFTI. CIFTI data includes grey matter voxels only (i.e., greyordinates), where cortical grey matter is surface-registered using MSMAll (Robinson et al., 2014) and subcortical grey matter is in volumetric space. This results in inherent data differences such as signal to noise ratio between cortical and subcortical regions, affecting matrix factorisation techniques in, e.g., finding an unbalanced number of cortical vs subcortical modes. To prevent such unbalances and make the results more easily comparable across CIFTI and Volumetric fMRI, we only used cortical greyordinates of CIFTI in this paper. ICA-FIX was used to remove artefacts from volumetric data before resampling to 32k_fs_LR space and projecting onto the surface. Standard HCP preprocessing includes spatial smoothing with a 2mm FWHM smoothing kernel. We applied additional smoothing to obtain 5mm FWHM (for both volumetric and CIFTI data), which improves the Signal to Noise Ratio (SNR) and was found to be useful for obtaining reliable subject-specific modes (Farahibozorg et al., 2021).

HCP data were predominantly used for analyses that aimed at examining timecourse and functional connectivity modelling using mPFMs, their reproducibility across datasets (HCP and UK Biobank), and validation of mPFMs against existing functional parcellations. Given the longer recordings per subject (1 hour vs 6 minutes), we deemed HCP more suitable for these analyses.

#### 2.1.2 UK Biobank (UKB)

We randomly selected 4999 subjects aged > 45 years from the May 2019 release of UK Biobank data (application number 8107). The UKB has approval from the North West Multi-centre Research Ethics Committee (MREC) to obtain and disseminate data and samples from the participants (http://www.ukbiobank.ac.uk/ethics/), and these ethical regulations cover the work in our study. Written informed consent was obtained from all participants prior to data collection. Resting state fMRI data in UKB (at the time that this research was conducted) was in standard volumetric space, and included 1 recording of ∼6 minute per subject. With a TR of 0.735s, data consisted of 490 time points per session. The standard UKB pipeline, which includes quality control, brain extraction, motion correction, artefact rejection using FSL-FIX, high-pass temporal filtering (sigma = 50.0s, Gaussian-weighted least-squares straight line fitting) and registration to standard MNI-2mm space was used for preprocessing (Alfaro-Almagro et al., 2018). Standard UKB preprocessing includes spatial smoothing with a 2mm FWHM smoothing kernel. We applied additional smoothing to obtain 5mm FWHM, which improves SNR and is useful for obtaining reliable subject-specific high dimensional PFMs. UKB was used for evaluating reproducibility of mPFMs (i.e. high-dimensional PFMs) across datasets, and their prediction accuracy for personalised traits. Given the wide range of imaging derived phenotypes (IDPs) and non-imaging derived phenotypes (nIDPs) in UK Biobank, this dataset was deemed more suitable for prediction of traits (see section 2.6 for details).

### 2.2 Estimating Resting State Networks

#### 2.2.1 Probabilistic Functional Modes (PFMs)

Details of PROFUMO/sPROFUMO models and inference are elaborated in our previous work (Farahibozorg et al., 2021; Harrison et al., 2020). Here we provide a brief summary of the model and its application to the data. sPROFUMO is a matrix factorisation model for big fMRI data, which uses hierarchical Bayes with two levels of modelling and inference: population and individual.

At the subject level, fMRI timeseries (**D**^sr^) are decomposed into a set of spatial maps (**P**^s^), time courses (**A**^sr^) and time course amplitudes (**H**^sr^), with residuals ***̅***^*sr*^ :

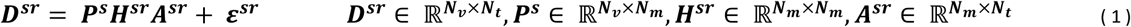

where s denotes subject, r denotes a recording session and **H**^sr^ denotes mode amplitudes. N_v_, N_m_ and N_t_ denote the number of voxels, modes and time points, respectively.

**P**^s^ denotes spatial mode layout across brain voxels, hereafter referred to as *spatial maps*, or just *maps*. These are modelled using a Double-Gaussian Mixture Model (DGMM), with one Gaussian component used to model signal and a second Gaussian distribution to model the background spatial noise in each voxel. To model the spatial maps hierarchically, in addition to **P**^s^ for each subject, a consensus set of group-level parameters is also estimated to capture both the group mean and variance. **A**^sr^ represents mode time courses per subject and scan. The time courses also model signal and noise elements separately, where the signal element is constrained by the Haemodynamic Response Function (HRF) and the noise element follows a Gaussian distribution. Connectivity between the timecourses of functional modes can have a consensus structure across individuals. To model this simultaneously for population and individuals, a second hierarchy is defined over the precision matrix ***α***^*sr*^ using Wishart distributions. To further allow for modelling of haemodynamic responses that govern the Blood Oxygenation Level Dependent (BOLD) signal, HRF-constrained autocorrelations are incorporated in the modelling of this functional connectivity between the modes. **H**^sr^ denotes mode amplitudes. These are modelled using a multivariate normal distribution, with a Bayesian hierarchy between the group and the subject levels. **H**^sr^ is diagonal (one value per PFM) and captures timecourse variance for each subject and recording.

To find a solution for the probabilistic model described above, sPROFUMO uses stochastic Variational Bayesian (VB) inference. This includes dividing subjects from large data into small batches, and optimising the parameters of an approximating distribution *q*, with the aim that it is as close as possible to the true posterior. The group model is maintained across batches and continuously updated over time, and within each iteration of each batch, it is used for top-down regularisation of subject-specific matrix factorisations, from which the posterior distributions for subject-specific spatial maps, time course correlations and amplitudes are inferred. Subsequently, posterior distributions are accumulated across individuals and fed back to the group to update the group model. The model iterates between these two levels of estimation until convergence.

We used the model’s default parameters for application to both HCP and UKB datasets. The forget rate (*β*) was set to 0.6; this controls the degree to which global parameter updates rely on the current compared with previous batches. Delay parameter (*τ*) was set to 5; this controls the degree to which initial batches influence the overall inference. Batch size was set to 50 subjects, and the number of batches was tuned such that each subject will be visited 2.5 times on average (i.e., normally chosen in 2 to 3 batches). Given the different number of subjects in the two datasets, to achieve the abovementioned number of updates per subjects, the overall group model was set to be updated at least 5000 times for UKB and 4000 times for HCP. For the UK Biobank (taking 150-mode decomposition as an example), the model was run on 10 processing cores on a compute node, taking ∼210 hours to complete, and the memory usage peaked at ∼100GB. For the HCP (taking 150-mode decomposition as an example), the model was run on 10 processing cores on a compute node, taking ∼103 hours to complete, and the memory usage peaked at ∼90GB.

When applying to each dataset, and as a part of sPROFUMO’s internal initialisation, voxel-wise de-meaning and variance normalisation was applied to rfMRI recordings. Additionally, to initialise the model using a realistic set of initial group maps that can place the parameters within a realistic ballpark, sPROFUMO internally applies an online PCA algorithm called MELODIC’s Incremental Group-PCA (MIGP) (Smith et al., 2014) followed by variational ICA. These initial maps, together with hyperpriors, are used to initialise the group-level parameters before conducting the full Bayesian inference at subject- and group-level. It is worth noting that, to make the results fully comparable across HCP and UKB, the same set of initial maps were used to initialise both datasets. Two features in sPROFUMO help the model escape local minima: firstly, we use 10 random re-initialisations to generate the initial group-level maps; secondly, the mini-batching technique in stochastic variational Bayesian inference reduce the risk of getting stuck in local minima.

We estimated 25, 50, 100, and 150 PFMs, each used in a subset of the analyses, as elaborated in Results section. Initial maps were estimated for each dimensionality independently, since the number of the initialised modes will have to match the final number of the estimated modes. However, it is worth noting that the probabilistic framework has the capacity to switch off some of these modes in the final decomposition, if there is not enough posterior evidence in the data to support specific modes in higher dimensionalities (e.g., in noisier datasets or noisier modes for specific subjects). This feature, which reduces the risk of false positive modes, has been investigated in detail in our previous work (Farahibozorg et al., 2021).

After estimating PFMs with various dimensionalities, we validated whether each mode existed at both group-level and subject-level. A few artefactual modes (typically ∼2-3 due to, e.g., white matter artefacts) were manually removed from subsequent analyses. At subject-level, spatial and temporal aspects of the modes were used for evaluation. In particular, subject-to-group similarity of spatial maps is a useful metric. Very low similarities (∼<0.1) would indicate highly noisy estimation of a mode at subject-level. Whereas, artefactually high similarities (>0.99) would indicate that posterior parameters were not estimable at subject level, and thus, the spatial map for that specific subject-PFM was reverted back to group priors. An example of subject-group spatial map similarity for d=150 in UK Biobank is shown in **Figure S 3**. In addition to this, we used mode amplitudes (H^sr^) and standard deviation of spatial maps across voxels to identify PFMs with weak signal and subject level. PFMs included in the subsequent analyses were reliably estimated at both subject- and group-level.

##### 2.2.1.1 Volumetric vs CIFTI PFMs

sPROFUMO was applied to volumetric and CIFTI data from HCP separately, and group-level spatial maps were used as the basis for computing the reproducibility scores reported in Results section 3.2.2. In order to compare volumetric and CIFTI results, volumetric PFMs were mapped onto CIFTI using linear regression at the subject level. More specifically, for each subject and run, we used volumetric PFM timeseries (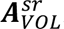) and amplitudes (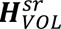), as well as raw CIFTI fMRI timeseries 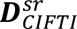. We computed spatial maps in CIFTI space (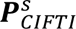) through back projection of 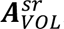 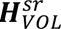 onto 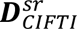 using linear regression with L2 regularisation. Regression betas were then converted to t-stat values. We next averaged 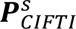 across subjects and obtained volume-to-surface mapped group spatial maps to compare against surface based spatial maps.

#### 2.2.2 Spatial ICA and Dual Regression

We applied the widely-used group level spatial ICA followed by dual regression (DR) to characterise subject-specific RSNs and compare against sPROFUMO results. In standard pipelines of large datasets such as HCP and UKB, ICA/ICA-DR have been used for RSN characterisation, thus providing a suitable baseline to compare sPROFUMO results against. We applied ICA to UKB data and identified 100 group-level spatial components using FSL *MELODIC,* and subsequently mapped these group-level results onto single subject data using FSL *dual_regression*. Subject-specific time courses for each mode and subject-specific spatial maps we obtained from stages 1 and 2 of Dual Regression, respectively. Mode amplitudes were computed as standard deviation of the time courses and functional connectivity was computed using Tikhonov/L2-regularised partial correlations. In order to make the final sPROFUMO and ICA/ICA-DR results fully comparable, the same PCA initialisation (using the MIGP approach) was used for both models. ICA-DR results were used to predict non-imaging phenotypes and compare PFM’s performance against.

### 2.3 Identifying temporally distinct subcomponents within large-scale RSNs

Low-dimensional representation of functional modes yields large-scale RSNs that are distributed across multiple brain areas. Matrix factorisation techniques (e.g. sPROFUMO or ICA) describe each of these modes with a single timecourse. We tested if there are distinct temporal subcomponents within each of these modes, and whether or not these subcomponents will be better represented by lowD large-scale RSNs vs highD mPFMs. For this purpose, we focused on 25-mode decomposition from HCP CIFTI rfMRI data of 1003 subjects, and conducted the following tests:

#### 2.3.1 Characterising distinct subcomponents within each conventional RSN

This step aimed to test whether a single prominent subcomponent can explain a majority of *group-level* temporal variance in the voxel-wise activity within each large-scale mode. For this purpose, we used PCA as follows:

a. N_voxel_ × N_time_ subject rfMRI timeseries were weighted by each subject PFM (d=25) spatial map, obtaining N_voxel_ × N_time_ weighted rfMRI timeseries (the data dimensions are unchanged, but voxels within a given map are up-weighted);
b. PCA was applied to each of these weighted timeseries, and 10 components were estimated per subject per mode, reducing the weighted subject fMRI data of each mode to N_voxel_ × 10. The resulting PCA maps (for each mode separately) were concatenated across subjects, resulting in a feature space of size N_voxel_ × (10*N_subject_). A second group-level PCA was next applied, and 10 group-level components were extracted per mode. The explained variance per PC was estimated (by considering the 10 singular values); the results of this analysis are reported in 3.1.1.

#### 2.3.2 Are temporal subcomponents best represented by mPFMs?

This step aimed to test whether multiple temporally distinct subcomponents exist within each large-scale RSN (i.e., distributed only), and whether subcomponents originating from large-scale RSNs can be better represented by large-scale (lowD) vs multiscale (highD) PFMs. This was done using temporal ICA, and further replicated using spatial ICA and PCA. This step, as well as the subsequent steps in 2.3.3 focused on *single-subject* modes.

a. Similar to step 2.3.1, N_voxel_ × N_time_ subject rfMRI timeseries were weighted by each subject PFM (d=25) spatial map, and 5 spatial PCs were extracted per mode per subject. The choice of 5 PCs was determined based on results of step 2.3.1, where PCs >6 were found to explain less than 3% of variance for all the PFMs (Figure S 4).
b. For each PFM the 5 PC maps were concatenated across subjects and group-level PCA was used to extract 5 spatial subcomponents per PFM.
c. These group-level PCs were then linearly regressed onto each subject’s PFM-weighted rfMRI timeseries (as described in step 2.3.1a), giving 5 temporal components per subject per mode, which were then concatenated across subjects to obtain 5 long timeseries per PFM.
d. Temporal ICA was applied to the resulting long timeseries to obtain timeseries for 5 temporally-independent subcomponents. These timeseries were split up into temporal chunks to obtain subject-specific tICA subcomponent timeseries.
e. Subject-specific tICA subcomponent timeseries were linearly regressed onto each subject’s PFM-weighted rfMRI timeseries (as described in step 2.3.2a) to obtain subject-specific tICA subcomponent spatial maps.
f. By correlating the subject-specific timecourses of each subcomponent with all the original lowD and highD mode timeseries, we applied a winner-takes-all approach to determine how many of the subcomponents originating from large-scale lowD modes were best-represented by lowD vs primary vs secondary mPFMs. An unpaired t-test was used to conduct statistical comparisons, and results were Bonferroni-corrected for multiple comparisons.

#### 2.3.3 Do secondary mPFMs emerge out of areas of spatial overlap and/or temporal co-activation?

1. This step aimed to test whether subcomponents identified in step 2.3.2 were exclusive to one lowD mode, or if they resided in the *temporal space* shared by multiple lowD modes:

a. Similar to step 2.3.2a, N_voxel_ × N_time_ subject fMRI timeseries were weighted by each subject PFM (d=25) spatial map, obtaining 25 weighted fMRI datasets per subject.
b. Linear regression was used to regress out the timecourses of every other (24) PFM from each PFM’s weighted fMRI. Then 5 PCs were extracted per mode per subject.
c. Steps 2.3.2b-f were repeated to test if subcomponents originating from the temporally-exclusive space of lowD modes were best-represented by lowD vs primary vs secondary mPFMs.
2. This step aimed to test whether subcomponents identified in step 2.3.2 were exclusive to a single lowD mode, or if they resided in the *spatial* space shared by multiple lowD PFMs:

a. Linear regression was used to regress out the spatial maps of every other (24) PFM from each PFM’s spatial map.
b. N_voxel_ × N_time_ subject fMRI timeseries were weighted by the resulting subject PFM (d=25) spatial map after regression, obtaining 25 weighted fMRI datasets per subject.
c. Linear regression was used to regress the spatial maps of every other (24) PFM from each PFM’s weighted fMRI. Then 5 PCs were extracted per mode per subject.
d. Steps 2.3.2b-f were repeated to test if subcomponents originating from the spatially-exclusive (non-overlapping) space occupied by lowD modes were best-represented by lowD vs primary vs secondary mPFMs.

### 2.4 Simulations

Based on real data (Methods section 2.3 and Results section 3.1), we found that Secondary mPFMs were likely due to the existence of temporally-distinct subcomponents within the Primary mPFMs, i.e., the ensemble would capture integrated-segregated modes of brain connectivity. Here we simulated datasets where overlapping distributed and localised RSNs exist in the brain, such that localised RSNs are spatial sub-nodes of distributed RSNs but have distinct timecourses. Using these simulations, we tested how well PFM decomposition can recover the co-existence of global-local RSNs in the brain, and compared its performance to spatial ICA as the standard technique. Details of simulations were as follows:

- Data: 10 resting state fMRI datasets were simulated, each consisting of 50 subjects and 2 runs per subject.
- 12 resting state modes were simulated in each dataset, and they were created from scratch in each dataset.
- 6 of these modes were distributed, covering >=2 brain areas, 6 of these modes were localised, covering 1 brain area. Distributed modes were allowed to spatially overlap with each other, such that on average, 1.3 modes included a given voxel.
- Localised modes were simulated such that they were primarily a spatial sub-node of one distributed mode, while spatially overlapping with two or multiple distributed modes.
- Data properties per subject and run: 10,000 voxels, 300 time points at a TR of 0.72s. Data was created as detailed below:

We defined a set of spatial maps **P_g_** at the group level. Each mode consisted of one or more randomly-selected contiguous blocks of voxels (i.e., parcels). Signal weights (per voxel per mode) were drawn from a Gamma distribution. Subject-specific spatial maps, **P_s_**, were defined based on the group maps by adding background Gaussian noise and applying spatial warps. Subject maps were generated to be spatially misaligned in reference to the group, with any a given subject mode having on average 83% overlap with the group-average (i.e., 17% spatial misalignment). Time courses were generated independently for each session, subject and mode to mimic the unconstrained resting state data (but following the between-mode temporal correlation structure described next). We assigned a hierarchical link between the group and subjects’ temporal correlation matrices, following a Wishart distribution. This was done to obtain a consistent functional connectivity pattern between subjects and the group. Time courses were initially simulated as semi-Gaussian neural time courses with amplified frequencies < 0.1Hz, and subsequently convolved with a random draw from the FLOBS haemodynamic basis functions (Woolrich et al., 2004) to mimic realistic BOLD signal. Finally, data were created using outer product of subject mode spatial maps and timecourses, and random noise was added to the outer product to create space-time data matrices. More details of simulation parameters are available in (Harrison et al., 2020).

These simulated modes were then estimated using PFM decomposition and ICA-Dual Regression, and results were compared with the ground truth. We initialised both models based on the same set of spatial bases from MIGP, to ensure that the observed differences were not due to initialisation of the probabilistic models.

### 2.5 mPFMs vs HCP-MMP

We applied sPROFUMO to cortical rfMRI data of 446 HCP subjects (matching the number of subjects for which HCP-MMP was available) and estimated 360 PFMs. The two decompositions were compared at two levels: First, we computed group correspondence between mPFMs and HCP-MMP. We binarised group-level mPFMs’ spatial maps by hard thresholding at 0.3 and computed their dice similarity to group-level parcels in HCP-MMP. Second, we examined subject-level correspondence by binarising subject-level mPFMs’ spatial maps by hard thresholding, and finding dice similarity between binarised mPFMs and subject-level parcels in HCP-MMP. The modes were paired based on group-level matching, and this ordering was used in both group-level and subject-level visualisations.

### 2.6 Out-of-sample prediction pipeline

We investigated the link between the spatial/temporal properties of mPFMs and a range of phenotypes in UKB and HCP. This was done using out-of-sample predictions in UKB and Canonical Correlation Analysis (CCA) in HCP. These accuracies were used to compare mPFMs vs conventional large-scale RSNs, primary vs secondary mPFMs, and mPFMs vs spatial ICA and Schaefer parcellation of same dimensionality. Depending on the question at hand in each subsection, we conducted predictions based on all or some of the following features: a) Mode spatial maps, b) Mode temporal network matrices (TNETs or temporal NetMats; partial correlation matrix between mode timecourses), c) Mode spatial NetMats (SNETS, the full spatial correlation matrix between mode maps).

#### 2.6.1 Imaging and non-imaging derived phenotypes (IDPs and nIDPs)

We used 893 Imaging-derived and non-Imaging-Derived phenotypes (IDPs and nIDPs) from UK Biobank as targets in the prediction pipeline. These included IDPs related to Grey Matter, White Matter and task fMRI, as well as nIDPs related to cognitive scores and Cardiovascular health metrics. A brief summary of these phenotypes is included in this subsection, and a full list is included in Table S 7-S 13.

##### Grey Matter

148 IDPs related to GM area and 148 IDPs related to GM thickness, generated with Freesurfer by parcellation of the white surface using Destrieux (a2009s) parcellation. More details about these IDPs (e.g. histograms) can be found on the UKB website: https://biobank.ctsu.ox.ac.uk/crystal/label.cgi?id=197.

##### White matter

3 IDPs extracted based on the volume of white matter hyperintensities using BIANCA. 162 IDPs extracted based on mean intensity for dtifit outputs (FA, MD, MO3) and NODDI outputs (ICVF, OD, ISOVF) for the 27 tracts segmented with the probabilistic tractography analysis. 288 IDPs extracted based on mean intensity for dtifit outputs (FA, MD, MO) and NODDI outputs (ICVF, OD, ISOVF) for the 48 tracts segmented with the TBSS-like analysis.

##### Task fMRI

UKB includes data for one task, which involves brain responses to images of emotionally-valenced faces and shapes. 1 IDP, called ‘contrast’ in the results section, was estimated as 90th percentile (z-statistic) of activity for response to faces vs shapes (https://biobank.ctsu.ox.ac.uk/crystal/field.cgi?id=25766).

##### Cognitive

Starting from 1172 nIDPs related to cognitive tests, we narrowed these down to 68 using two criteria. Firstly, one of the authors prefiltered the outcome measures manually, by only including active measures that were indicative of subject’s performance. For example, among outputs directly related to “Reaction Time”, which entails viewing two cards (A and B) and pressing buttons when identical, “Mean time to correctly identify matches”, was included whereas “Index for card A in round” was excluded. Secondly, we only selected tests that had a non-NaN value in at least 25% of the subjects. The final list of 68 tests belonged to the following categories: *Reaction time, Trail making, Matrix pattern completion, Numeric memory, Prospective memory, Pairs matching, Symbol digit substitution* and *Fluid intelligence*. More details about these nIDPs are available here: https://biobank.ctsu.ox.ac.uk/crystal/label.cgi?id=100026.

##### Cardiovascular health metrics

Starting from 992 nIDPs related to cardiovascular health, we narrowed these down to 77 using two criteria. Firstly, one of the authors prefiltered the outcome measures manually, by excluding metrics that were not directly related to health. For example, “LV stroke volume (2.0)” and “Systolic blood pressure, manual reading (0.0)” were included whereas “Completion status of test (0.0)” and “Program category (0.0)” were excluded. Secondly, we only selected tests that had a non-NaN value in at least 25% of the subjects. More details about these nIDPs are available in the following links: https://biobank.ctsu.ox.ac.uk/crystal/label.cgi?id=100011, https://biobank.ctsu.ox.ac.uk/crystal/label.cgi?id=104 and https://biobank.ctsu.ox.ac.uk/crystal/label.cgi?id=100012

#### 2.6.2 Confound removal in UK Biobank

When conducting brain-behaviour analyses in big data, e.g., association or out-of-sample predictions, imaging confounds can significantly distort the interpretability of the results (Alfaro-Almagro et al., 2020; Snoek et al., 2019). For example, if a common confounding factor such as head size or head motion is correlated with both brain features and behavioural targets, this is likely to inflate our prediction accuracies (Snoek et al., 2019). Therefore, we opted to de-confound the data before conducting any brain-behaviour analyses. This was done by linearly regressing out the confounds from both imaging and non-imaging variables.

For UK Biobank, we started with a comprehensive set of 602 confounds proposed by Alfaro-Almagro et al. (2020). We reduced this set by: a) selecting conventional confounds including age, age squared, sex, age × sex, site, head size and head motion; b) reducing the remaining confounds using Principal Component Analysis. The top PCs that explained >85% of the variance were kept. These two steps yielded 82 variables. Importantly, we applied deconfounding taking into account cross-validation folds in order to avoid leakage of information from test to train data, as proposed by (Snoek et al., 2019).

#### 2.6.3 Elastic-Net prediction and cross-validation

We used Python 3.6.5 and scikit-learn 0.19.1 (Pedregosa et al., 2011) to set up the prediction pipeline. We used ElasticNet regression and nested 5-fold cross validations to conduct out-of-sample predictions, with 20/80 test/train ratio. Next, we Gaussianised each predictor and target variable across subjects using quantile transformation (*QuantileTransformer*). In each cross-validation loop, the training set was used to compute the quantile transforms which were then applied to both train and test data. Similarly, deconfounding was also conducted within each cross-validation loop, where training set was used to estimate the regression parameters (or “betas”), which were then applied to de-confound both train and test data. Finally, we used *ElasticNetCV* to predict target variables in the test set. Hyperparameters related to ElasticNet; i.e., ratio of Lasso to Lasso+Ridge regularisation (*L1 ratio* varied between [0.1,0.5,0.7,0.9,0.95,0.99,1.0] with 10 alphas per *L1 ratio*) we optimised using nested cross-validations within the training set. Finally, we computed correlations between estimated and actual values of the target variables across subjects in the test set, which will be reported as prediction accuracies.

#### 2.6.4 Feature spaces: large-scale vs multiscale PFMs

The comparison between conventional large-scale RSNs and mPFMs aimed to determine any added benefit of the latter, and additionally to determine the added benefit of having both types of primary and secondary modes in a decomposition, as opposed to one type only. For this purpose, we focussed on interactions between the modes, i.e., spatial/temporal correlations, and their prediction power for phenotypes in UKB. Specifically, the comparison was made between prediction power of correlations in three scenarios: large-scale RSNs vs mPFMs; between primary and secondary mPFMs vs within primary mPFMs; between primary and secondary mPFMs vs within secondary mPFMs.

To summarise mode interactions, two types of feature matrices were used, Spatial NetMats (SNETs) and partial temporal NetMats (TNETs). TNETs were computed based on precision matrices and with a Tikhonov regularisation parameter 0.01. SNETs and TNETs are of size N_mode_ × N_mode_ for each subject in each PFM decomposition.

Based on highD decomposition (d=100), we divided each of the NetMats into three sub-matrices

1. SNET_pri2pri_ : N_pri_ × N_pri_ and TNET_pri2pri_ : N_pri_ × N_pri_, with N_pri_ = 25
2. SNET_sec2sec_ : N_sec_ × N_sec_ and TNET_sec2sec_ : N_sec_ × N_sec_, with N_sec_ = 75
3. SNET_pri2sec_ : N_pri_ × N_sec_ and TNET_pri2sec_ : N_pri_ × N_sec_
4. Based on large-scale lowD decomposition (d=25), we obtained SNET_low_dim_ : N_lowD_ × N_lowD_ and TNET_low_dim_ : N_lowD_ × N_lowD_

We applied the Fisher r-to-Z transformation, and unwrapped these matrices into 1D vectors. Since Pri2Pri, Sec2Sec and LowD2LowD matrices (but not Pri2Sec) are symmetric, only the upper diagonal elements were kept before unwrapping. Next, we used SVD to dimension-reduce each of these eight N_subject_ × N_edges_ matrices to feature matrices of size N_subject_ × 200, which were fed into ElasticNet. The dimensionality reduction was done to obtain the same number of features to use in predictions, reducing the possibility of one mode type outperforming due to the number of features. Five-fold cross-validations were performed and accuracies computed as detailed in section 2.6.3. Cross-validations were repeated k times for each phenotype category, k being: GM: 20, WM: 20, task fMRI-condition: 200 (since there is only 1 phenotype in this category), task fMRI-contrast: 200, cognitive: 20, cardiovascular metrics: 20.

For each IDP/nIDP category, accuracies were computed for each phenotype, and pooled across the number of phenotypes in that category (N_phenotype_) and number of repeats (k), yielding two vectors of size k* N_phenotype_ to be compared between different mode types.

#### 2.6.5 Feature spaces: mPFMs vs standard techniques

##### 2.6.5.1 mPFMs and spatial ICA features

The comparison of mPFMs to highD ICA aimed to examine any added benefit of mPFMs over state-of-the-art RSN decompositions for capturing individualistic traits and phenotype predictions. For this purpose, three types of feature matrices were used in predictions, spatial maps (SMAPs), SNETs, and TNETs. HighD decompositions with 100 RSNs from UKB data were used in these analyses. This dimensionality was optimised based on a left-out phenotype (age): we compared age prediction using 50, 100, 150, and 200 dimensional decompositions and found 100 to be optimal for both mPFMs and highD ICA. It is worth noting that d=100 was found to be optimal here for UK Biobank data. The optimal choice may vary for datasets with a smaller number of subjects, or datasets with longer recording sessions per subject. Feature spaces were estimated using a pipeline similar to the Uni-mode predictions described in our previous study (Farahibozorg et al., 2021):

- For SMAPs, estimating 100 modes resulted in a matrix of N_subject_ × N_voxel_ × 100. We performed dimensionality reduction across the second dimension (per mode) using sparse dictionary learning (Mairal et al., 2010) and obtained feature matrix of size (N_subject_ × 500) × 100. Next, we used the feature space of each PFM separately in phenotype predictions.
- For SNETs and TNETs, we started with matrices of size N_subject_ × N_mode_ × N_mode_. After applying the Fisher r-to-Z transformation per subject, we used rows of correlation matrices (i.e., one per mode) for each subject without additional dimensionality reduction, thus obtaining 100 feature matrices of size N_subject_ × 99 from SNETs and 100 feature matrices of size N_subject_ × 99 from TNETs that were used separately for predictions. TNETs were computed using partial correlations with Tikhonov regularisation.

##### 2.6.5.2 Schaefer Parcellation features

The standard Schaefer parcellation in MNI space, Schaefer2018_100Parcels_17Networks_order_FSLMNI152_2mm.nii.gz, from https://github.com/ThomasYeoLab/CBIG/tree/master/stable_projects/brain_parcellation/Schaefer2018_LocalGlobal/Parcellations/MNI was applied to pre-processed fMRI timeseries of UK Biobank subjects; timecourses of voxels within each parcel were averaged to obtain one timecourse per parcel/mode. Subsequently, similar to the PFM/sICA procedure explained above, TNET features were computed from these timecourses using Fisher r-to-Z transformed partial correlations with Tikhonov regularisation. These resulted in 100 feature matrices of size N_subject_ × 99 which were used in uni-mode predictions.

##### 2.6.5.3 Summary of features from mPFM/sICA/Schaefer

We used features of each mode separately in predictions.

On the one hand, for PFM and sICA, SMAP, SNET and TNET feature matrices were concatenated horizontally, yielding 100 features matrices of size N_subject_ × 799 to use in cross-validated ElasticNet predictions, as elaborated in section 2.6. Accuracies were computed for each mode and pooled across modes to compare PFMs and ICA-DR.

On the other hand, mPFMs and spatial ICA both yield spatial and temporal features, whereas hard parcellations such as Schaefer generate TNET features only. This is due to the fact that hard parcellations yield binarised spatial topographies for the modes/parcels, with fixed hard boundaries between the parcels. Therefore, the number of features derived from the Schaefer parcellation will be lower. To balance this number across the three methods, we used cross-validated feature selection based on correlation to the target in the training set to select a subset of features from mPFM/sICA, matching the number of features (99 per mode) from the Schaefer parcellation.

For each phenotype category (e.g., cardiovascular, cognitive etc.), accuracies were computed for each mode and each phenotype, and pooled across the number of phenotypes in that category (N_phenotype_) and number of modes (100), yielding three vectors of length 100* N_phenotype_ to compare mPFMs with ICA-DR and the Schaefer Parcellation.

### 2.7 Canonical Correlation Analysis

Canonical Correlation Analysis (CCA) was used to find a single multivariate mapping between a set of PFM features and a set of non-imaging variables. Each CCA component estimates a linear combination of PFM features and a linear combination of phenotypes, such that the transformed outputs are maximally correlated. In order words, CCA components project data onto common axes of subject variability that co-vary between brain and behaviour. In previous studies of rfMRI functional connectivity using spatial ICA, CCA has identified a positive-negative axis of brain-behaviour associations in HCP data (Smith et al., 2015). Here, using HCP and focussing on the same set of phenotypes, we conduced CCA aiming to examine the behavioural relevance of mPFMs in the context of existing literature. 1001 out of 1003 subjects that were included in both fMRI and phenotype recordings were included in this analysis.

#### 2.7.1 Non-imaging phenotypes

We used 158 phenotypes for the CCA. These were sub-selected from a wider range of phenotypes using a set of criteria described in (Smith et al., 2015), and included metrics related to cognition such as various measures of fluid intelligence, executive function, language, episodic memory, working memory, attention; metrics of emotion such as life satisfaction, friendship, loneliness; metrics of psychiatric and life function such as depression, anxiety, aggression; metrics of alertness related to sleep, five metrics of personality related to agreeableness, extraversion, neuroticism, conscientiousness and openness, metrics of physical health related to sensory-motor function, and lifestyle metrics related to substance use such as alcohol, tobacco and drug. A full list of these phenotypes is presented in Table S 14.

#### 2.7.2 Confound removal in HCP

Similar to section 2.6.2, we de-confounded both feature and phenotype matrices before conducting CCA. For this purpose, we used 13 imaging confounds that are included regularly in HCP studies: acquisition reconstruction software version; age; age squared; sex; age × sex; sex × age squared; race, ethnicity; height; weight; a summary statistic quantifying average subject head motion during the resting-state fMRI acquisitions; the cube-root of total brain volume (including ventricles), as estimated by FreeSurfer; the cube-root of total intracranial volume, as estimated by FreeSurfer.

#### 2.7.3 mPFM feature spaces

For CCA, we need a single feature matrix of size N_subject_ × N_feature_ estimated from brain imaging to be paired with a single phenotype matrix of size N_subject_ × N_phenotype_ from behavioural/non-imaging traits. Therefore, we required a further condensed feature space compared to what described earlier in trait prediction. We extracted feature summaries separately from SMAPs, SNETs, and TNETs, and ran CCA using each feature type independently. Feature summaries were created as follows:

- For the spatial maps (SMAPs), estimating 150 modes resulted in a **P**_grand_ matrix of (N_subject_ × N_voxel_)x 150, yielding ∼35 million features per subject. To extract a few hundred features that can meaningfully capture the essence of subject SMAPs, we used unsupervised learning by applying FMRIB’s Linked ICA for big data (BigFLICA) (Gong et al., 2021; Groves et al., 2011). This is an ICA framework originally proposed for multimodal data fusion, and was applied here in two steps. First, subject SMAPs were dimension-reduced across voxels using sparse dictionary learning, yielding a matrix of size (N_subject_ × 1000) × 150. Next, each PFM was used as a separate “modality” within bigFLICA to obtain a feature matrix of size N_subject_ × 500. Using FLICA here allows us to preserve subject-specific variations in each mPFM and efficiently summarise them across numerous modes to obtain a set of independent features to characterise the sources of population variations in mPFMs.
- For the spatial and partial temporal correlation matrices (SNET, TNET), we started with matrices of size N_mode_ × N_mode_ for each subject. After applying the Fisher r-to-Z transformation, we flattened these matrices by taking the above-diagonal elements. With 150 PFMs, this resulted in two feature matrices, one for SNET another for TNET, each of size N_subject_ × (150*149/2) = N_subject_ × 11175.

Each of these three matrices were used separately in CCA estimation, and results were combined post-hoc, as elaborated in the next subsection.

#### 2.7.4 Conducting CCA

CCA analysis consisted of the following steps:

1. One side of CCA received mPFM feature matrices as input: SMAPs: 1001 × 500 (N_subject_ × N_FLICA_); SNETs: 1001 × 11175 (N_subject_ × N_NET_ELEMENT_); TNETs: 1001 × 11175 (N_subject_ × N_NET_ELEMENT_). Preprocessing steps similar to (Smith et al., 2015) were applied to these matrices, which included normalisation and dimensionality reduction using SVD. Each feature matrix was reduced to a matrix of size 1001 × 50. The dimension reduction helps to avoid an over-determined (rank deficient) CCA solution.
2. The other side of CCA received phenotype matrices as input: 1001 × 158. Preprocessing steps similar to (Smith et al., 2015) were applied to these matrices, which included inverse Gaussian transformation and dimensionality reduction using SVD. The phenotype matrix was reduced to a matrix of size 1001 × 50.
3. Imaging confounds were regressed out of both phenotype and PFM feature matrices, as elaborated in section 2.6.2.
4. CCA was conducted for three pair-wise comparisons: 1) PFM SMAP vs phenotypes; 2) PFM SNET vs phenotypes; 3) PFM TNET vs phenotypes. This was done using the ‘canoncorr’ function in Matlab.
5. CCA yields a linear transformation of PFM feature matrix (X) and phenotype feature matrix (Y) so as to maximise their correlation; i.e. Y*A=U ∼ X*B=V, where U and V are the linearly-transformed versions of ICA-DR and PFM feature matrices, respectively.
6. By finding correlations between columns of U and V for the top 50 CCA components, we estimated shared variances for each pairwise comparison.
7. We finally tested how many of the CCA components were significantly correlated. For this purpose, we conducted multi-level block permutations (Winkler et al., 2015) which takes family structure of HCP data into account. In each iteration, X in step 5 was kept fixed, and rows of Y were randomly permuted (i.e., permuting subjects, while keeping family members together), and step 6 was repeated. Across 10,000 permutations, we constructed a null distribution for the correlations of the top 50 CCA components. The correlation value corresponding to the top 5% of the null distribution for the *first* CCA component was used as threshold for p-value<0.05 significance level for all the CCA components. This yields a significance threshold that is Family-Wise Error-rate (FWE) corrected for multiple comparisons (across CCA components).

## 3 Results

**Figure 2** illustrates four examples of the well-known large-scale RSNs, estimated using ICA in HCP’s standard pipelines. Underneath, we have shown one primary mPFM (the closest spatial match) and two example secondary mPFMs, matching subsystems of these RSNs. Default Modes (DMN 1&2) closely resemble the two DMNs that have been reported in the previous literature (Andrews-Hanna et al., 2010), and distinct from each other in various ways, e.g.,: DMN1 features a para-hippocampal node that is missing in DMN2; DMN1 encompasses posterior IPL and Angular Gyrus, whereas DMN2 encompasses anterior IPL; lateral temporal areas such as the anterior temporal lobes are more pronounced in DMN2; DMN1 encompasses ventromedial PFC, whereas DMN2 encompasses dorsomedial PFC. The secondary mPFMs under each DMN highlight a few of their subnodes that contribute to distinct functionalities. Similarly, the illustrated Language network has been reported in previous meta-analytic analyses of language processing in the brain (Lipkin et al., 2022), featuring a predominantly left-lateralised network that covers temporal, inferior parietal, inferior frontal and pre-motor regions. Each of the secondary mPFMs then highlight subparts of this network. Finally, one of the brain’s motor networks located at the central sulcus and two of its subsystems are shown on the right hand side of the figure. In the next section, we dive into understanding what these mPFMs are, and why they appear as separate, yet overlapping, entities in high-dimensional decomposition of resting-state fMRI (rfMRI) data using sPROFUMO.

**Figure 2.**
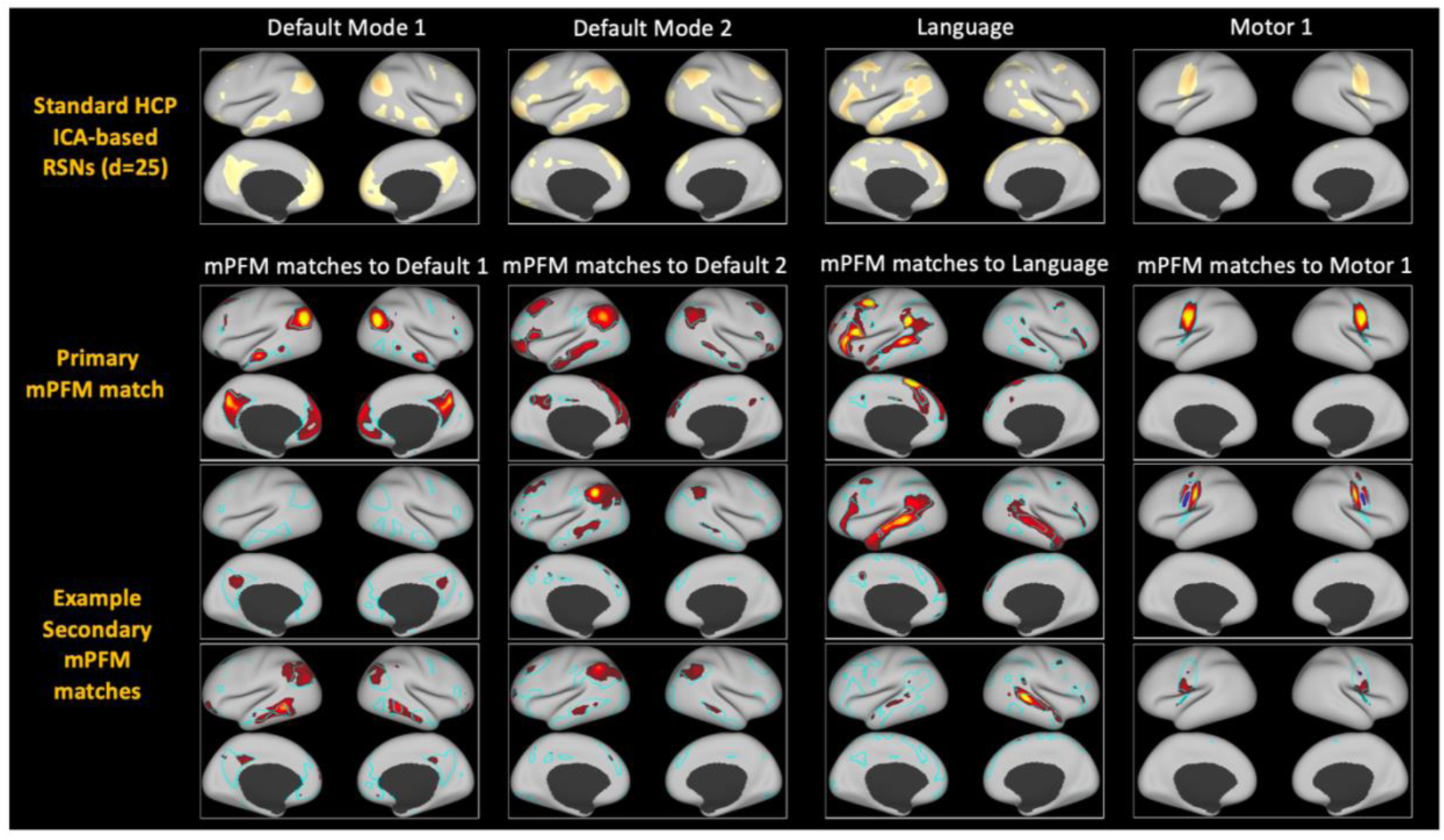
Four well-known RSNs and example primary and secondary mPFM matches to each. Top panel shows conventional RSNs, included in HCP’s standard Resting State Networks (RSNs) releases, and estimated using ICA (d=25). mPFMs are a subset of the RSNs that are obtained from a high-dimensional (d=150) decomposition of cortical rfMRI data from 1003 HCP subjects. mPFMs are overlayed on cyan-coloured borders of the ICA-based RSNs for easier visual comparison. As can be noted, primary mPFMs are, in principle, the well-known RSNs: Default Mode 1 (DMN1), Default Mode 2, Language and Motor 1 networks. Secondary mPFMs depicted underneath show subsystems of these networks.

We applied sPROFUMO to rfMRI data from 1003 HCP subjects and separately to 4999 UK Biobank subjects. Results predominantly focus on a 25-mode low-dimensional (lowD) PFM decomposition and its comparison to higher-dimensional (highD, >100) PFM decompositions, to investigate the functional relevance and the added benefit of mPFMs. Table S 1 includes a summary of the terminology used in the paper.

The following subsections aim to unravel the properties of the brain function that give rise to mPFMs, validating them based on simulations and real data, as well as demonstrating the utility of mPFMs for capturing individualistic traits and prediction of cognition and health. HCP data, compared to UK Biobank, provides higher data quality and longer fMRI recordings (1 hour vs ∼6 minutes) per subject. Thus, HCP data was primarily used for untangling the functional relevance of mPFMs, and how they yield a more comprehensive summary of brain activity and connectivity. UK Biobank, given its larger sample size and more extensive phenotyping, was primarily used for evaluating mPFMs’ performance in out-of-sample prediction of phenotypes. Both datasets were used for reproducibility analyses.

### 3.1 The functional relevance of mPFMs

Focussing on 25 and 150 sPROFUMO RSNs from rfMRI data of 1003 HCP subjects, we investigated the properties of the brain function that underlie mPFMs, and their added benefit over large-scale (distributed-only) decompositions for capturing multiscale information processing in the brain. We use the following terminology in the rest of the paper (see Figure 1): a) large-scale RSNs (yellow): 25 modes from low-dimensional decomposition; b) “primary” mPFMs (orange): 25 distributed modes from high-dimensional PFM decomposition that best-match the low-dimensional RSNs; c) “secondary” mPFMs (blue): remaining 125 modes from high-dimensional PFM decomposition.

#### 3.1.1 Multiple distinct subcomponents within each conventional RSN

Each functional mode is typically characterised by two key characteristics: a) the spatial topography across brain voxels (i.e., a spatial map), from which one can derive spatial correlations or overlaps with other modes; b) a timecourse, and the resulting temporal correlations with other modes (i.e., functional connectivity).

This representation assumes that voxels within a mode are highly correlated with each other, such that they can be summarised with one consensus timecourse. In other words, it assumes that temporal variability across voxels within each RSN’s spatial map is negligible. To determine whether lowD decomposition complies with this assumption, we asked: how many components are needed to explain a majority of variance in the voxel-wise activity within each large-scale RSN? We used Principal Component Analysis (PCA), which projects data onto new orthogonal axes of variation, and allows to identify distinct subcomponents based on the proportion of variance explained (details in 2.3.1). We found that (Figure S 4) the first Principal Component could only explain up to 53.2% of variance of temporal variability, with the second to fifth Principal Components each explaining ∼5-10% of variance.

Next, we applied follow-up temporal ICA on data of each mode from lowD decomposition, as detailed in 2.3.2. Temporal ICA projects data onto statistically independent axes of variations and allow us to identify *temporally independent subcomponents* more interpretably. We restricted the number of subcomponents to 5 per mode; i.e., 5 subcomponents for each of the 25 modes. Figure 3a shows examples of these subcomponents for the Default Mode and Language Networks. We found that these subcomponents were spatially and temporally distinct, with one subcomponent being spatiotemporally best-represented by the original large-scale decomposition compared with the rest of the subcomponents (Figure S 5). Figure 3b shows the temporal correlation of each of the best-matching subcomponents to the large-scale RSN that they originated from. Interestingly, we found that even the temporal representation of the best-represented subcomponent ranged from Pearson correlation coefficient = 0.22±0.13 to 0.70±0.09 across the 25 large-scale RSNs (i.e., in many cases there is not a strong temporal match). We replicated this finding using two alternative subcomponent identification techniques, spatial ICA and PCA (Figure S 6).

**Figure 3.**
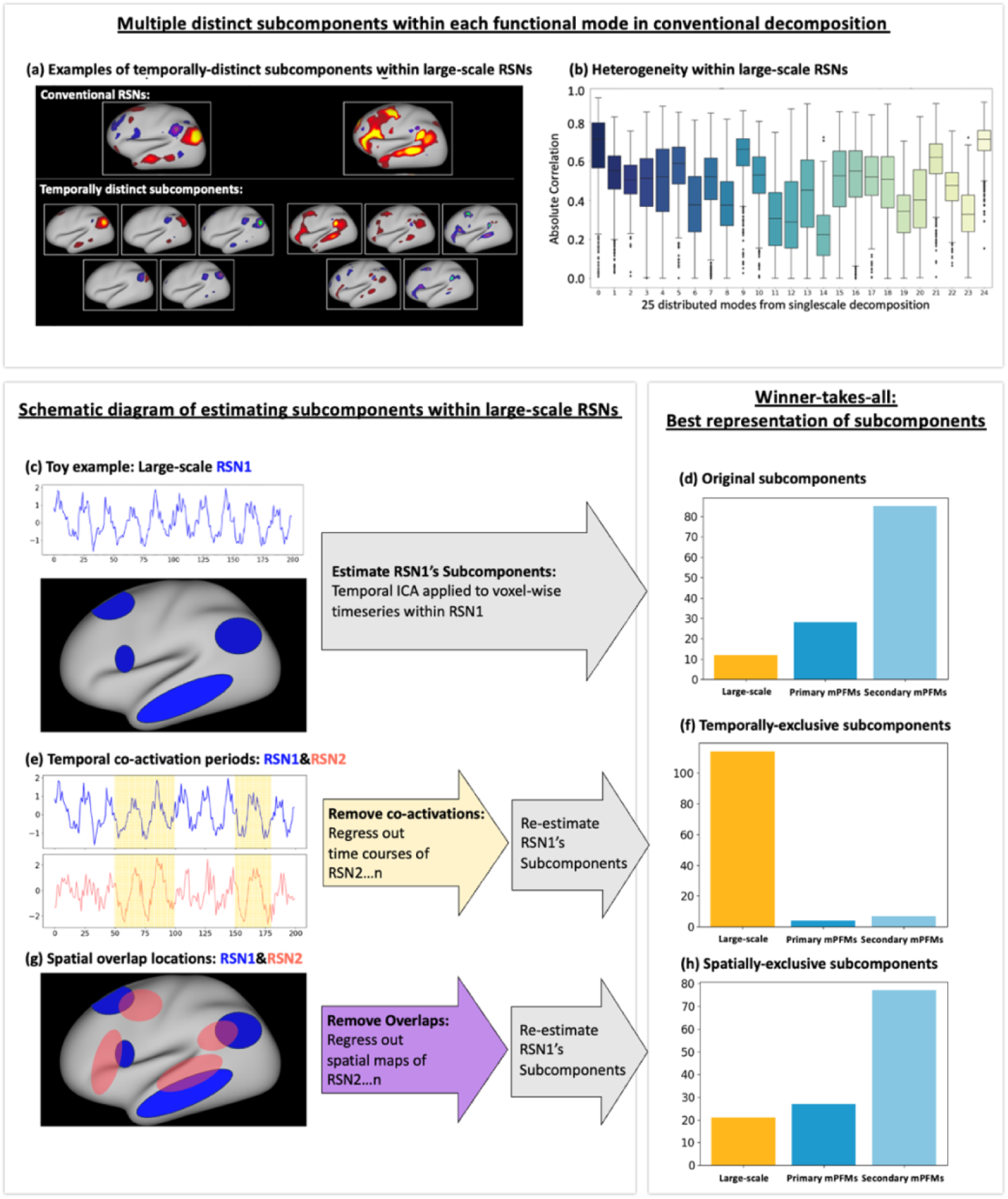
Temporally distinct subcomponents within conventional large-scale RSNs give rise to multiscale Probabilistic Functional Modes (mPFMs) A low-dimensional PFM decomposition consisting of 25 modes was estimated from cortical rfMRI data of 1003 HCP subjects, yielding conventional large-scale RSNs. Follow-up temporal ICA was applied to voxel-wise timeseries within each RSN to identify temporally-distinct subcomponents. a) Examples of two conventional RSNs and 5 subcomponents within each RSN. b) Temporal correlation of the best-matching subcomponents to the large-scale RSNs that they originated from. c) A schematic illustration of subcomponent estimation. d) A winner-takes-all approach was applied, and it was found that of the 25×5 subcomponents, only 12 were best represented by the large-scale RSNs that they originated from, whereas 28 and 85 subcomponents were better represented by primary and by secondary mPFMs, respectively. e) A schematic illustration of estimating temporally-exclusive subcomponents. f) A winner-takes-all approach was applied, and it was found that of the 25×5 temporally-exclusive subcomponents, 114 (i.e., the majority) were best represented by large-scale RSNs, 4 by primary and 7 by secondary mPFMs. g) A schematic illustration of estimating spatially-exclusive subcomponents. h) A winner-takes-all approach was applied, and it was found that of the 25×5 spatially-exclusive subcomponents, 21 were best represented by large-scale RSNs, 27 by primary and 77 by secondary mPFMs. These results show that temporally-distinct subcomponents within large-scale RSNs give rise to secondary PFMs, and that periods of temporal co-activation between subnetworks of large-scale RSNs give rise to secondary PFMs.

These results provide evidence that voxel-wise temporal variability within large-scale RSNs results in *temporally distinct* subcomponents that are not well-represented by one timecourse, as assigned in conventional RSN decompositions. Instead, multiple subcomponents are needed to explain this voxel-wise temporal variability.

#### 3.1.2 Temporal subcomponents were best represented by mPFMs

We next tested if the 5×25 subcomponents described in the previous subsection were better represented by mPFMs, compared with lowD large-scale RSNs alone. For this purpose, we applied a winner-takes-all approach: the timecourse of each of the subcomponents was correlated with: a) large-scale RSNs; b) primary mPFMs and c) secondary mPFMs. As shown in Figure 3d, we found that only 12 of the 125 subcomponents were temporally best represented by the large-scale RSNs, 9 of which were significantly higher (Bonferroni-corrected, Table S 2). The remaining 28 and 85 subcomponents were better represented by the primary and secondary mPFMs, respectively (28/28 and 83/85 statistically significant). Therefore, the ensemble of multiscale modes in mPFMs can capture the temporally distinct subcomponents within conventional large-scale RSNs.

#### 3.1.3 Secondary mPFMs were due to temporal co-activation periods of multiple large-scale RSNs

Resting state networks in the brain might be spatially and temporally correlated. Therefore, it is plausible that the subcomponents that give rise to the secondary mPFMs are either due to spatially overlapping regions or due to distinct temporal co-activation periods of multiple large-scale RSNs. To test this hypothesis, we compared the original subcomponents (described in the previous subsection) with two alternative scenarios: a) temporally-exclusive subcomponents: estimated from each large-scale RSN after the timecourses of the other 24 RSNs were regressed out of its timecourse; b) spatially-exclusive subcomponents: estimated from each large-scale RSN after spatial maps of the other 24 RSNs were regressed out of its spatial map. Conceptual illustrations of these two analyses are included in Figure 3e and g, respectively.

As shown in Figure 3f, we found that a majority of the temporally-exclusive subcomponents (114 out of 125, with 109 statistically significant, Table S 3) were temporally best represented by the *lowD decomposition*, unlike the original subcomponents that were best represented by mPFMs. Conversely, as shown in Figure 3h, the majority of the spatially-exclusive subcomponents (104 out of 125, 100 statistically significant, Table S 4) were instead best represented by primary and secondary mPFMs, i.e., similar to original subcomponents. All results were Bonferroni-corrected for multiple comparisons.

Put together, these analyses suggest that temporal co-activation periods of two or multiple large-scale RSNs from lowD decomposition give rise to new mixed-scale RSNs that are temporally distinct from the existing RSNs. These emerge as secondary mPFMs. In other words, secondary mPFMs are likely due to transient dynamic connectivity between subnetworks of large-scale modes, that are eliminated by temporal averaging (across voxels and/or timepoints) in low-dimensional decompositions.

#### 3.1.4 Functional connectivity modelling using mPFMs

Having examined timecourse modelling in mPFMs, we next investigated the functional connectivity (i.e., temporal correlations) between the modes to compare large-scale RSNs (lowD, d=25) with mPFMs (d=100 and 150). This was done using 4999 subjects in UK Biobank (Figure S 7a), and separately using 1003 subjects in HCP (Figure S 7b). Figure S 7-right shows the histograms of between-subject consistency of functional connectivity. We observed that in both datasets, between-subject consistency was increased with dimensionality. In HCP, this was increased from 0.19±0.09 for d=25 to 0.46±0.06 and 0.59±0.05 for d=100 and 150, respectively. In UK Biobank, this was increased from 0.07±0.07 for d=25 to 0.43±0.13 and 0.70±0.12 for d=100 and 150, respectively. Therefore, mPFMs yielded more consistent functional connectivity values than lowD. Interestingly, subject-specific functional connectivity became sparser for mPFMs (Figure S 7-left). This sparsity can be understood in light of the previous subcomponent analysis: by attempting to merge the time courses of multiple temporally distinct subcomponents, conventional RSNs can yield spurious connectivity between large-scale modes, which in reality can be due to one or a few of the underlying subcomponents. mPFMs capture the subcomponents using secondary modes, thus reducing such spurious connectivity, and yielding sparser subject-specific functional connectivity. As a result, functional connectivity in mPFMs becomes less contaminated by modelling inaccuracies, and more consistent across individuals.

#### 3.1.5 Cross-scale interactions of mPFMs predict individualistic traits

mPFMs can directly characterise the *spatiotemporal correlations between modes across multiple scales of information processing in the brain*. We evaluated the utility of temporal and spatial correlations between mPFMs for out-of-sample prediction of a range of phenotypes in UK Biobank. This was performed using volumetric rfMRI data from 4999 UK Biobank subjects and 893 phenotypes divided into 6 categories: region-wise cortical area (148 phenotypes) and thickness (148), White Matter (WM) tracts’ microstructure (451), task fMRI response to the contrast of emotional faces and shapes (1), cognition (68) and cardiovascular health (77), see 2.6.1 for details of phenotypes, feature extraction and prediction pipeline. Figure 4a&b, show functional and spatial connectivity of large-scale RSNs (d=25) and mPFMs (d=100) for an example subject. As demonstrated in these panels, cross-subject consistency of both functional and spatial connectivity was higher in mPFMs compared with large-scale RSNs.

**Figure 4.**
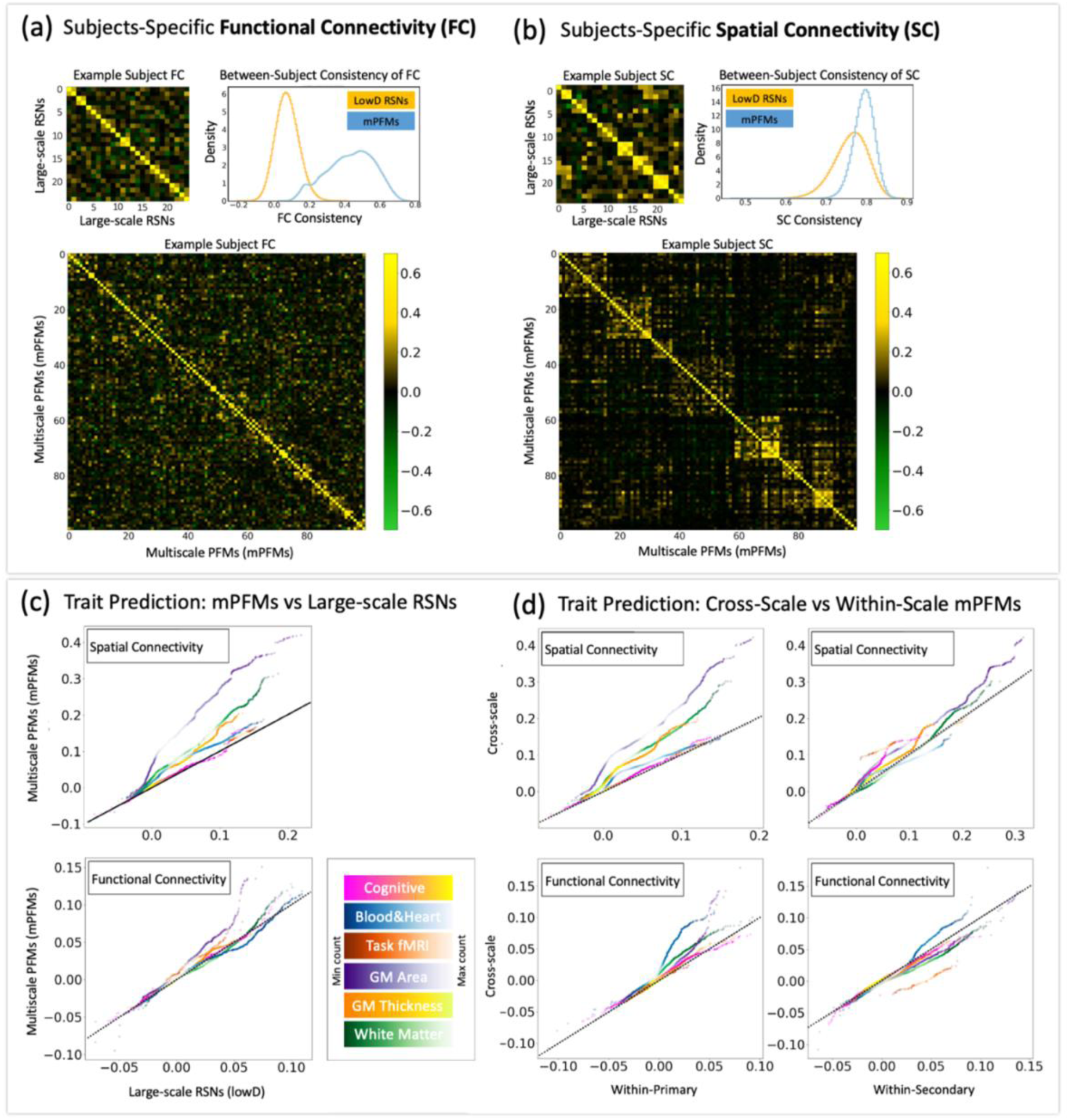
Spatiotemporal interaction of mPFMs captures subject variability that is predictive of personalised traits in UK Biobank. a) Examples of temporal correlations (functional connectivity) between the modes, which were used as one set of features for trait prediction. Functional connectivity matrices are shown separately for conventional large-scale RSNs from lowD (25 mode) decomposition and mPFMs (highD, 100 mode) decomposition. Histograms show that between-subject consistency of functional connectivity was notably higher in mPFMs. b) Examples of spatial connectivity between the modes which were used as another set of features for trait prediction. Spatial correlation matrices are shown separately for conventional large-scale RSNs from (lowD, 25 mode) decomposition and mPFMs (highD, 100 mode) decomposition. Histograms show that between-subject consistency of spatial connectivity was notably higher in mPFMs. c) Comparing the prediction performance of mPFMs with large-scale RSNs for a range of 893 phenotypes across 6 categories. Spatial (top) and temporal (bottom) correlations were used as predictors. d) Cross-scale correlations refer to spatial and temporal interactions between primary and secondary mPFMs, an aspect of the brain’s functional connectivity that conventional distributed-only RSNs or localised-only parcellations cannot capture. The prediction performance of this cross-scale spatial (top) and temporal (bottom) correlations were compared to that of within primary or within secondary modes, and a clear improvement was observed, especially in spatial domain. Density scatter plots are used to compare prediction performances, and colours denote densities, as shown in colourbars. Dotted black lines denote equal performance.

First, we compared the prediction performance of mPFMs with large-scale RSNs, shown in Figure 4c. We found that spatial connectivity of mPFMs, after matching the number of features as elaborated in 2.6.4, significantly outperformed (Bonferroni corrected, Table S 5) that of large-scale decomposition in predicting all phenotype categories except cognitive scores. Similarly, functional connectivity of mPFMs significantly outperformed that of large-scale decomposition in predicting region-wise cortical area and task fMRI phenotypes. This shows that subject-specific spatiotemporal correlations between mPFMs carry additional information about each person’s traits that is otherwise missed in large-scale decomposition.

Next, we divided spatiotemporal interactions of mPFMs into three subtypes: i) within primary mPFMs; ii) within secondary mPFMs, and iii) between primary and secondary mPFMs. Aiming to evaluate the added benefit of cross-scale interactions (iii), we compared its prediction performance to within-scale interactions (i.e., i&ii). As shown in Figure 4d-top, cross-scale spatial connectivity significantly outperformed (Bonferroni corrected, Table S 5) within-secondary for region-wise cortical area and thickness, task fMRI and cognitive phenotype categories, and they additionally outperformed within-primary for all the phenotype categories, except cognitive scores. Cross-scale functional connectivity, as shown in Figure 4d-bottom, performed on par with or worse than within-secondary, but significantly outperformed (Bonferroni corrected, Table S 5) within-primary for region-wise cortical area, thickness, cardiovascular health metrics.

These results demonstrate that spatial and temporal connectivity of the ensemble of mPFMs yields improved biomarkers for individualistic traits in two ways: first, compared with conventional large-scale RSNs, and second, subject variability captured in spatiotemporal correlations *across* multiple scales is more phenotypically relevant compared with within a single scale.

### 3.2 Validation of mPFMs

Our results so far have focussed on resolving the origins and functional relevance of mPFMs in modelling brain connectivity. Here we set to validate this new decomposition based on simulations, its reproducibility between HCP and UK Biobank datasets and similarity to HCP’s Multimodal Parcellation.

#### 3.2.1 Simulations

Based on real data in section 3.1, we found evidence that secondary mPFMs likely emerge to explain temporally-distinct subcomponents within the conventional large-scale RSNs. In other words, we showed that the ensemble of multiscale modes captures the integrated-segregated functional brain connectivity. Here we simulated datasets, where distributed and localised modes co-existed, such that localised modes were spatial sub-modes of the distributed modes, with distinct timecourses. Using these simulations, we tested how well PFM decomposition can recover the co-existence of multiscale modes, and compared its performance to spatial ICA, which is the current standard technique for RSN modelling. Across 10 simulated datasets, each included 12 modes, 6 distributed and 6 localised. Figure 5a shows examples of group-average simulated modes in one of the datasets. We compared three elements of the estimated modes with the ground truth: group spatial maps, subject spatial maps, and subject time courses, shown in Figure 5 b, c, d, respectively. For the group spatial maps, the accuracy of PFM decomposition for distributed and localised modes was 0.93 and 0.78, respectively, which was reduced to 0.74 and 0.47 in spatial ICA modes. For the subject spatial maps, the accuracy of PFM decomposition for distributed and localised modes was 0.87 and 0.70, respectively, which was nearly halved (0.48 and 0.31) in spatial ICA modes. For the subject timecourses, the accuracy of PFM decomposition for distributed and localised modes was 0.95 and 0.64, respectively, which was reduced to 0.78 and 0.33 in spatial ICA modes. Therefore, at both group and subject level, PFMs can recover the simulated multiscale modes with overall (average) accuracy of ∼0.8 or higher.

**Figure 5.**
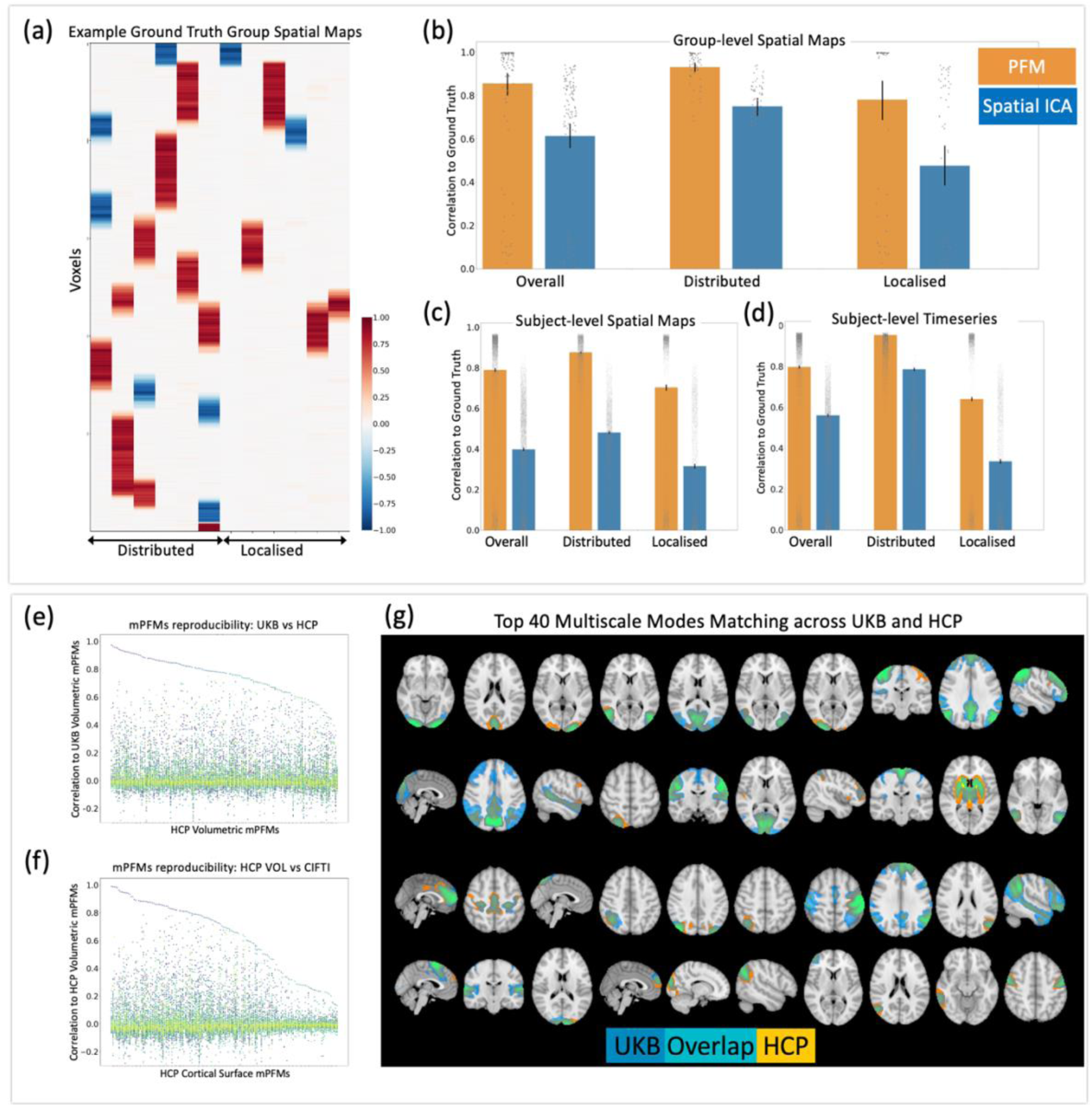
Validation of mPFMs using synthetic data (top panel) and reproducibility in real data (bottom panel) Simulations: a) Examples of simulated group-average spatial maps (one of ten simulated datasets). 12 modes were simulated, 6 mimicking distributed modes, 6 mimicking localised subcomponents of these modes. Aiming to validate findings of section 3.1.1, we observed that PFM decomposition can well capture: b) group spatial maps; c) subject spatial maps; d) subject timecourses of these co-existing modes. The performance was ∼50% higher than that of spatial ICA at group level and up to 100% higher at subject level. Real data reproducibility: PFM decomposition with 150 modes was compared at the group level between: e) UK Biobank (UKB) and HCP volumetric fMRI; each HCP mode is shown on the x axis and correlations to all UKB modes on the y axis. Clear one -to-one pairing can be seen for the top 100 modes. f) A similar comparison was conducted between HCP volumetric and surface-based fMRI data types. g) The top 40 matching mPFMs between HCP and UKB (both volumetric fMRI) are illlustrated. HCP modes are shown in yellow, UKB in blue and overlap in green/cyan.

#### 3.2.2 Reproducibility across datasets and data formats

The next step of validation involved cross-dataset reproducibility tests. We applied sPROFUMO separately to three rfMRI datasets and obtained 150 modes per run: 1) 1003 HCP subjects, volumetric whole-brain rfMRI (HCP VOL); 2) 1003 HCP subjects, CIFTI cortical rfMRI (HCP CORT); 3) 4999 UKB subjects, volumetric whole-brain rfMRI (UKB VOL). All these scenarios yielded mPFMs consisting of primary and secondary modes. We paired these mPFMs across runs based on spatial correlation of group-level spatial maps.

Firstly, by comparing HCP VOL and UKB VOL (Figure 5e), we found 100 modes to show very good matching (absolute correlation >= 0.7), 40 modes to show good matching (absolute correlation between 0.5 and 0.7), and 10 modes to show mediocre matching (absolute correlation <= 0.5). Some of the modes in the latter category were due to differences in structured noise across datasets (Figure S 8). The top 40 best matching mPFMs across these datasets are shown in Figure 5g.

Secondly, by comparing HCP VOL and HCP CORT (Figure 5f), we found 74 modes to show very good matching (absolute correlation >= 0.7), 28 modes to show good matching (absolute correlation between 0.5 and 0.7), and 48 modes to show mediocre matching (absolute correlation <= 0.5). Some of the modes in the latter category were due to structured noise differences across data-types, and the remainder were due to inherent differences between volumetric and CIFTI fMRI reconstructions.

##### 3.2.2.1 Reproducibility across multiple dimensionalities

We additionally tested for reproducibility of the mPFMs across multiple dimensionalities: 50, 100 and 150. This was done separately for UK Biobank and HCP datasets, with each dimensionality starting from a different independent initialisation. Results are shown in **Figure S 9**, starting from low-dimensional d=25 decomposition, we compared 25 versus 50, 50 versus 100 and 100 versus 150. The modes were sorted and paired based on spatial correlations between the group-level spatial maps. For the UK Biobank volumetric fMRI, we found mean±STD of the paired correlations to be: 0.90 ± 0.11 (d = 25 vs 50), 0.84 ± 0.14 (d = 50 vs 100), 0.82 ± 0.14 (d = 100 vs 150). For the HCP Cortical CIFTI fMRI we found mean±STD of the paired correlations to be: 0.88 ± 0.08 (d = 25 vs 50), 0.90 ± 0.11 (d = 50 vs 100), 0.88 ± 0.11 (d = 100 vs 150). These strong pairings show that, as we increase the dimensionality, the existing modes at lower dimensions are largely maintained, and new modes are added to the ensemble, indicating an eventual addition of more and more secondary PFMs.

#### 3.2.3 Validating secondary mPFMs using HCP’s Multi-Modal Parcellation

As the next step, we validated mPFMs based on the HCP’s Multimodal Parcellation v1.0 (HCP-MMP), aiming to validate the secondary mPFMs. We compared mPFMs against the HCP-MMP, which includes 360 cortical parcels. This parcellation includes both group-average and subject-specific parcels, allowing us to validate secondary mPFMs at both levels. We applied sPROFUMO to cortical rfMRI data of 446 HCP subjects (matching the number of subjects in HCP-MMP) and estimated 360 modes. Similar to before, this yielded a multiscale ensemble of modes (i.e., mPFMs).

We found an interesting group correspondence (Figure S 10a) between secondary mPFMs and HCP-MMP, especially 95 parcel-like mPFMs showed a strong correspondence to HCP-MMP (dice similarity >=0.5), illustrated in Figure S 10c. 73 mPFMs showed less than 0.2 dice similarity to any HCP-MMP parcels; these included conventional low-dimensional modes and the less well-known variants of distributed modes, and typically occupied multiple distant sub-regions. We further found that modes with high group-level correspondence also showed high subject-level correspondence (Figure S 10b), with the top 95 matching modes at the group-level also showing correlation of 0.47±0.19 (average ± standard deviation across subjects and modes) at the subject-level. These evaluations add to the previous validation steps, by demonstrating that those of the mPFMs that are parcel-like have clear spatial similarities to the state-of-the-art individualised parcellations, both at the group- and the subject-level.

In summary, this section on mPFM validation sheds light on robustness and interpretation of the multiscale representation of the RSNs. Specifically, in section 3.2.1, we simulated scenarios resembling multiscale modes; i.e., temporally distinct pairs of modes where one mode is a spatial sub-node of the other, thus mimicking the primary and secondary modes in real data. We showed that these can be recovered using PFM decomposition much more accurately than ICA decomposition. Next, in section 3.2.2, we demonstrated reproducibility of mPFMs across two large-scale datasets (HCP and UKB), two standard fMRI data formats (volumetric and CIFTI), and a range of dimensionalities (25, 50, 100, 150) in both HCP and UKB. These results demonstrate that the multiscale representation is a stable finding and not a byproduct of certain data properties or choices of preprocessing or dimensionality. Finally, in section 3.2.3, we demonstrated that secondary mPFMs can be reliably linked to the well-established HCP-MMP parcellation. Considering that the multiscale representation is (as shown in earlier in Figure 1, and section 3.2.2.1), in principle, an ensemble of the large-scale RSNs (primary mPFMs), with the addition of the fine-grained secondary mPFMs, this analysis validated that the secondary mPFMs (especially the parcel-like) are meaningful functional entities, and not due to fMRI smoothness or noise properties.

### 3.3 mPFMs yield enhanced biomarkers of personalised traits

We have so far investigated the brain properties underlying mPFMs, and their added benefit over conventional large-scale RSNs. Here we focus on the usefulness of individual-specific mPFMs for providing biomarkers for traits and disease. This was done using two steps: 1-population covariations between mPFMs and behavioural traits in HCP; 2-out of sample prediction of phenotypes in UK Biobank. CCA and phenotype prediction results serve two distinct, complementary goals. mPFMs provide a new type of RSN representation, which is subject-specific and multiscale. Therefore, we aimed to test: 1-how interpretably they can be linked to non-imaging variables (CCA analysis in HCP), and 2-compared to existing techniques, how accurately they can predict non-imaging variables (phenotype prediction in UK Biobank).

#### 3.3.1 Positive-negative axis of population covariation between mPFMs and behaviour

The goal of CCA analysis was mainly qualitative, and to illustrate how interpretably mPFMs can be linked to personalised traits. In order to achieve this, we referred to the previous literature and investigated whether mPFMs can capture one of the key findings in brain-behaviour literature: a positive-negative mode of population covariation that has been discovered to link brain features to cognition and lifestyle. Positive traits such as language comprehension or life satisfaction are placed at one end of this axis, and negative traits such as antisocial behaviour or substance use at the other end. This was originally reported by Smith et al., (2015) using HCP data, and has been replicated on other large datasets since (Goyal et al., 2022).

Canonical Correlation Analysis (CCA), allows to investigate associations between a set of features from brain data and a set of personalised traits (i.e., phenotypes), all in a single integrated multivariate analysis. Using CCA, we identified modes of population co-variation between brain and phenotypes, as pairs of canonical components along which the phenotypes and mPFMs co-varied in a similar way across subjects. We used the same analysis pipeline as Smith et al., (2015), as detailed in section 2.7.

To examine the link between personalised traits and spatiotemporal characteristics of mPFMs, CCA was conducted on features related to spatial maps, spatial and functional connectivity of mPFMs. Figure S 11a shows results of correlation between brain and phenotypes after CCA transformation. As shown in Figure 6a, we identified the raw behavioural metrics that were most strongly linked to the top CCA components. This was done by correlating the behavioural metrics with the topmost CCA components related to spatial maps, spatial and functional connectivity, and taking the average correlation across the three. This unravelled a single axis of population variation: at one end of this axis, we found behavioural traits positively associated with the prominent CCA component. These included metrics related to cognition and emotion such as performance on language tests, executive function, self-regulation, and life satisfaction. At the other end of this axis, we found behavioural traits negatively associated with the prominent CCA component. These included metrics related to psychiatric disorders and substance use such as rule breaking, antisocial behaviour, alcohol, tobacco and cannabis use. Therefore, mPFMs, similar to existing literature, provide a simple interpretable mapping between brain features and positive versus negative traits.

**Figure 6.**
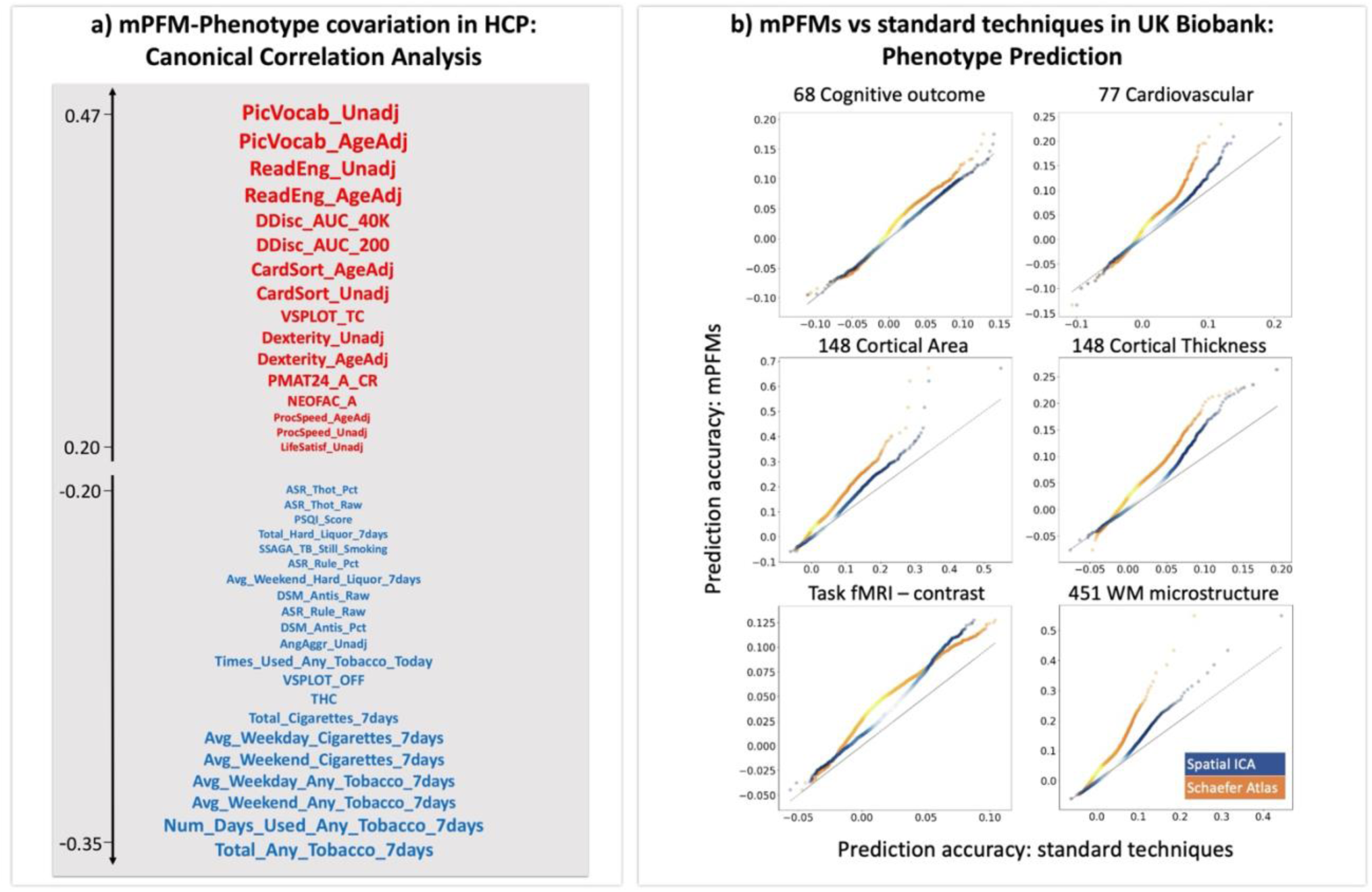
mPFMs yield biomarkers for behaviour and traits. a) shared population covariation between mPFMs and behaviour in HCP: A positive-negative axis of population covariation identified based on top CCA components of spatial and temporal mPFMs features. b) mPFMs yield enhanced fMRI biomarkers of traits and health compared with current standard techniques in UKB. 893 phenotypes across six categories were used. Comparisons between trait prediction accuracies of mPFMs (y-axis) with spatial ICA and the Schaefer parcellation (x-axis blue and orange, respectively) are shown. Correlation coefficient between actual and predicted phenotypes in the test set was used as metric of accuracy. Density scatter plots are used to compare mPFM’s prediction accuracies to that of the standard techniques, each dot corresponds to the prediction accuracy of a mode for a phenotype/trait, and brighter colours (e.g. light yellow and light blue) denote higher densities. Dotted grey lines denote equal performance. 100-mode decompositions were used for all the methods, and prediction accuracies were computed per mode, in order to obtain a more detailed summary of the modes. mPFMs outperformed the standard techniques for all the categories.

#### 3.3.2 Enhanced predictions of personalised traits in UK Biobank

Next, we evaluated the potential of mPFMs for out-of-sample prediction of traits. For this purpose, we compared mPFMs to two of the most commonly used techniques in the literature: spatial ICA, the current main RSN modelling technique in the core UK Biobank pipeline (Alfaro-Almagro et al., 2018); and the Schaefer atlas, a high dimensional hard parcellation derived from the Yeo parcellation (Schaefer et al., 2018). For this purpose, we used the 100-dimensional Schaefer parcellation and further applied the other two methods to volumetric rfMRI data of 4999 UK Biobank subjects, estimating 100 mPFMs and 100 ICs. We used features of each functional mode (feature matrix, X) to make predictions about phenotypes (target, y), details in 2.6. We used ElasticNet regression with 5-fold nested cross-validation, and predicted 893 phenotypes divided into 6 categories: region-wise cortical area and thickness (296), White Matter (WM) tracts’ microstructure (451), task fMRI response to the contrast of emotional faces vs shapes (1), cognition (68) and cardiovascular health metrics (77). Correlation coefficient between actual and predicted phenotypes in the test sets was used as metric of accuracy. Results are shown in Figure 6b, with prediction accuracies of mPFMs shown on the y-axis, and accuracies of spatial ICA modes on the x-axis. We found that mPFMs significantly outperformed (Bonferroni corrected, Table S 6) Schaefer parcellation in prediction of all the phenotype categories, and significantly outperformed spatial ICA in prediction of all the phenotype categories except cognitive traits. These results were replicated using higher dimensionalities, as shown in **Figure S 12**.

These results, taken together with the findings of section 3.1.5, demonstrate three key benefits of mPFMs over single-scale representations for phenotype predictions: firstly, the spatiotemporal connectivity between the large-scale and fine-grained PFMs (i.e., cross-scale connectivity) outperforms that of within-scale connectivity -this was illustrated using the PFM modelling framework only, since standard techniques do not yield multiscale representations; secondly, the spatiotemporal connectivity of the full multiscale representation outperforms that of lowD large-scale-only RSNs; thirdly, the full set of features obtained from the multiscale highD representation (including spatial topographies and spatiotemporal connectivity) outperforms that of standard single-scale highD representations obtained from ICA and Schaefer atlas.

## 4 Discussion

In this paper, we presented Multiscale Probabilistic Functional Modes (mPFMs), a new representation of the brain’s functional modes, that improves fMRI modelling and its utility for prediction of cognition and health. mPFMs were identified using stochastic Probabilistic Functional Mode (sPROFUMO) modelling and, unlike any existing representation, include an ensemble of modes across multiple scales. sPROFUMO does not impose or prevent this behaviour by design, instead, it allows the modes to be spatiotemporally correlated, and accumulate Bayesian evidence across individuals in large populations. Therefore, if a multiscale organisation arises from the results, this is driven by the structure in the data. Indeed, as demonstrated in the Results section, the presence of multiple distinct subcomponents within each large-scale (distributed) mode indicates spatiotemporal multiplicity within these modes, which cannot be captured by distributed-only or localised-only representations. Put together, our results demonstrate mPFMs’ potential to provide a useful representation of multiscale information processing in the brain, and enhance fMRI-derived biomarkers for traits and disease. We release this representation publicly; we have made the group-level maps openly available, whereas subject-level maps will be released in the future as part of ongoing work to rebuild a “v2” UKB pipeline (Alfaro-Almagro et al., 2018).

Multiscale modes are compatible with the notion of hierarchal information processing in the brain; that brain function is organised in a hierarchical manner, with each level of the hierarchy underlying increasingly complex and abstract processing (Cordes et al., 2002; Deco and Kringelbach, 2017; Friston, 2008; Vidaurre et al., 2017). Hierarchical processing is enabled via interactions between local unimodal information processing and distributed higher level cognitive function. Such interactions have been well-established, e.g., in visual search tasks, where interactions between sub-regions of the visual cortex and fronto-parietal attention networks allows for top-down modulation of visual processing by attention, as well as bottom-up modulation of attention by the incoming stimulus(Buschman and Miller, 2007; Gazzaley and Nobre, 2012). Interactions between brain systems at different levels of these hierarchies cannot be directly modelled using existing functional mode representations, considering that they can either be used to measure interactions between distributed modes (lowD decompositions) or localised regions (highD parcellations). As a result, the interactions *between* these two layers will have to be modelled using post-hoc techniques, such as hierarchical clustering (Smith et al., 2013a) or module detection(Sporns and Betzel, 2016). Even then, the resulting connectivity will depend on the assumptions of the post-hoc techniques, and change depending on the choice of parameters, rather than being integrated within the original brain function mapping.

In contrast, mPFM inherently incorporates such multiscale connectivity within fMRI decomposition. Crucially, as we demonstrated in the Results, these interactions provide stronger biomarkers of cognition and health compared with connectivity within large-scale or within localised modes alone. This complements previous brain-behaviour research (Bolt et al., 2018), that compared the association of individual brain regions and their respective whole-brain networks with task performance. The authors showed that for tasks such as working memory and relational, areal-level associations to task performance will become redundant when large-scale network covariation in accounted for, unlike arithmetic task, where areal-level and network-level associations showed distinctively significant roles. This study, similar to ours, highlights the need for multiscale representations.

Additionally, by allowing the modes to be spatially overlapping, mPFMs can capture functional multiplicity in the brain’s organisation, where a single region contributes to different functions depending on the context. Until recently, spatially overlapping functional modes have been largely absent from resting state literature. Specifically, studies have commonly relied on either spatial ICA (Beckmann et al., 2005; Calhoun and Adali, 2012), that imposes spatial independence between the modes, or hard parcellations, that enforce rigid boundaries between the modes (Craddock et al., 2012; Yeo et al., 2011). Therefore, the literature has relied on spatial bases that minimise or prevent spatial overlap. However, growing evidence is now highlighting the importance of spatial overlap in modelling cross-individual variability in brain function (Bijsterbosch et al., 2019). Importantly, recent successful characterisation of brain connectivity using gradients or connectopy mapping (Haak et al., 2018; Margulies et al., 2016) has brought functional multiplicity to the fore of resting state connectivity. These techniques have unravelled multiple overlapping connectivity patterns within brain regions, e.g., retinotopic organisation of the visual cortex, and follow-up studies have highlighted the cognitive and clinical relevance of these overlapping connectopy maps. As such, allowing for spatial overlap, in addition to and not instead of temporal correlations between the modes, is proving imperative for obtaining a more comprehensive representation of brain function. mPFMs are, on the one hand, similar to conventional spatial ICA modes, in that they capture large-scale prominent networks in the brain. One the other hand, mPFMs additionally allow multiple overlapping subcomponents within these networks to co-exist. As such, mPFMs can be viewed as a new category of brain function mapping that stands somewhere in-between conventional RSNs and the more recent gradients of connectivity. In this context, future studies would be valuable, to compare mPFMs to gradients of connectivity. Additionally, comparison to the recently-introduced tensor-based decomposition methods such as NASCAR (Li et al., 2023) and Tucker (Han et al., 2024) would be valuable. Unlike spatial/temporal ICA, and similar to PROFUMO, these methods allow the estimation of RSNs that are both spatially and temporally correlated.

In addition to their functional relevance, mPFMs also provided enhanced biomarkers for prediction of individualistic traits in UK Biobank. To benchmark of mPFMs against standard ICA-based or hard parcellation techniques, we used a range of imaging-derived and non-imaging derived phenotypes such as cortical geometry, white matter microstructure, brain response to cognitive tasks, behavioural cognitive scores, and cardiovascular health outcome. The overall prediction accuracies of mPFMs were found to be significantly higher than standard techniques for various phenotype categories. One key reason for this higher prediction accuracy is likely the more comprehensive spatiotemporal feature set in mPFMs. On the one hand, hard parcellations yield binary spatial maps for each mode/parcel, and no spatial overlaps between the modes, thus summarising individual-specific brain features solely in temporal space. Somewhat similarly, Spatial ICA, due to its assumption of spatial independence between the modes, minimises spatial overlap by design (Farahibozorg et al., 2021; Kiviniemi et al., 2009; Smith et al., 2013b). Consequently, both approaches yield singlescale decompositions of functional modes. Therefore, large-scale modes, and their spatiotemporal correlation with each other and with localised modes, cannot be fully characterised by the standard techniques. Therefore, mPFMs yield a richer set of spatial and temporal features that, even after matching the number of features between the methods, can outperform the standard techniques in reflecting individualistic traits in brain function.

## 5 Acknowledgements

We would like to thank Dr. Matthew F Glasser for providing subject-specific HCP-MMP (v1.0) parcellations and helpful input on an earlier version of the manuscript. We are grateful to HCP and UK Biobank for making these invaluable resources available, and to the UK Biobank and HCP participants for dedicating their time to make these data possible.

## Funding

- Royal Academy of Engineering under the Research Fellowship programme RF2122-21-310 (SRF)
- Wellcome Trust grants 215573/Z/19/Z (SMS and MWW), 106183/Z/14/Z (MWW). Wellcome Centre for Integrative Neuroimaging is supported by core funding from the Wellcome Trust (203139/Z/16/Z).
- MRC Mental Health Pathfinder grant MC_PC_17215 (SMS)
- National Institute of Health grants R01 MH128286 and R01 MH132962 (JDB)
- The computational aspects of this research were partly carried out at Oxford Biomedical Research Computing (BMRC), that is funded by the NIHR Oxford BRC with additional support from the Wellcome Trust Core Award Grant Number 203141/Z/16/Z. The views expressed are those of the author(s) and not necessarily those of the NHS, the NIHR or the Department of Health.

For the purpose of open access, the author has applied a CC BY public copyright license to any Author Accepted Manuscript version arising from this submission.

## Author contribution

Conceptualisation: SRF, SJH, MWW, SMS; Methodology: SRF, SJH, JDB, MWW, SMS; Investigation: SRF, MWW, SMS, Software: SRF, SJH, SMS; Validation: SRF; Data curation, Formal Analysis and Visualisation: SRF; Resources: MWW, SMS; Funding acquisition: SRF, MWW, SMS; Writing—original draft: SRF; Writing—review & editing: SRF, SJH, JDB, MWW, SMS.

## Competing interests

Authors declare that they have no competing interests related to this manuscript. At the time of contributing to this research, SJH was affiliated with Universities of Oxford & Otago, he is currently employed by ADInstruments.

## Data availability

- UK Biobank data is available upon registration and applying for data access from UK Biobank website: http://www.ukbiobank.ac.uk/register-apply.
- Human Connectome Project data is available upon registration from HCP website: https://db.humanconnectome.org.
- Group average mPFMs obtained from HCP and UK Biobank are available here: https://www.fmrib.ox.ac.uk/ukbiobank/PROFUMO.

## Code availability

- Code for sPROFUMO is currently available from the following repository, it will be made available in an upcoming FSL release: https://git.fmrib.ox.ac.uk/profumo/profumo/-/tree/sprofumo-cpp-clean.
- Code used for simulations is available from the following public repository: https://git.fmrib.ox.ac.uk/profumo/PFM_Simulations.

## 7 Supplementary Materials

**Figure S 1.**
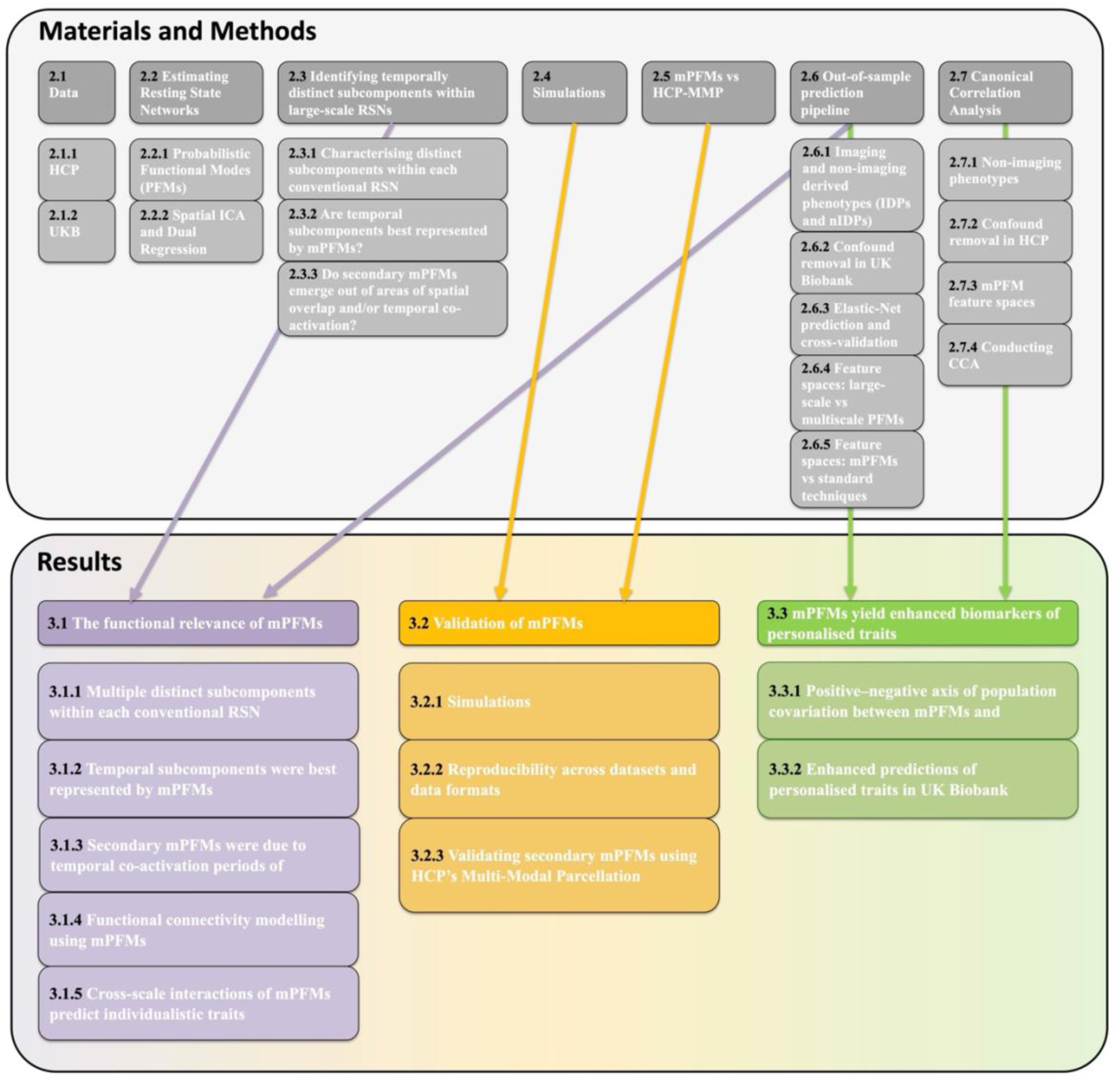
A Flowchart of different subsections of Methods and Results as well as the key analysis steps feeding into each Results subsection. Note that since Data and Estimating RSNs feed into multiple Results sections, they are not linked via arrows to a void unnecessary business of the graph.

**Figure S 2.**
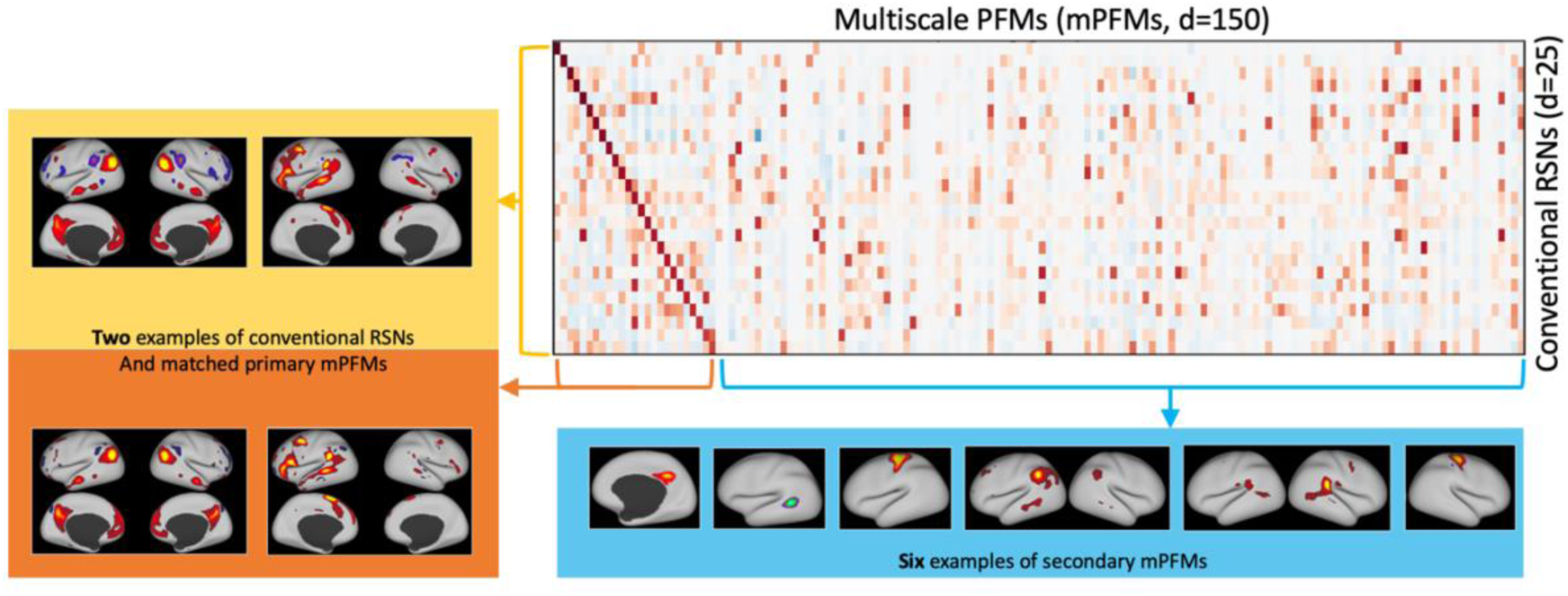
Supplement to Figure 1: This figure provides an alternative visualisation of the comparison between multiscale Probabilistic Functional Modes (mPFMs) and conventional large-scale RSNs. This visualisation provides additional clarification that when mPFMs are spatially paired with large-scale RSNs from 25-mode decomposition (yellow), 25 of the mPFMs show a clear one-to-one matching, labelled as Primary mPFMs (orange). Secondary mPFMs (blue) are the remaining 125 mPFMs, that start appearing with increased dimensionality.

**Figure S 3.**
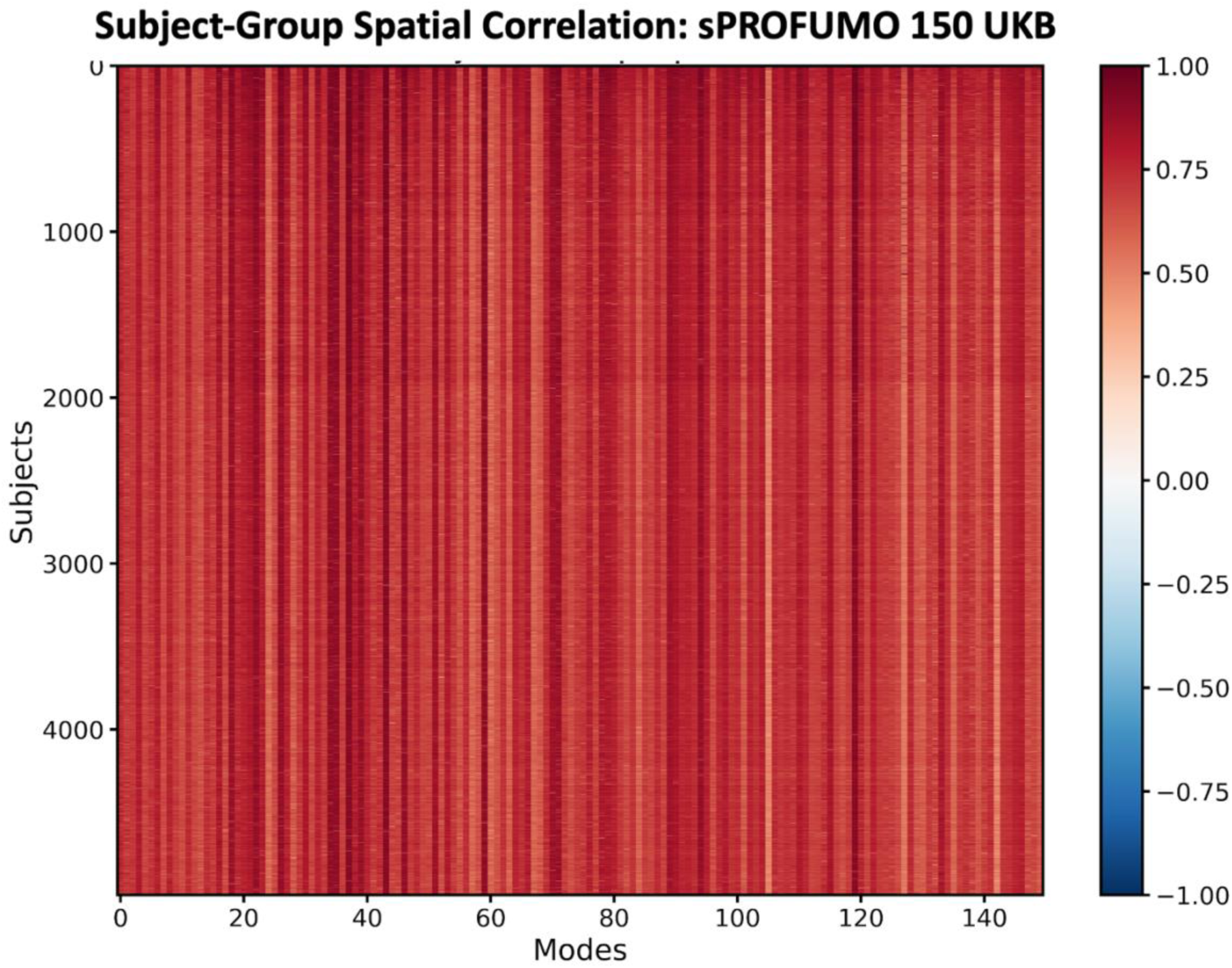
An example of Subject-group similarity of spatial maps obtained from 150-PFM decomposition of rfMRI data from 5000 subjects in UK Biobank (UKB) This can be used to examine which subject-PFM pairs have been reliably estimated. Bad subject-PFMs can be identified using two heuristic criteria: firstly, if the spatial correlation to the group is very small (<0.1), this indicates a very noisy estimation of that specific PFM for that subject – which would be worth considering removal. Secondly, if the spatial correlation is artefactually high (>0.99), this indicates that posteriors were not reliably estimable at subject level, and therefore that subject-PFM was reverted to group priors – such modes should be removed. In general, in the example shown here, all the modes were estimated reliably for all the subjects. It is worth noting that some PFMs are generally of higher SNR (e.g., Visual or Motor), and they will be more consistent across subjects (columns with darker red colours), whereas other lower-SNR PFMs will be less consistent across subjects (columns with lighter red colours).

**Figure S 4.**
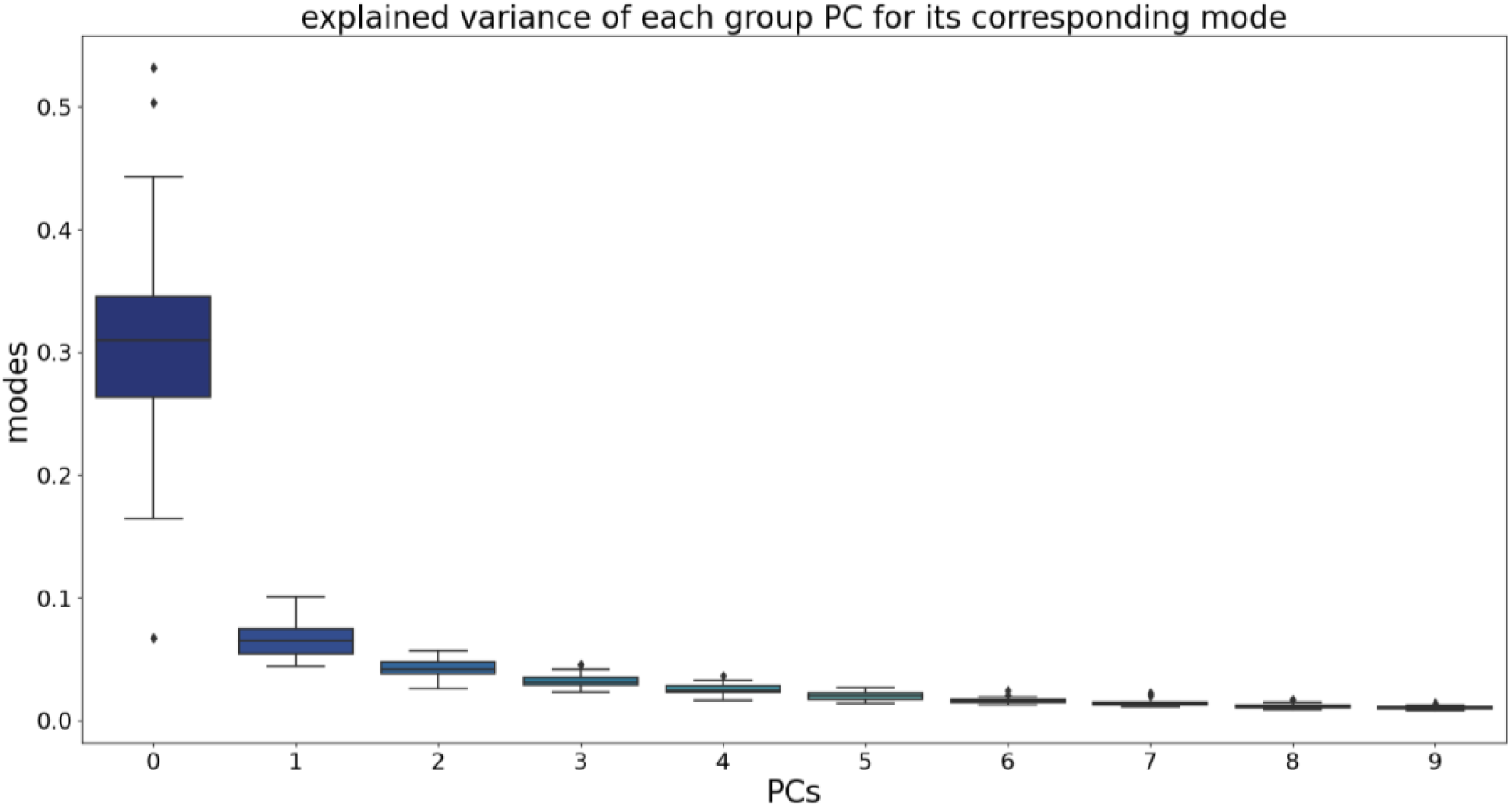
Supplement to Section 3.1.1. Principal Component Analysis was used to test if one temporal component is sufficient to capture temporal variability within conventional large-scale RSNs from lowD decompositions of rfMRI. LowD refers to 25 modes obtained from PFM decomposition of rfMRI in HCP. The top 10 PCs per lowD mode are illustrated on the x-axis, and their explained variance on the y-axis. Boxplots show median and confidence intervals across 25 modes.

**Figure S 5.**
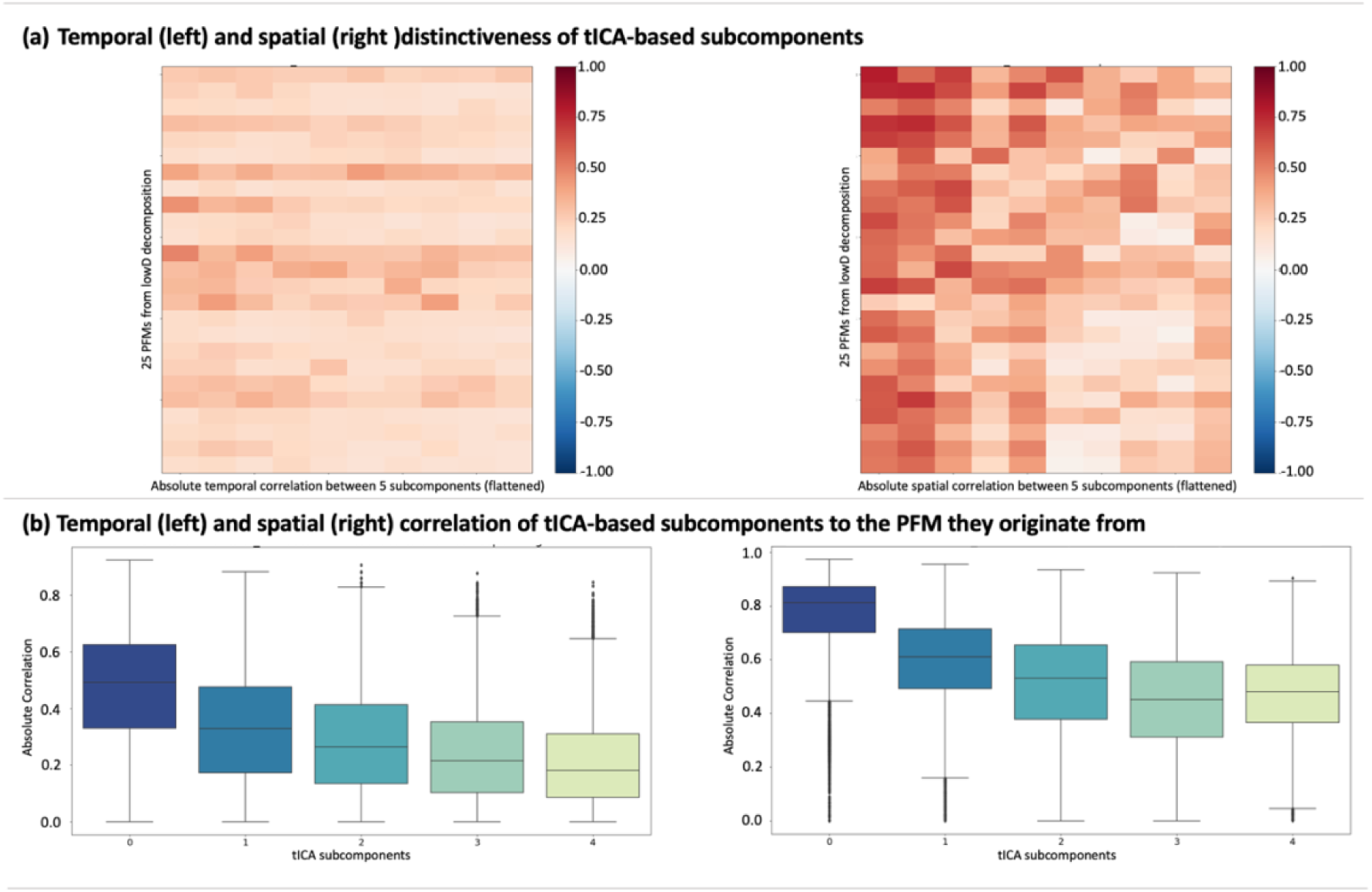
Supplement to section 3.1.1. a) The degree to which 5×25 subcomponents estimated using temporal ICA are spatially (right) and temporally (left) correlated with each other (i.e. testing for distinctiveness). The 25 rows correspond to the 25 modes, and the flattened above-diagonal of 5×5 correlation matrices (correlations between subcomponents estimated within-subject and then averaged over subjects) appear on the x-axis. b) Temporal (left) and spatial (right) correlation of 5×25 subcomponents with their respective lowD modes. Boxplots show median and confidence intervals across 25 modes.

**Figure S 6.**
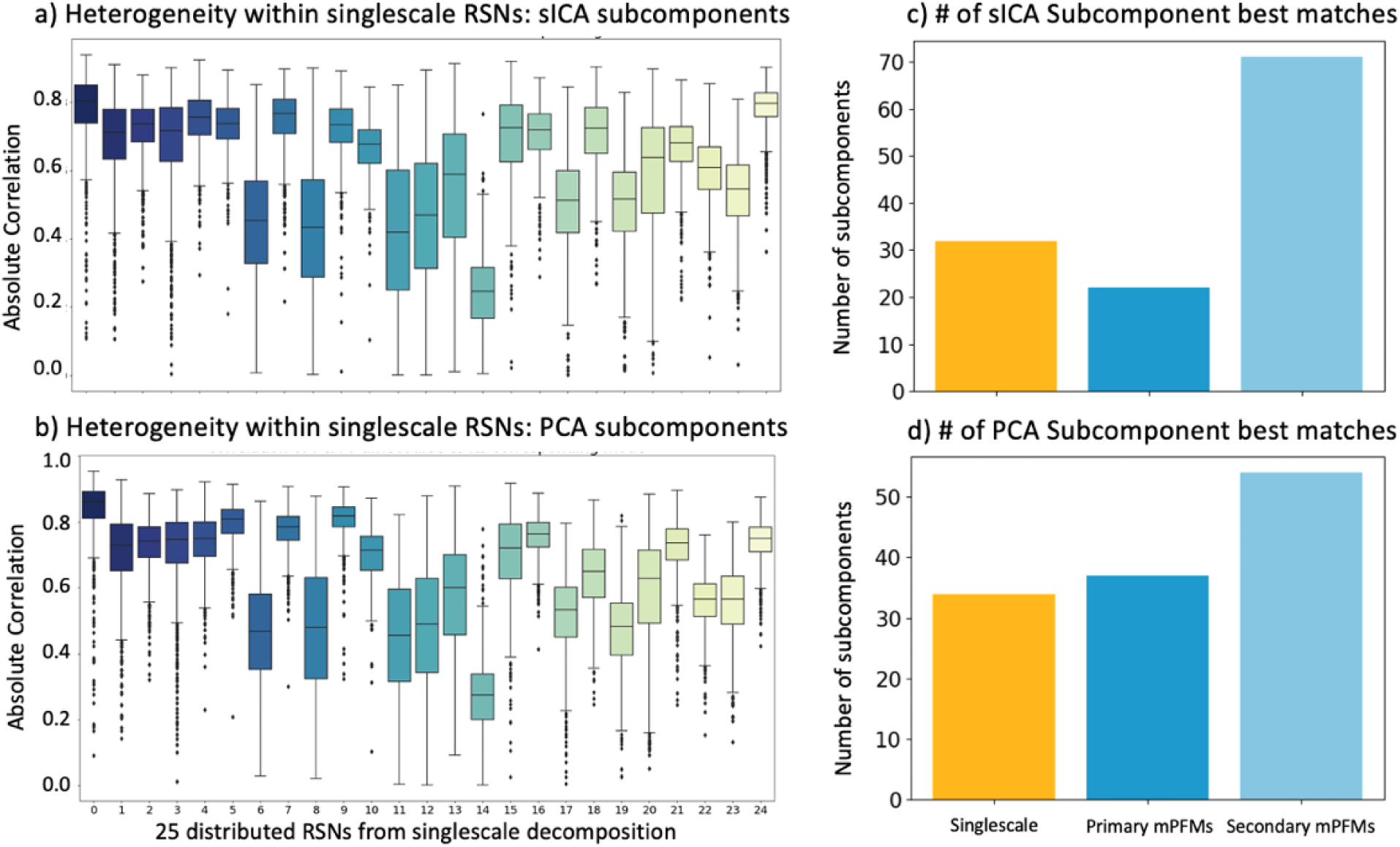
Supplement to Section 3.1.1. A low-dimensional (lowD) PFM decomposition consisting of 25 modes was estimated from cortical rfMRI data of 1003 HCP subjects, yielding conventional large-scale RSNs, and follow-up spatial ICA (top) and PCA (bottom) applied to voxel-wise timeseries within these RSNs to identify temporally-distinct subcomponents. This analysis was conducted to confirm that subcomponent identification results in section 3.1.1are not specific to the choice of temporal ICA as the subcomponent identification technique. Temporal correlation of the best-matching a) sICA and b) PCA subcomponents to the large-scale RSNs that they originated from; A winner-takes-all approach was applied: c) of the 25×5 sICA subcomponents 32, 22 and 71 were best represented by large-scale decomposition, primary and secondary mPFMs, respectively; d) of the 25×5 PCA subcomponents 34, 37 and 54 were best represented by large-scale decomposition, primary and secondary mPFMs, respectively.

**Figure S 7.**
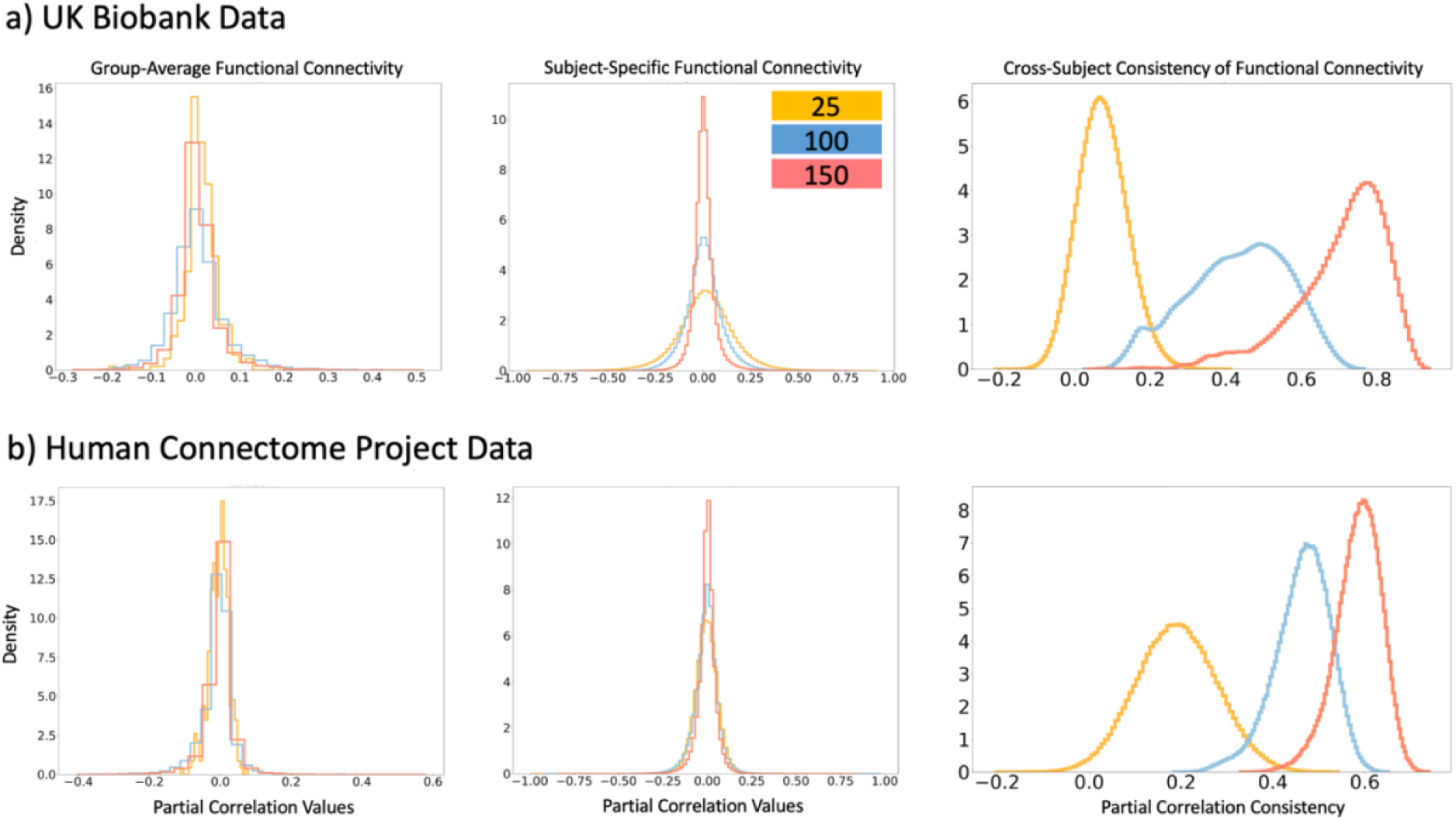
Supplement to section 3.1.4. Functional connectivity values between modes, estimated using regularised partial correlations. Three PFM dimensionalities are compared: 25, 100 and 150 based on a) 4999 subjects in UK Biobank data (volumetric fMRI) and b) 1003 subjects in Human Connectome Project data (cortical CIFTI). Left: distributions of group average Functional connectivity values. Middle: distributions of subject specific functional connectivity values. Right: Cross subject consistencies of Functional connectivity, which is calculated as Pearson correlation coefficient between vectorised functional connectivity matrices across subjects. Subject-specific functional connectivity becomes sparser and more consistent acorss subjects in mPFMs, leading to less sparse group-average functional connectivity.

**Figure S 8.**
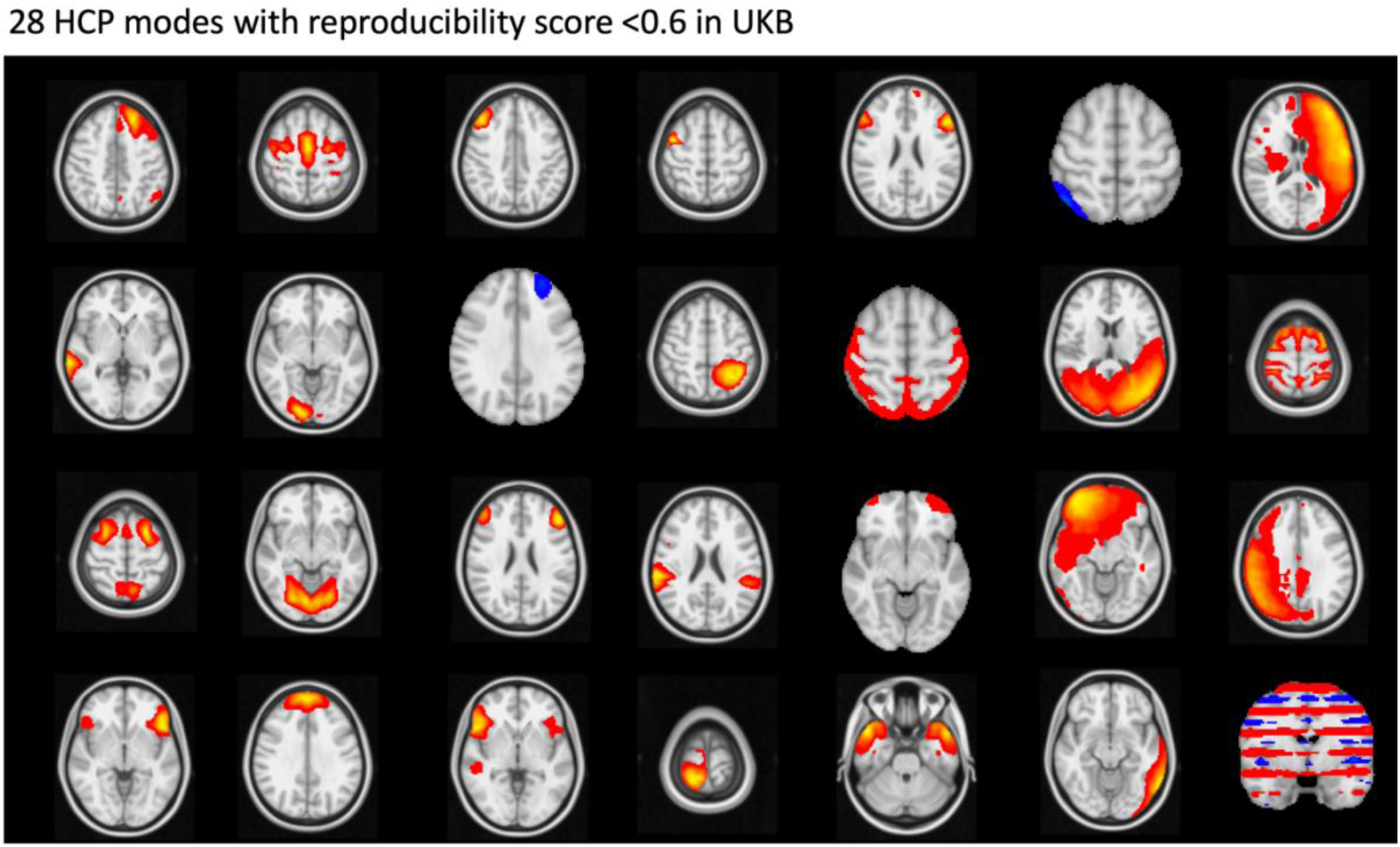
Supplement to Figure 5. 28 Volumetric HCP modes with reproducibility scores of <0.6 in UKB. Several of these modes were artefactual.

**Figure S 9.**
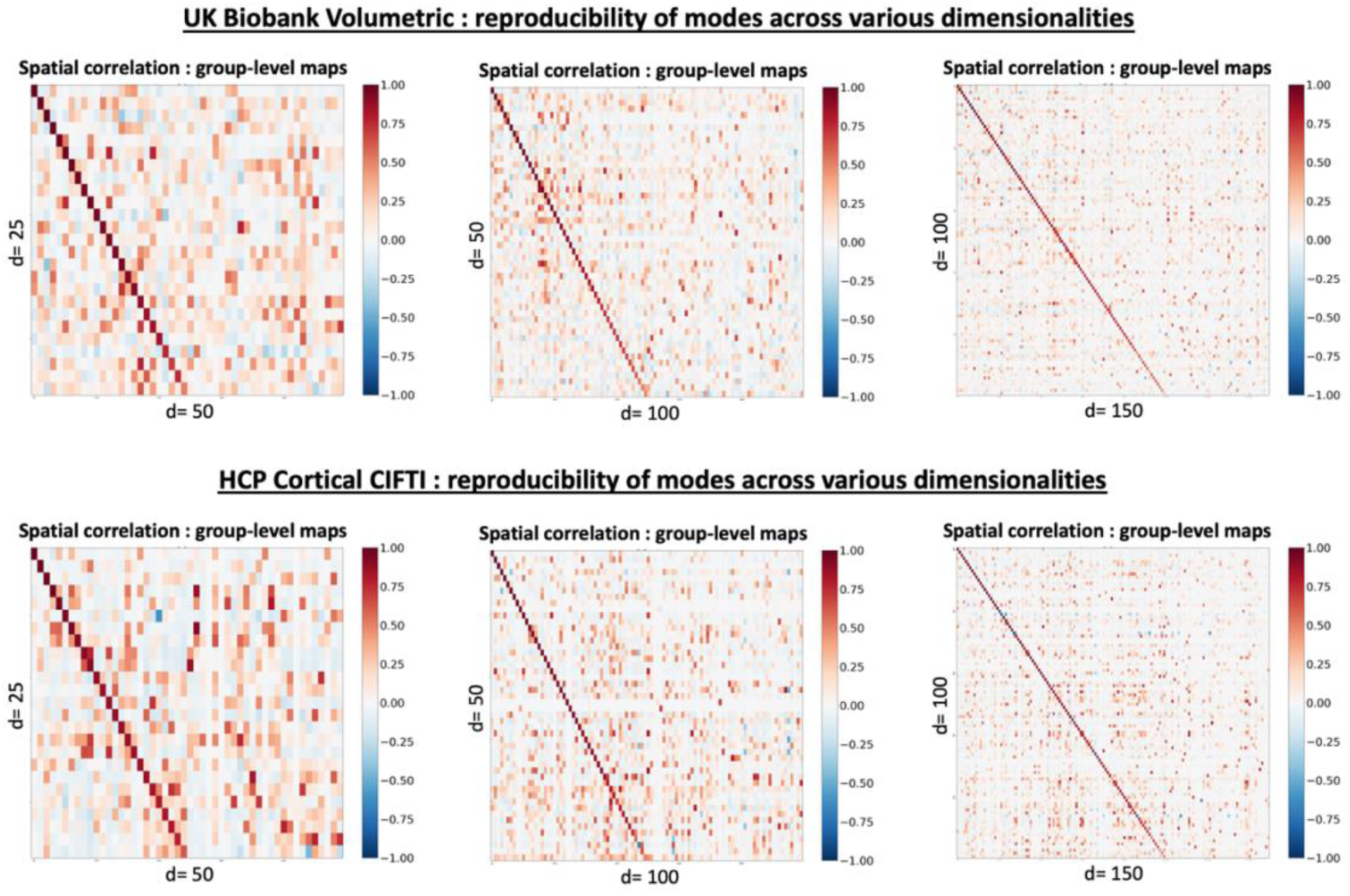
Supplement to section 3.2.2.1. Reproducibility of mPFMs across multiple dimensionalities in two independent datasets, UK Biobank and HCP starting from low-dimensional d=25 decomposition, we compared 25 vs 50, 50 vs 100 and 100 vs 150. The modes are sorted and paired based on spatial correlations between the group-level spatial maps. Strong diagonal elements show that, as we increase the dimensionality, the existing modes at lower dimensions are largely maintained, and new modes are added to the ensemble. This indicated an eventual addition of more and more secondary PFMs. Mean±STD of the diagonal values: for the top panel, UK Biobank volumetric fMRI: **25 vs 50**: 0.90 ± 0.11, **50 vs 100**: 0.84 ± 0.14, **100 vs 150**: 0.82 ± 0.14. For the bottom panel, HCP Cortical CIFTI fMRI: **25 vs 50**: 0.88 ± 0.08, **50 vs 100**: 0.90 ± 0.11, **100 vs 150**: 0.88 ± 0.11.

**Figure S 10.**
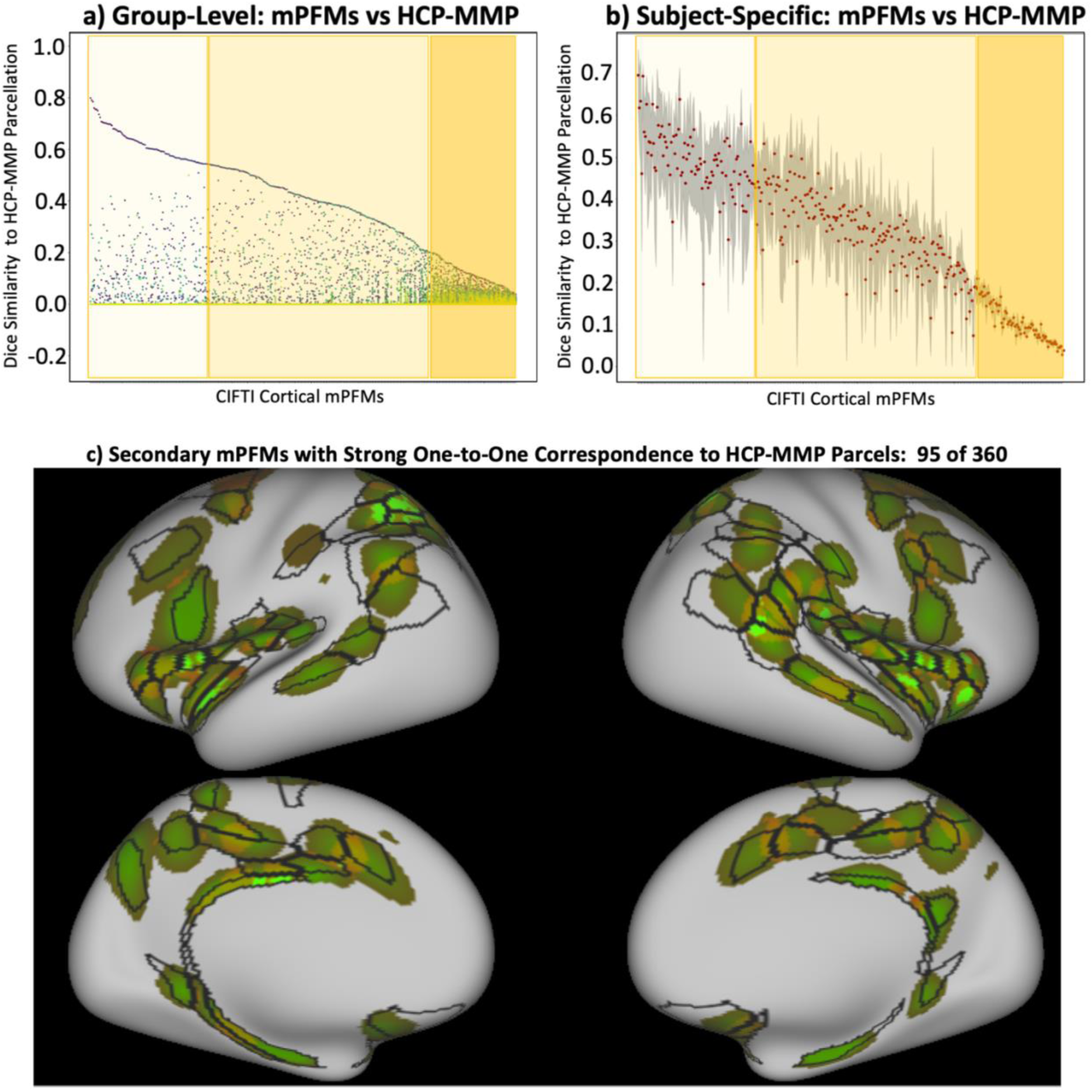
Supplement to section 3.2.3. a) group-level HCP-MMP parcellation with 360 parcels compared to a 360-mPFM decomposition of HCP. 95 modes were parcel-like (lightest shade of yellow) and showed a clear one-to-one match to HCP-MMP parcels. The remaining large-scale and mixed-scale mPFMs are marked with dark and medium shades of yellow, respectively. 73 mPFMs showed less than 0.2 dice similarity to any HCP-MMP parcels (highlighted with the darkest shade of yellow); these included conventional low-dimensional modes and the less well-known variants of large-scale modes, and typically occupied multiple distant sub-regions. b) subject-level HCP MMP (with the same ordering as Fig5a). Median showed in red, and grey error margins show 25 to 75 percentile across subjects. Good pairing for 95 parcel-like modes were observed here as well. c) Top 95 matched pairs of modes between mPFM and HCP-MMP. Black contours denote group-level HCP-MMP parcels and green-brown patches denote thresholded group-level PFMs.

**Figure S 11.**
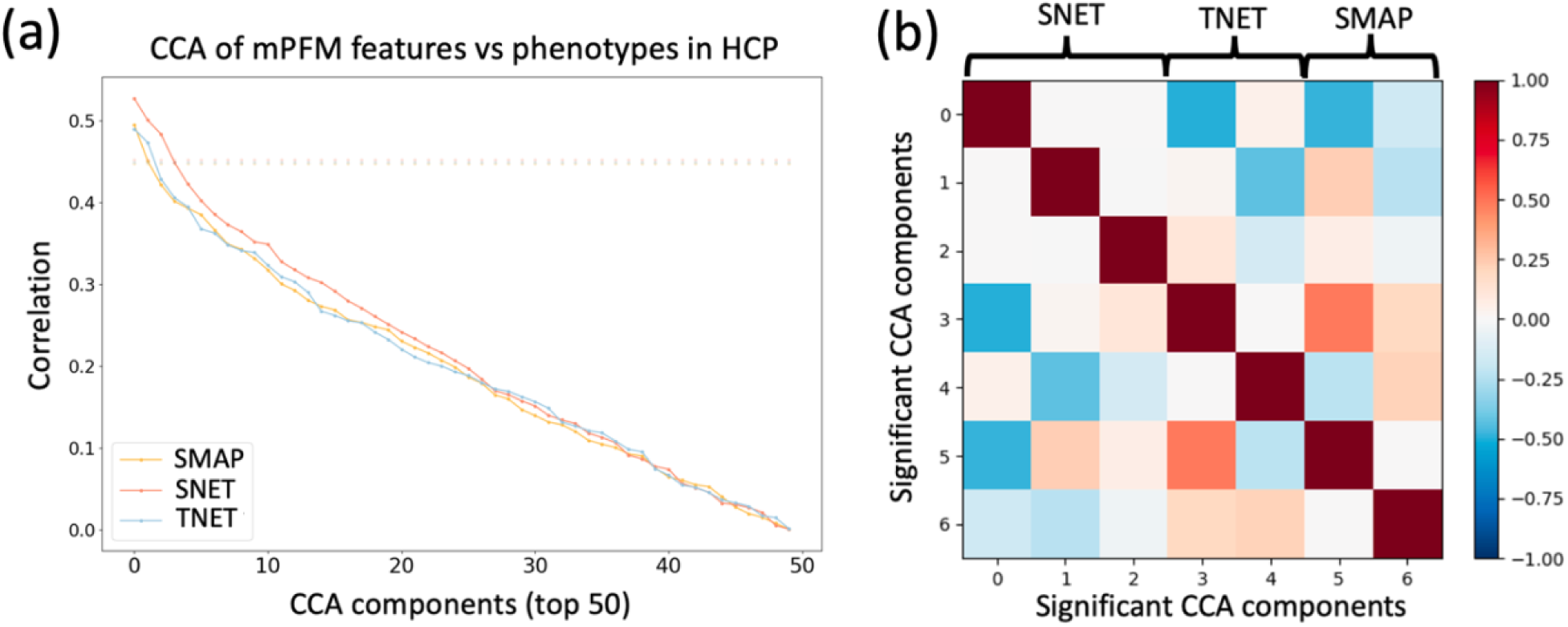
Supplement to Figure 6a shared population covariation between mPFMs and behaviour in HCP. a) Canonical Correlation Analysis (CCA) was used to compute modes of population co-variation between behavioural traits and spatial maps (SMAPs)/Spatial correlations (SNETs)/temporal correlations (TNETs) of mPFMs. Statistical significance of CCA components was determined using multi-level block permutations that takes family structure of HCP data into account (Winkler et al., 2015). Correlation values for the top CCA components of SMAP, SNET and TNET were found to be 0.49, 0.52 and 0.49, respectively. Using the recently-introduced GEMMR tool (Helmer et al., 2024), we verified that given these correlation values, our utilised sample size (1001) is sufficient to yield at least 85% power and at most 10% error for the association strength, weight, scores and loadings; b) 2, 3 and 2 significant CCA components were identified for SMAPs, SNETs and TNETs, respectively, and their correlations are shown. we found maximum correlations between transformed pairs SNET-SMAP, SNET-TNET, and SMAP-TNET to be −0.498, −0.484 and 0.483, respectively, indicating that the phenotype-transformed SMAPs, SNETs, and TNETs capture distinct aspects of subject variability.

**Figure S 12.**
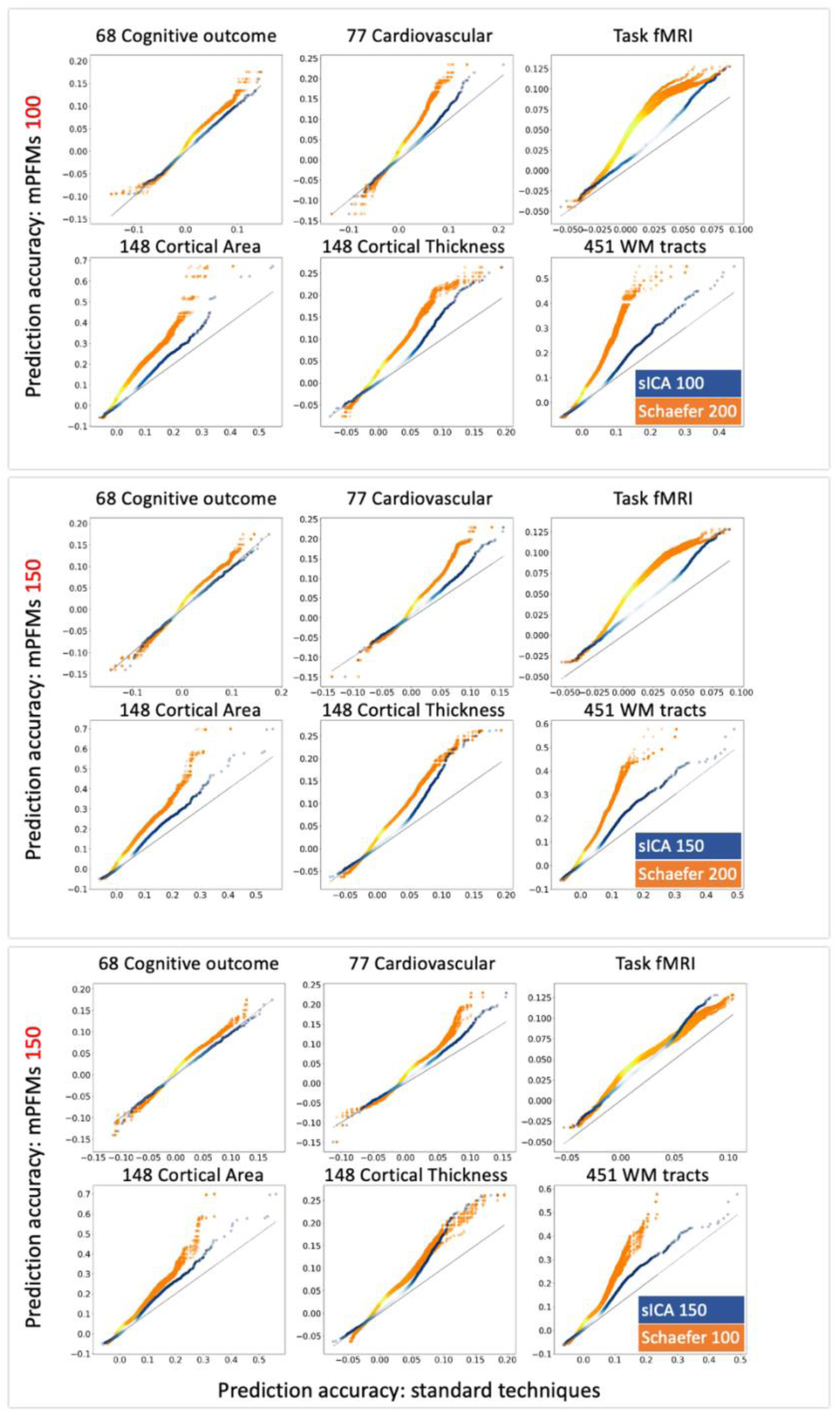
Supplement to Figure 6b replicating trait prediction results of mPFMs vs standard techniques using different choices of dimensionality. Top panel: comparing mPFM 100with spatial ICA (sICA, blue) 100 and Schaefer (orange) 200; middle panel: comparing mPFM 150 with sICA 150 and Schaefer 200; bottom panel: comparing mPFM 150 with sICA 150 and Schaefer 100. In all three scenarios, similar to the original result where d=100 was used for all three methods, mPFMs significantly outperformed the standard techniques in trait predictions. This was especially evident in comparison to Schaefer parcellation. Note that when the number of modes is mismatched between mPFMs/sICA and Schaefer atlas, we have randomly selected a subset of the modes/parcels (e.g., 100 out of 150 in the bottom panel), and repeated that sub-selection 10 times for plotting (hence 10 orange graphs vs 1 blue, given that sICA and mPFM dimensionalities are always matched).

**Table S 1.** A summary of terminology used throughout the paper.

| Terminology | Description |
| --- | --- |
| Functional Modes | Modes of functional brain connectivity, where one or multiple brain regions work in synchrony such that they can be described with a single timecourse. |
| PfMs | Probabilistic Functional Modes: functional modes estimated using sPROFUMO software, which estimates the modes simultaneously for big populations and individuals, thus yielding *personalised* functional modes. |
| RSNs | Resting State Networks: functional modes estimated from resting state fMRI (rfMRI). Resting state refers to brain activity from task-free recordings, where participants are instructed to not think about anything specific while lying in the scanner. |
| Large-scale or Distributed RSNs/PfMs | 25 modes from low-dimensional (lowD) decomposition of rfMRI data using PfMs, unless specified otherwise (e.g., in Figure 2). |
| Multiscale PfMs (mPfMs) | Modes from high-dimensional (highD) decomposition of rfMRI data using PfMs. Depending on dataset/application, we have used different number of modes as highD decomposition. Specifically, for HCP data we have used 50, 100, 150 or 360 modes, for UK Biobank data we have used 150 or 100 modes. Justifications are included the corresponding Methods subsections. |
| Primary mPfMs | 25 distributed modes from high-dimensional PFM decomposition of rfMRI that best-match the large-scale RSNs. |
| Secondary mPfMs | New modes from high-dimensional PFM decomposition that emerge as we increase the dimensionality to $\geq 100$. |
| Spatial Maps, or SMAPs | Spatial organisation of functional modes across the brain voxels. |
| Functional Connectivity | A matrix of size $N_{\text{mode}} \times N_{\text{mode}}$ including partial temporal correlations between timecourses of functional modes, also referred to as TNET. L2/Tikhonov regularisation is used in estimation of partial correlations. |
| Spatial Connectivity | A matrix of size $N_{\text{mode}} \times N_{\text{mode}}$ including spatial correlation coefficients between SMAPs of functional modes, also referred to as SNET. This is an indicator of *spatial overlap* between the modes. |

**Table S 2.** T-value and p-values related to subcomponent analysis: original subcomponents. Statistically significant p-values (Bonferroni-corrected threshold to account for multiple comparisons) are shown in bold font. Supplement to **Figure 3**c&d.

| Winner:<br>Large-Scale |  | Winner:<br>Primary mPFMs |  | Winner:<br>Secondary mPFMs |  |  |  |
| --- | --- | --- | --- | --- | --- | --- | --- |
| t-values | p-values | t-values | p-values | t-values | p-values | t-values | p-values |
| 0.46 | 6.42e-01 | 19.81 | **5.57e-74** | 22.96 | **4.70e-94** | 23.08 | **7.82e-95** |
| 40.45 | **7.04e-213** | 17.29 | **8.47e-59** | 14.19 | **8.30e-42** | 16.34 | **2.07e-53** |
| 1.86 | 6.35e-02 | 22.53 | **2.74e-91** | 17.81 | **7.82e-62** | 29.73 | **8.81e-140** |
| 7.60 | **6.64e-14** | 14.21 | **7.29e-42** | 4.64 | **4.01e-06** | 17.66 | **5.87e-61** |
| 36.36 | **3.06e-185** | 10.26 | **1.54e-23** | 28.59 | **6.15e-132** | -0.61 | 5.43e-01 |
| 37.73 | **1.35e-194** | 5.33 | **1.21e-07** | 31.04 | **9.76e-149** | 24.86 | **1.12e-106** |
| 15.01 | **4.24e-46** | 44.60 | **3.17e-240** | 19.73 | **1.70e-73** | 15.69 | **8.63e-50** |
| -2.64 | 8.33e-03 | 15.12 | **1.07e-46** | 10.46 | **2.31e-24** | 10.89 | **3.61e-26** |
| 7.04 | **3.49e-12** | 14.02 | **6.60e-41** | 24.87 | **1.03e-106** | -0.01 | 9.94e-01 |
| 7.98 | **4.06e-15** | 57.85 | **1.13e-321** | 10.30 | **1.00e-23** | 4.77 | **2.08e-06** |
| 35.48 | **3.05e-179** | 23.84 | **7.72e-100** | 5.45 | **6.42e-08** | 11.08 | **5.24e-27** |
| 15.05 | **2.63e-46** | 14.47 | **3.29e-43** | 12.29 | **1.95e-32** | 40.87 | **1.13e-215** |
|  |  | 4.88 | **1.26e-06** | 4.77 | **2.08e-06** | 32.10 | **5.01e-156** |
|  |  | 16.06 | **8.24e-52** | 16.35 | **1.92e-53** | 18.47 | **8.46e-66** |
|  |  | 7.10 | **2.36e-12** | 51.75 | **2.76e-285** | 26.74 | **2.64e-119** |
|  |  | 37.67 | **3.76e-194** | 9.99 | **1.76e-22** | 26.53 | **7.10e-118** |
|  |  | 10.92 | **2.62e-26** | 27.84 | **8.20e-127** | 16.47 | **4.19e-54** |
|  |  | 3.23 | **1.27e-03** | 52.06 | **3.32e-287** | 35.81 | **1.68e-181** |
|  |  | 15.55 | **4.96e-49** | 17.32 | **5.24e-59** | 39.36 | **1.46e-205** |
|  |  | 17.40 | **1.91e-59** | 44.44 | **3.60e-239** | 61.30 | **0.00e+00** |
|  |  | 6.98 | **5.22e-12** | 10.31 | **9.63e-24** | 29.06 | **3.88e-135** |
|  |  | 33.82 | **7.52e-168** | 31.74 | **1.39e-153** | 17.77 | **1.28e-61** |
|  |  | 37.02 | **9.35e-190** | 24.91 | **5.39e-107** | 29.35 | **3.66e-137** |
|  |  | 26.00 | **2.47e-114** | 15.64 | **1.69e-49** | 11.63 | **1.99e-29** |
|  |  | 27.83 | **9.80e-127** | 24.38 | **1.78e-103** | 13.61 | **7.77e-39** |
|  |  | 16.07 | **6.90e-52** | 21.47 | **2.08e-84** | 34.87 | **4.61e-175** |
|  |  | 19.13 | **9.34e-70** | 22.20 | **3.81e-89** | 9.84 | **7.32e-22** |
|  |  | 38.03 | **1.39e-196** | 38.65 | **8.15e-201** | 54.03 | **3.83e-299** |
|  |  |  |  | 61.69 | **0.00e+00** | 23.29 | **2.93e-96** |
|  |  |  |  | 27.37 | **1.24e-123** | 25.14 | **1.54e-108** |
|  |  |  |  | 27.32 | **2.92e-123** | 14.85 | **2.95e-45** |
|  |  |  |  | 49.52 | **1.59e-271** | 74.34 | **0.00e+00** |
|  |  |  |  | 88.06 | **0.00e+00** | 6.00 | **2.82e-09** |
|  |  |  |  | 44.50 | **1.58e-239** | 14.35 | **1.34e-42** |
|  |  |  |  | 3.59 | **3.42e-04** | 19.80 | **6.30e-74** |
|  |  |  |  | 11.52 | **6.05e-29** | 24.71 | **1.17e-105** |
|  |  |  |  | 9.78 | **1.22e-21** | 19.73 | **1.75e-73** |
|  |  |  |  | 22.43 | **1.26e-90** | 13.38 | **1.12e-37** |
|  |  |  |  | 11.77 | **4.55e-30** | 13.04 | **5.38e-36** |
|  |  |  |  | 40.02 | **5.40e-210** | 62.63 | **0.00e+00** |
|  |  |  |  | 34.89 | **3.22e-175** | 16.04 | **1.02e-51** |
|  |  |  |  | 14.68 | **2.54e-44** | 37.97 | **3.27e-196** |
|  |  |  |  |  |  | 49.84 | **1.71e-273** |

**Table S 3.** T-value and p-values related to subcomponent analysis: temporally-exclusive subcomponents. Statistically significant p-values (Bonferroni-corrected threshold to account for multiple comparisons) are shown in bold font. Supplement to **Figure 3**e&f.

| Winner:<br>Large-Scale |  |  |  | Winner:<br>Primary mPFMs |  | Winner:<br>Secondary mPFMs |  |
| --- | --- | --- | --- | --- | --- | --- | --- |
| t-values | p-values | t-values | p-values | t-values | p-values | t-values | p-values |
| 30.98 | **2.52e-148** | 46.74 | **6.02e-254** | 1.52 | 1.28e-01 | 6.41 | **2.20e-10** |
| 28.70 | **1.00e-132** | 37.99 | **2.40e-196** | 3.12 | **1.88e-03** | 15.88 | **8.34e-51** |
| 15.48 | **1.20e-48** | 4.87 | **1.31e-06** | 4.21 | **2.78e-05** | 1.55 | 1.22e-01 |
| 4.11 | **4.30e-05** | 34.01 | **3.38e-169** | 1.13 | 2.59e-01 | 6.07 | **1.82e-09** |
| 41.43 | **2.13e-219** | 20.80 | **3.71e-80** |  |  | 11.45 | **1.27e-28** |
| 1.62 | 1.06e-01 | 35.79 | **2.27e-181** |  |  | 11.26 | **8.68e-28** |
| 15.50 | **9.33e-49** | 10.34 | **7.19e-24** |  |  | 1.69 | 9.20e-02 |
| 10.65 | **3.63e-25** | 10.02 | **1.32e-22** |  |  |  |  |
| 67.89 | **0.00e+00** | 1.27 | 2.04e-01 |  |  |  |  |
| 16.01 | **1.50e-51** | 62.64 | **0.00e+00** |  |  |  |  |
| 26.71 | **4.26e-119** | 37.78 | **6.91e-195** |  |  |  |  |
| 7.64 | **5.18e-14** | 37.84 | **2.51e-195** |  |  |  |  |
| 42.21 | **1.46e-224** | 25.98 | **3.70e-114** |  |  |  |  |
| 15.03 | **3.57e-46** | 34.11 | **7.63e-170** |  |  |  |  |
| 24.92 | **4.92e-107** | 42.65 | **1.83e-227** |  |  |  |  |
| 6.33 | **3.61e-10** | 10.44 | **2.72e-24** |  |  |  |  |
| 22.85 | **2.21e-93** | 22.93 | **6.88e-94** |  |  |  |  |
| 60.43 | **0.00e+00** | 58.42 | **0.00e+00** |  |  |  |  |
| 25.70 | **2.91e-112** | 25.22 | **4.90e-109** |  |  |  |  |
| 5.16 | **3.02e-07** | 26.83 | **6.54e-120** |  |  |  |  |
| 27.78 | **2.27e-126** | 22.02 | **5.46e-88** |  |  |  |  |
| 8.02 | **3.00e-15** | 8.93 | **1.96e-18** |  |  |  |  |
| 50.27 | **3.33e-276** | 54.66 | **6.28e-303** |  |  |  |  |
| 33.88 | **2.63e-168** | 3.63 | **2.99e-04** |  |  |  |  |
| 27.60 | **3.55e-125** | 3.79 | **1.58e-04** |  |  |  |  |
| 17.19 | **3.05e-58** | 2.38 | 1.75e-02 |  |  |  |  |
| 13.68 | **3.60e-39** | 72.20 | **0.00e+00** |  |  |  |  |
| 71.00 | **0.00e+00** | 44.52 | **1.04e-239** |  |  |  |  |
| 43.53 | **3.04e-233** | 21.80 | **1.47e-86** |  |  |  |  |
| 34.39 | **9.00e-172** | 23.88 | **3.76e-100** |  |  |  |  |
| 30.32 | **8.00e-144** | 4.15 | **3.59e-05** |  |  |  |  |
| 35.19 | **2.98e-177** | 58.48 | **0.00e+00** |  |  |  |  |
| 38.02 | **1.57e-196** | 24.23 | **1.77e-102** |  |  |  |  |
| 33.46 | **2.09e-165** | 8.54 | **4.82e-17** |  |  |  |  |
| 28.81 | **1.82e-133** | 105.56 | **0.00e+00** |  |  |  |  |
| 23.01 | **2.10e-94** | 29.69 | **1.62e-139** |  |  |  |  |
| 21.48 | **1.81e-84** | 23.61 | **2.41e-98** |  |  |  |  |
| 49.70 | **1.10e-272** | 34.32 | **2.69e-171** |  |  |  |  |
| 30.70 | **2.12e-146** | 27.49 | **2.07e-124** |  |  |  |  |
| 20.28 | **6.98e-77** | 69.98 | **0.00e+00** |  |  |  |  |
| 16.30 | **3.74e-53** | 10.83 | **6.31e-26** |  |  |  |  |
| 24.46 | **5.31e-104** | 9.19 | **2.15e-19** |  |  |  |  |
| 63.10 | **0.00e+00** | 14.30 | **2.33e-42** |  |  |  |  |
| 3.10 | 1.97e-03 | 4.85 | **1.40e-06** |  |  |  |  |
| 12.88 | **3.22e-35** | 38.13 | **2.64e-197** |  |  |  |  |
| 18.38 | **3.02e-65** | 23.52 | **1.00e-97** |  |  |  |  |
| 12.50 | **2.12e-33** | 11.53 | **5.92e-29** |  |  |  |  |
| 78.92 | **0.00e+00** | 74.21 | **0.00e+00** |  |  |  |  |
| 35.71 | **8.46e-181** | 18.10 | **1.39e-63** |  |  |  |  |
| 19.95 | **7.70e-75** | 42.58 | **5.12e-227** |  |  |  |  |
| 30.69 | **2.35e-146** | 31.93 | **6.42e-155** |  |  |  |  |
| 25.75 | **1.29e-112** | 38.12 | **3.17e-197** |  |  |  |  |
| 19.17 | **4.85e-70** | 91.42 | **0.00e+00** |  |  |  |  |
| 38.45 | **1.86e-199** | 29.99 | **1.49e-141** |  |  |  |  |
| 21.55 | **5.77e-85** | 30.16 | **9.74e-143** |  |  |  |  |
| 23.77 | **2.17e-99** | 27.38 | **1.14e-123** |  |  |  |  |
| 2.54 | 1.12e-02 | 22.64 | **5.45e-92** |  |  |  |  |

**Table S 4.**
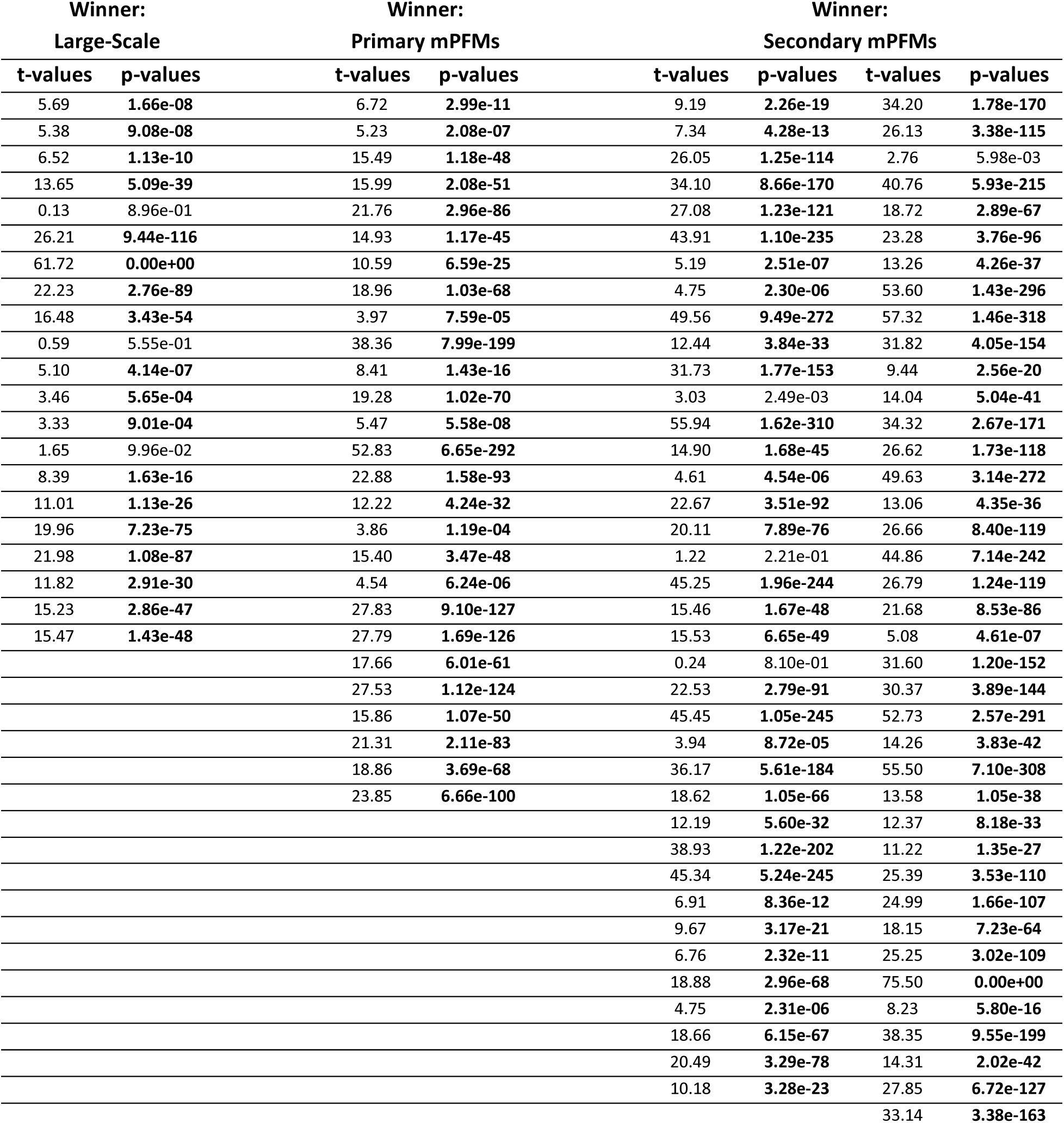
T-value and p-values related to subcomponent analysis: spatially-exclusive subcomponents. Statistically significant p-values (Bonferroni-corrected threshold to account for multiple comparisons) are shown in bold font. Supplement to **Figure 3**g&h.

**Table S 5.** T-value and Bonferroni-corrected (36 comparisons) p-values to compare phenotype-prediction performance of Spatial and Temporal/Functional connectivity features related to: a) LowD large-scale RSNs vs highD mPFMs and b) within-scale vs cross-scale mPFMs. Pri: Primary, Sec: Secondary, Statistically significant values are shown in bold font. Supplement to **Figure 4**.

| T-Value<br>(corrected P-Value) |  |  |  |  |  |  |
| --- | --- | --- | --- | --- | --- | --- |
|  | Spatial connectivity |  |  | Temporal connectivity |  |  |
|  | lowD > mPFMs | Pri2Pri > Pri2Sec | Sec2Sec > Pri2Sec | lowD > mPFMs | Pri2Pri > Pri2Sec | Sec2Sec > Pri2Sec |
| Cognitive | -1.464<br>(0.148) | -1.243<br>(0.219) | **-3.526**<br>**(7.92E-04)** | -0.6182<br>(0.539) | -1.5678<br>(0.122) | -0.487<br>(0.628) |
| GMA | **-21.657**<br>**(1.37E-47)** | **-21.017**<br>**(3.89E-46)** | **-14.978**<br>**(2.21E-31)** | **-4.6115**<br>**(8.88E-06)** | **-2.6633**<br>**(8.64E-03)** | **3.3587**<br>**(1.01E-03)** |
| GMt | **-9.226**<br>**(2.81E-16)** | **-12.042**<br>**(1.14E-23)** | **-4.916**<br>**(2.34E-06)** | -1.0236<br>(0.31) | -0.1672<br>(0.867) | **2.0203**<br>**(0.045)** |
| WM | **-26.469**<br>**(6.31E-94)** | **-25.468**<br>**(2.19E-89)** | -0.583<br>(0.56) | 1.0486<br>(0.295) | **-9.8976**<br>**(5.09E-21)** | **7.1892**<br>**(2.75E-12)** |
| Task-Contrast | **-15.190**<br>**(4.15E-35)** | **-7.165**<br>**(1.48E-11)** | **-56.461**<br>**(1.94E-124)** | **-7.0706**<br>**(2.56E-11)** | -0.6513<br>(0.516) | **35.8817**<br>**(7.73E-89)** |
| Cardiovascular | **-7.272**<br>**(2.68E-10)** | **-4.296**<br>**(5.09E-05)** | 0.663<br>(0.51) | 0.2176<br>(0.828) | **-5.4334**<br>**(6.59E-07)** | -0.5626<br>(0.575) |

**Table S 6.** T-values and Bonferroni-corrected (6 comparisons) p-values to compare phenotype-prediction performance of mPFMs vs spatial ICA (sICA) and Schaefer Parcellation of the same dimensionality (100 modes). Statistically significant values have been shown in bold font. Supplement to **Figure 6**.

|  | T-value |  |  | Corrected P-Value |  |  |
| --- | --- | --- | --- | --- | --- | --- |
|  | mPFM > sICA | mPFM > Schaefer | sICA > Schaefer | mPFM vs sICA | mPFM vs Schaefer | sICA vs Schaefer |
| Cognitive | 1.261 | **7.385** | **7.252** | 0.212 | **3.61E-10** | **6.21E-10** |
| GMA | **15.137** | **26.991** | **23.107** | **8.57E-32** | **7.90E-59** | **8.59E-51** |
| GMt | **13.964** | **33.923** | **31.499** | **9.64E-29** | **2.09E-71** | **3.06E-67** |
| WM | **12.675** | **51.843** | **57.347** | **1.07E-31** | **6.96E-192** | **5.19E-209** |
| Task-Contrast | **14.572** | **14.496** | **5.086** | **1.93E-19** | **2.38E-19** | **5.75E-06** |
| Cardiovascular | **7.974** | **17.409** | **17.33** | **1.24E-11** | **3.07E-28** | **4.04E-28** |

**Table S 7.**
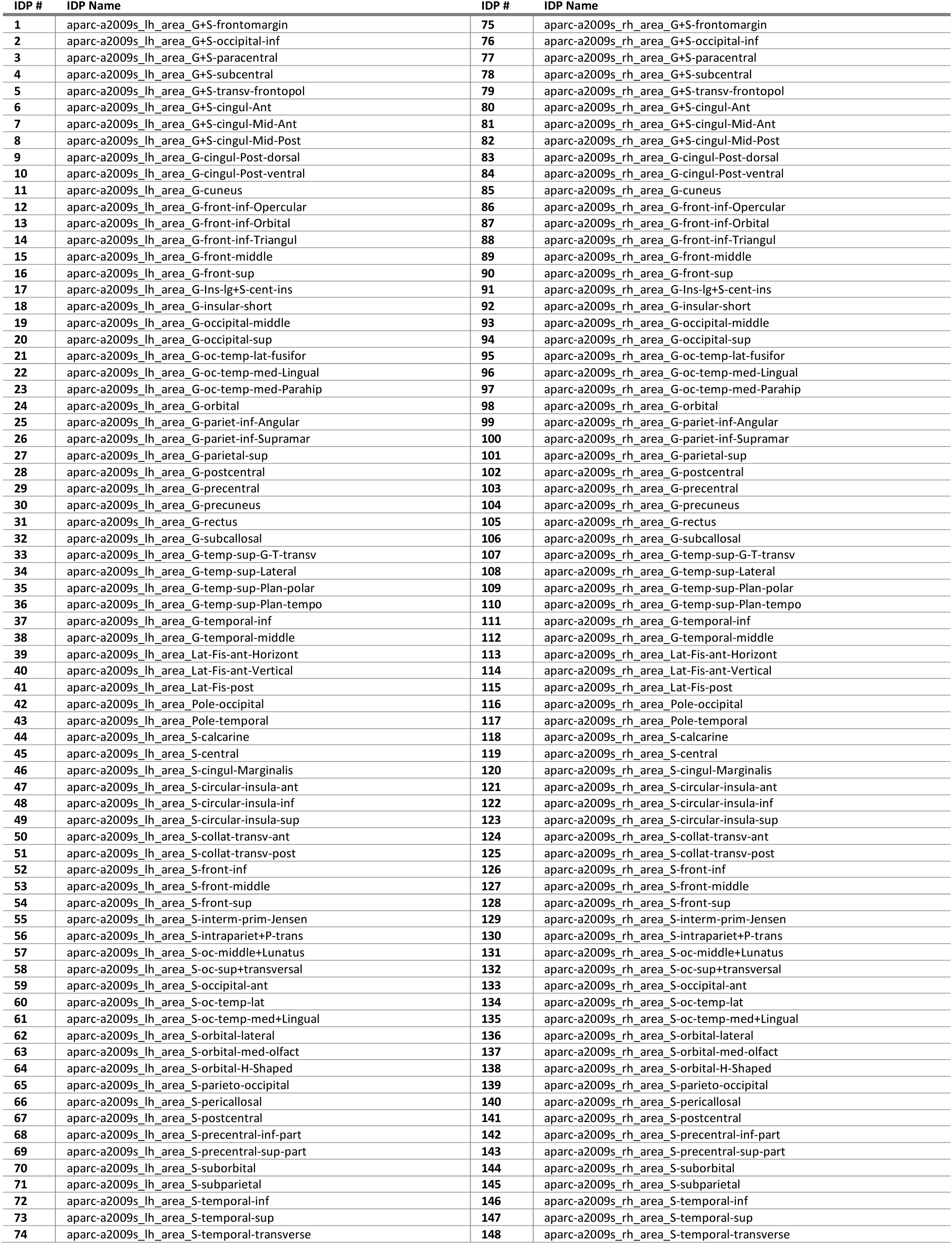
Names of UK Biobank Imaging Derived Phenotypes (IDPs) related to GM area: 148. Supplement to Figure 6.

**Table S 8.**
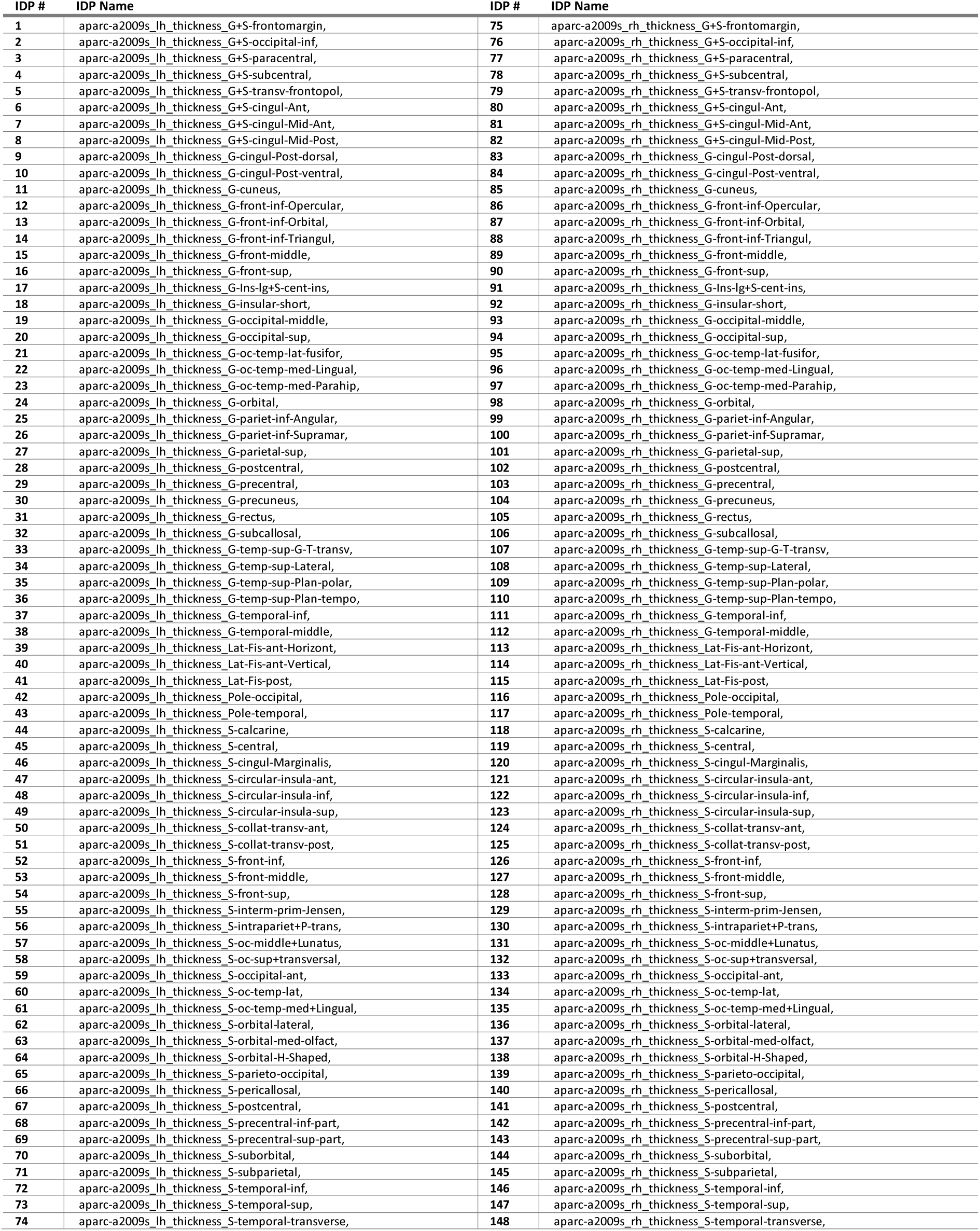
Names of UK Biobank Imaging Derived Phenotypes (IDPs) related to Grey Matter thickness: 148. Supplement to Figure 6.

**Table S 9.**
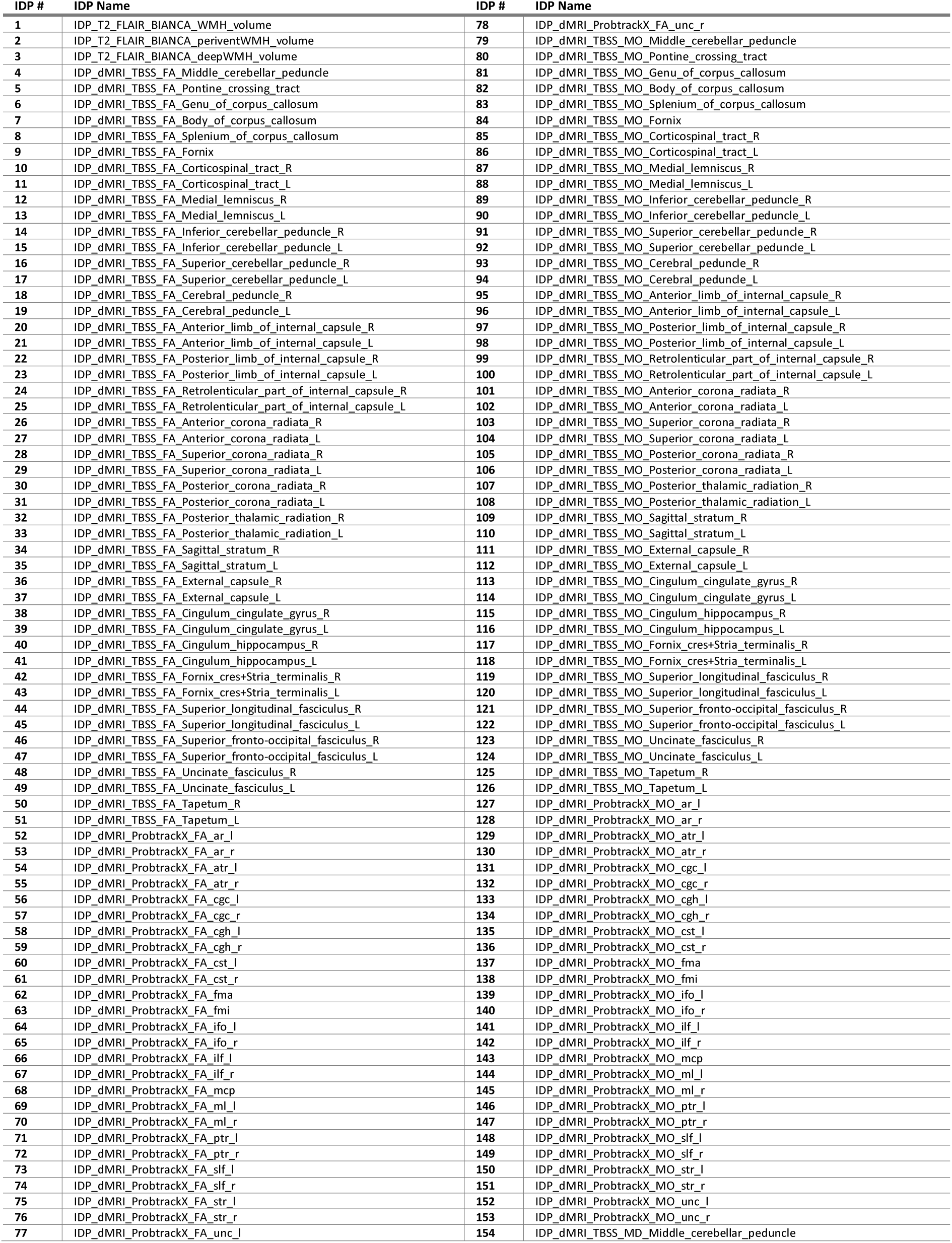
Names of UK Biobank Imaging Derived Phenotypes (IDPs) related to White Matter: 453. Supplement to Figure 6.

**Table S 10.**
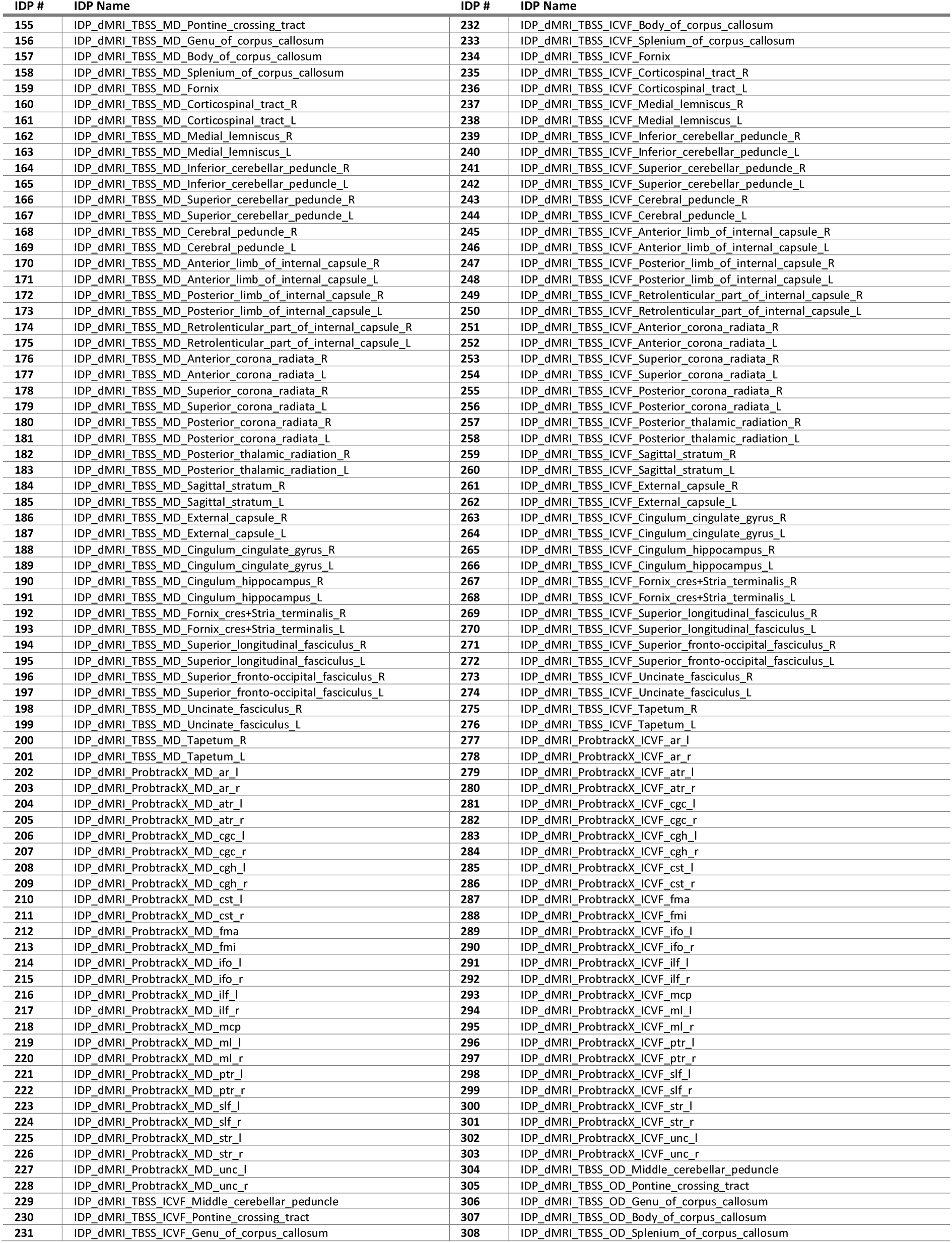
Names of UK Biobank Imaging Derived Phenotypes (IDPs) related to White Matter-continued: 453. Supplement to Figure 6.

**Table S 11.**
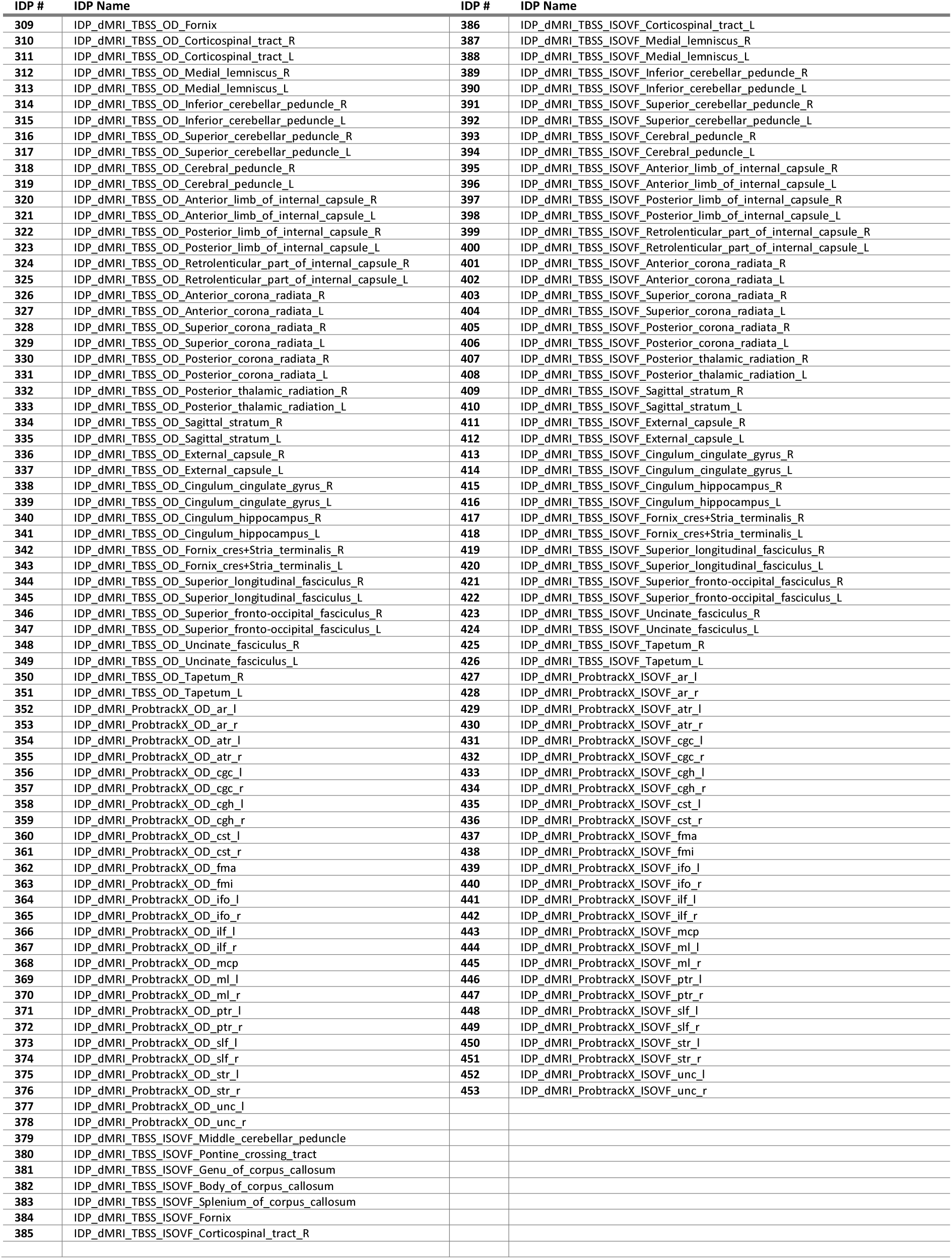
Names of UK Biobank Imaging Derived Phenotypes (IDPs) related to White Matter-continued: 453. Supplement to Figure 6.

**Table S 12.**
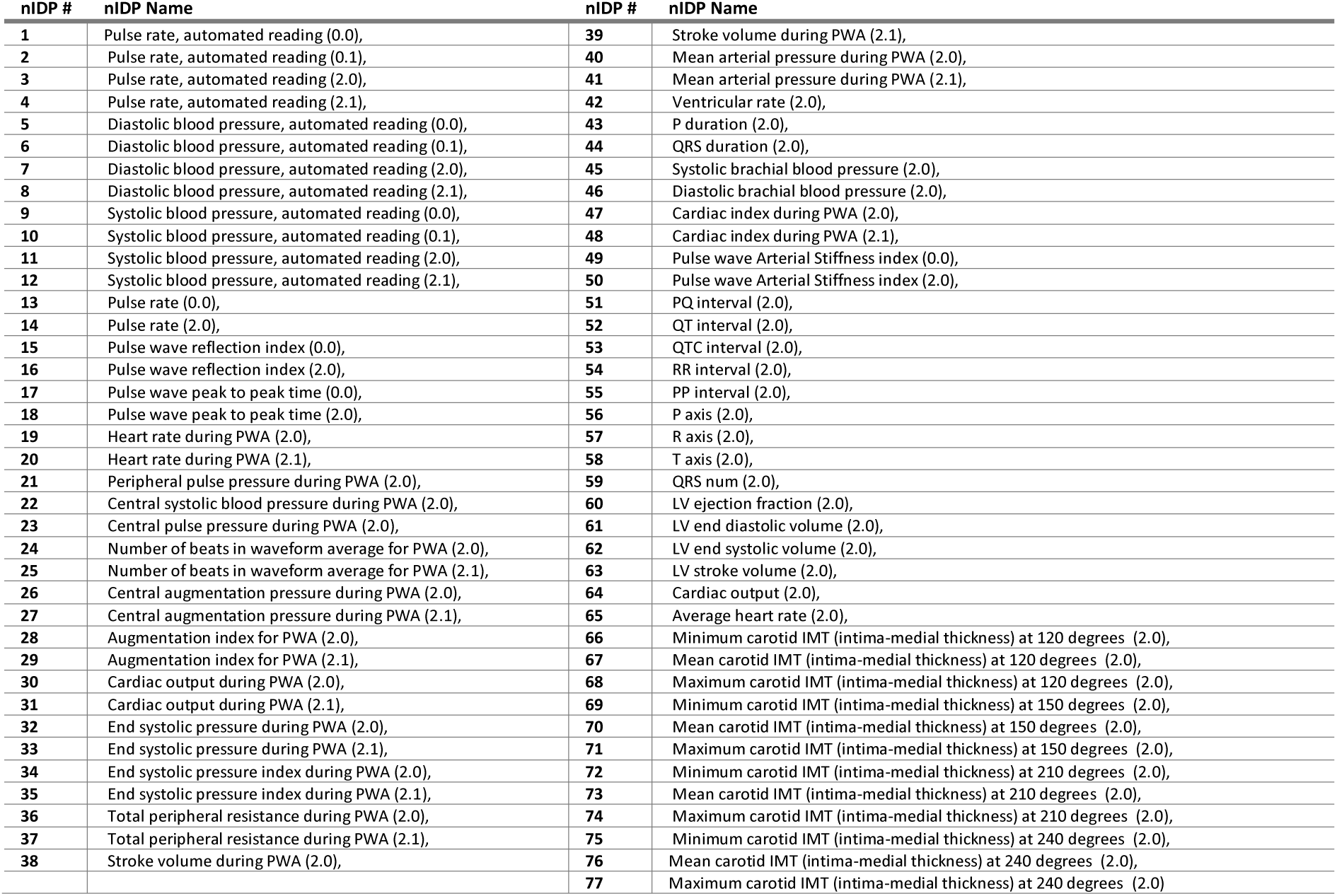
Names of UK Biobank Non-Imaging Derived Phenotypes (nIDPs) related to Cardiovascular Health: 77. Supplement to Figure 6.

| nIDP # | nIDP Name | nIDP # | nIDP Name |
| --- | --- | --- | --- |
| 1 | Pulse rate, automated reading (0.0), | 39 | Stroke volume during PWA (2.1), |
| 2 | Pulse rate, automated reading (0.1), | 40 | Mean arterial pressure during PWA (2.0), |
| 3 | Pulse rate, automated reading (2.0), | 41 | Mean arterial pressure during PWA (2.1), |
| 4 | Pulse rate, automated reading (2.1), | 42 | Ventricular rate (2.0), |
| 5 | Diastolic blood pressure, automated reading (0.0), | 43 | P duration (2.0), |
| 6 | Diastolic blood pressure, automated reading (0.1), | 44 | QRS duration (2.0), |
| 7 | Diastolic blood pressure, automated reading (2.0), | 45 | Systolic brachial blood pressure (2.0), |
| 8 | Diastolic blood pressure, automated reading (2.1), | 46 | Diastolic brachial blood pressure (2.0), |
| 9 | Systolic blood pressure, automated reading (0.0), | 47 | Cardiac index during PWA (2.0), |
| 10 | Systolic blood pressure, automated reading (0.1), | 48 | Cardiac index during PWA (2.1), |
| 11 | Systolic blood pressure, automated reading (2.0), | 49 | Pulse wave Arterial Stiffness index (0.0), |
| 12 | Systolic blood pressure, automated reading (2.1), | 50 | Pulse wave Arterial Stiffness index (2.0), |
| 13 | Pulse rate (0.0), | 51 | PQ interval (2.0), |
| 14 | Pulse rate (2.0), | 52 | QT interval (2.0), |
| 15 | Pulse wave reflection index (0.0), | 53 | QTC interval (2.0), |
| 16 | Pulse wave reflection index (2.0), | 54 | RR interval (2.0), |
| 17 | Pulse wave peak to peak time (0.0), | 55 | PP interval (2.0), |
| 18 | Pulse wave peak to peak time (2.0), | 56 | P axis (2.0), |
| 19 | Heart rate during PWA (2.0), | 57 | R axis (2.0), |
| 20 | Heart rate during PWA (2.1), | 58 | T axis (2.0), |
| 21 | Peripheral pulse pressure during PWA (2.0), | 59 | QRS num (2.0), |
| 22 | Central systolic blood pressure during PWA (2.0), | 60 | LV ejection fraction (2.0), |
| 23 | Central pulse pressure during PWA (2.0), | 61 | LV end diastolic volume (2.0), |
| 24 | Number of beats in waveform average for PWA (2.0), | 62 | LV end systolic volume (2.0), |
| 25 | Number of beats in waveform average for PWA (2.1), | 63 | LV stroke volume (2.0), |
| 26 | Central augmentation pressure during PWA (2.0), | 64 | Cardiac output (2.0), |
| 27 | Central augmentation pressure during PWA (2.1), | 65 | Average heart rate (2.0), |
| 28 | Augmentation index for PWA (2.0), | 66 | Minimum carotid IMT (intima-medial thickness) at 120 degrees (2.0), |
| 29 | Augmentation index for PWA (2.1), | 67 | Mean carotid IMT (intima-medial thickness) at 120 degrees (2.0), |
| 30 | Cardiac output during PWA (2.0), | 68 | Maximum carotid IMT (intima-medial thickness) at 120 degrees (2.0), |
| 31 | Cardiac output during PWA (2.1), | 69 | Minimum carotid IMT (intima-medial thickness) at 150 degrees (2.0), |
| 32 | End systolic pressure during PWA (2.0), | 70 | Mean carotid IMT (intima-medial thickness) at 150 degrees (2.0), |
| 33 | End systolic pressure during PWA (2.1), | 71 | Maximum carotid IMT (intima-medial thickness) at 150 degrees (2.0), |
| 34 | End systolic pressure index during PWA (2.0), | 72 | Minimum carotid IMT (intima-medial thickness) at 210 degrees (2.0), |
| 35 | End systolic pressure index during PWA (2.1), | 73 | Mean carotid IMT (intima-medial thickness) at 210 degrees (2.0), |
| 36 | Total peripheral resistance during PWA (2.0), | 74 | Maximum carotid IMT (intima-medial thickness) at 210 degrees (2.0), |
| 37 | Total peripheral resistance during PWA (2.1), | 75 | Minimum carotid IMT (intima-medial thickness) at 240 degrees (2.0), |
| 38 | Stroke volume during PWA (2.0), | 76 | Mean carotid IMT (intima-medial thickness) at 240 degrees (2.0), |
|  |  | 77 | Maximum carotid IMT (intima-medial thickness) at 240 degrees (2.0) |

**Table S 13.** Names of UK Biobank Non-Imaging Derived Phenotypes (nIDPs) related to Cognition: 68. Supplement to Figure 6.

| nIDP # | nIDP Name |
| --- | --- |
| 1 | Number of incorrect matches in round (0.1) |
| 2 | Number of incorrect matches in round (0.2) |
| 3 | Number of incorrect matches in round (2.1) |
| 4 | Number of incorrect matches in round (2.2) |
| 5 | Number of incorrect matches in round (2.3) |
| 6 | Number of times snap-button pressed (0.0) |
| 7 | Number of times snap-button pressed (0.1) |
| 8 | Number of times snap-button pressed (0.2) |
| 9 | Number of times snap-button pressed (0.3) |
| 10 | Number of times snap-button pressed (0.4) |
| 11 | Number of times snap-button pressed (0.10) |
| 12 | Number of times snap-button pressed (0.11) |
| 13 | Number of times snap-button pressed (2.0) |
| 14 | Number of times snap-button pressed (2.1) |
| 15 | Number of times snap-button pressed (2.2) |
| 16 | Number of times snap-button pressed (2.3) |
| 17 | Number of times snap-button pressed (2.4) |
| 18 | Number of times snap-button pressed (2.5) |
| 19 | Number of times snap-button pressed (2.7) |
| 20 | Number of times snap-button pressed (2.10) |
| 21 | Number of times snap-button pressed (2.11) |
| 22 | Prospective memory result (0.0) |
| 23 | Prospective memory result (2.0) |
| 24 | FI8 : chained arithmetic (2.0) |
| 25 | FI9 : concept interpolation (2.0) |
| 26 | Duration to complete numeric path (trail #1) (2.0) |
| 27 | Mean time to correctly identify matches (2.0) |
| 28 | Number of incorrect matches in round (0.0) |
| 29 | Time elapsed (2.0) |
| 30 | Time elapsed (2.1) |
| 31 | Time elapsed (2.2) |
| 32 | Time elapsed (2.3) |
| 33 | Time elapsed (2.4) |
| 34 | Time elapsed (2.5) |
| 35 | Time elapsed (2.6) |
| 36 | Time elapsed (2.7) |
| 37 | Time elapsed (2.8) |
| 38 | Digits entered correctly (2.0) |
| 39 | Digits entered correctly (2.1) |
| 40 | Digits entered correctly (2.2) |
| 41 | Digits entered correctly (2.3) |
| 42 | Digits entered correctly (2.4) |
| 43 | Digits entered correctly (2.5) |
| 44 | Digits entered correctly (2.6) |
| 45 | Digits entered correctly (2.7) |
| 46 | Total errors traversing numeric path (trail #1) (2.0) |
| 47 | Mean time to correctly identify matches (0.0) |
| 48 | Fluid intelligence score (0.0) |
| 49 | Fluid intelligence score (2.0) |
| 50 | Duration to complete numeric path (trail #1) (0.0) |
| 51 | Duration to complete alphanumeric path (trail #2) (0.0) |
| 52 | Digits entered correctly (2.8) |
| 53 | Maximum digits remembered correctly (2.0) |
| 54 | Attempted fluid intelligence (FI) test. (2.0) |
| 55 | FI1 : numeric addition test (0.0) |
| 56 | FI1 : numeric addition test (2.0) |
| 57 | FI3 : word interpolation (0.0) |
| 58 | FI3 : word interpolation (2.0) |
| 59 | FI4 : positional arithmetic (0.0) |
| 60 | FI4 : positional arithmetic (2.0) |
| 61 | FI5 : family relationship calculation (0.0) |
| 62 | FI5 : family relationship calculation (2.0) |
| 63 | FI6 : conditional arithmetic (0.0) |
| 64 | FI6 : conditional arithmetic (2.0) |
| 65 | FI7 : synonym (0.0) |
| 66 | FI7 : synonym (2.0) |
| 67 | Duration to complete alphanumeric path (trail #2) (2.0) |
| 68 | Number of puzzles correctly solved (2.0) |

**Table S 14.**
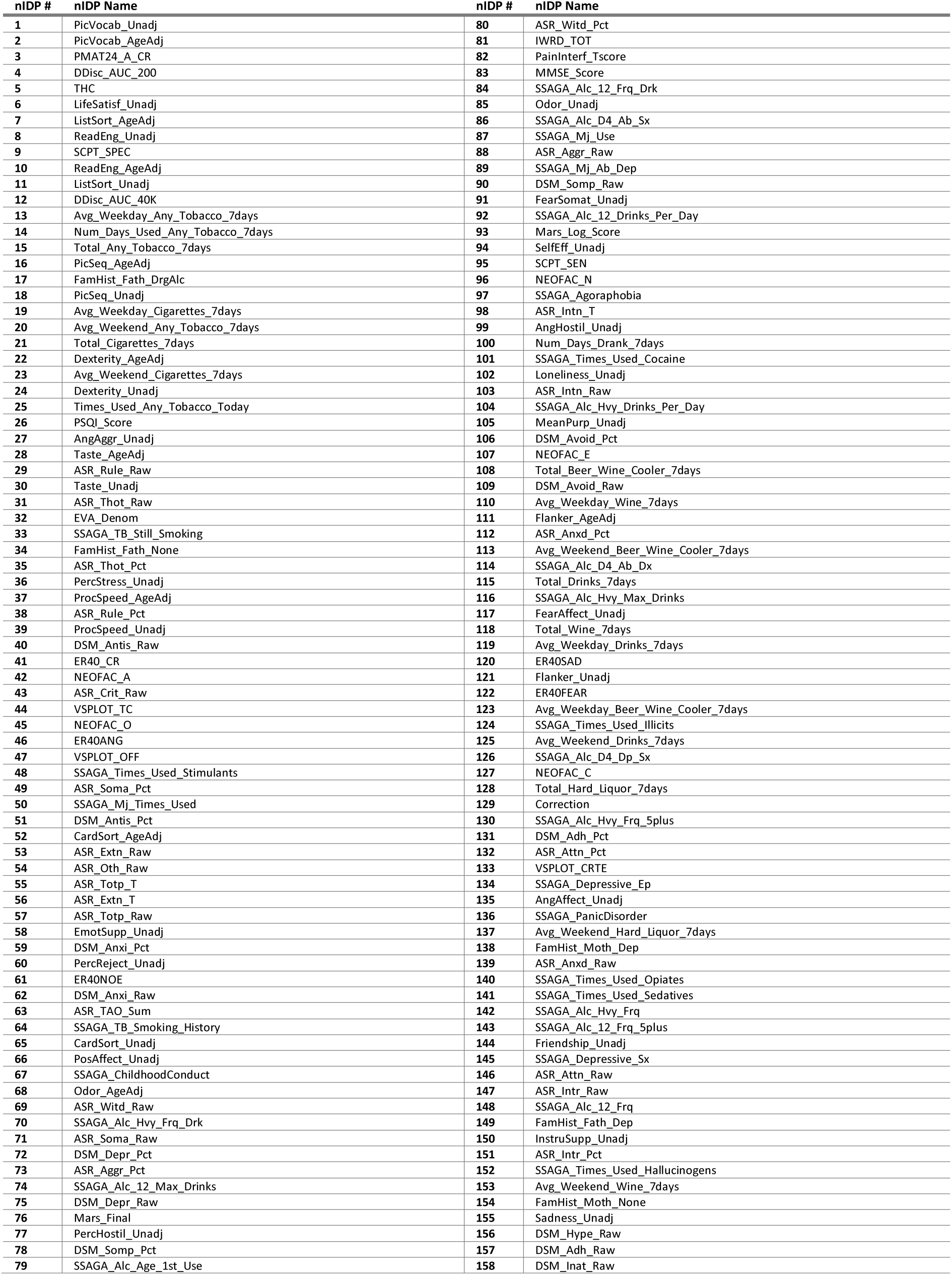
Names of Human Connectome Project Non-Imaging Derived Phenotypes (nIDPs) used in CCA. Supplement to Figure 6.

## Notes

### Competing Interest Statement

The authors have declared no competing interest.

### Summary of Updates

The manuscript has been reformatted to match the target journal requirements, and minor adjustments were made in that process. Additionally, during the peer review process, we revised several subsections and added new figures. In particular: New figure 2 and relevant results section; new supplementary figures S1, S9, and S12, as well as their corresponding Methods and Results revisions. Sections 2.2.1, 2.3, Results paragraph 1, sections 3.1, 3.2.2.1, 3.2.3, 3.3, and Discussion were revised.

